# Development of the first low nanomolar Liver Receptor Homolog-1 agonist through structure-guided design

**DOI:** 10.1101/639732

**Authors:** Suzanne G. Mays, Autumn R. Flynn, Jeffery L. Cornelison, C. Denise Okafor, Hongtao Wang, Guohui Wang, Xiangsheng Huang, Heather N. Donaldson, Elizabeth J. Millings, Rohini Polavarapu, David D. Moore, John W. Calvert, Nathan T. Jui, Eric A. Ortlund

## Abstract

As a key regulator of metabolism and inflammation, the orphan nuclear hormone receptor, Liver Receptor Homolog-1 (LRH-1), has potential as a therapeutic target for diabetes, nonalcoholic fatty liver disease, and inflammatory bowel diseases. Discovery of LRH-1 modulators has been difficult, in part due to the tendency for synthetic compounds to bind unpredictably within the lipophilic binding pocket. Using a structure-guided approach, we exploited a newly-discovered polar interaction to lock agonists in a consistent orientation. This enabled the discovery of the first low nanomolar LRH-1 agonist, one hundred times more potent than the best previous modulator. We elucidate a novel mechanism of action that relies upon specific polar interactions deep in the LRH-1 binding pocket. In an organoid model of inflammatory bowel disease, the new agonist increases expression of LRH-1-conrolled steroidogenic genes and promotes anti-inflammatory gene expression changes. These studies constitute major progress in developing LRH-1 modulators with potential clinical utility.

## INTRODUCTION

Liver Receptor Homolog-1 (LRH-1; NR5A2) is a nuclear hormone receptor (NR) that is highly expressed in the liver and tissues of endodermal origin. It is indispensable during embryonic development, where it plays a role in maintenance of pluripotency^1^, as well as in the development of the liver and pancreas^2^. In adults, LRH-1 controls diverse transcriptional programs in different tissues related to metabolism, inflammation, and cellular proliferation. Targeted metabolic pathways include bile acid biosynthesis^3^, reverse cholesterol transport^4–5^, *de novo* lipogenesis^6–7^, and glucose phosphorylation and transport^8–9^. The ability to modulate lipid and glucose metabolism suggests therapeutic potential for LRH-1 agonists in metabolic diseases such as nonalcoholic fatty liver disease, type II diabetes, and cardiovascular disease. Indeed, the phospholipid LRH-1 agonist dilauroylphosphatidylcholine (DLPC; PC 12:0/12:0) improves glucose tolerance, insulin sensitivity, and triglyceride levels in obese mice^6^. These anti-diabetic effects occur in an LRH-1-dependent manner and have been primarily attributed to a reduction of *de novo* lipogenesis^6^. In addition, targeting LRH-1 in the gut has therapeutic potential for inflammatory bowel disease, where LRH-1 overexpression ameliorates disease-associated inflammation and cell death^10^.

While small molecule LRH-1 modulators are highly sought, the large and lipophilic LRH-1 binding pocket has been extremely challenging to target. A promising class of agonists developed by Whitby and colleagues features a bicyclic hexahydropentalene core scaffold^11–12^. The best-studied of this class, named RJW100, was discovered as a part of an extensive synthetic effort to improve acid stability and efficacy of a related compound, GSK8470^12^ (Figure 1A). We recently determined the crystal structure of LRH-1 bound to RJW100 and made a surprising discovery: it exhibits a completely different binding mode than GSK8470, such that the bicyclic cores of the two agonists are perpendicular to each other (Figure 1A)^13^. As a result, the two compounds use different mechanisms to activate LRH-1 but exhibit similar activation profiles in luciferase reporter assays^13^. A tendency for ligands in this class to bind unpredictably in the hydrophobic pocket has likely been a confounding factor in agonist design; however, insights from the LRH-1-RJW100 structure have provided new strategies to improve activity.

**Figure 1.**
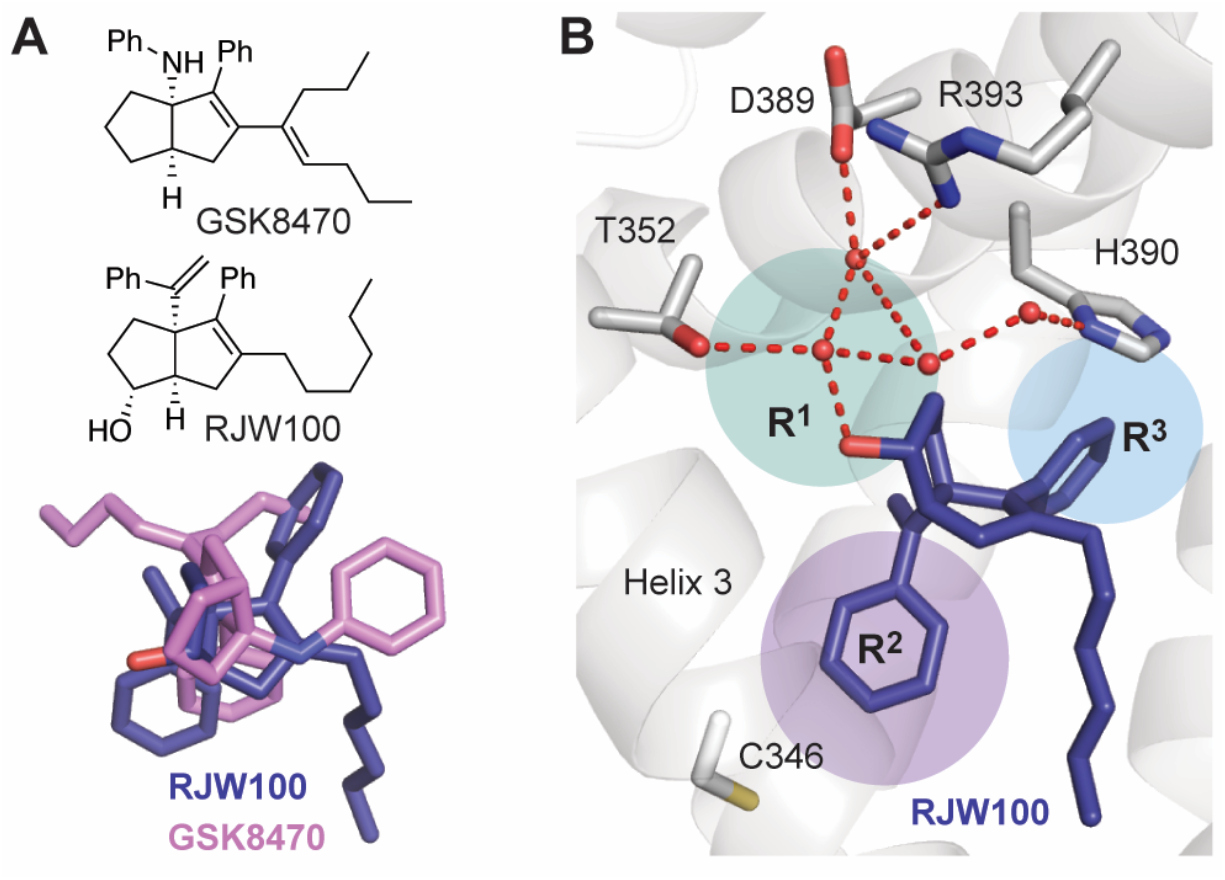
Structure-based design of LRH-1 agonists. A. Top, chemical structures of the agonists GSK8470 and RJW100. Bottom, superposition of GSK8470 and RJW100 (from PDB 3PLZ and 5L11, respectively)^12–13^ show the very different binding modes for these similar agonists. B. RJW100 interacts with LRH-1 residue Thr352 via water. The four water molecules shown coordinate a group of polar residues deep in the binding pocket. The colored circles indicate the areas targeted by modifications to the RJW100 scaffold in this work.

In the LRH-1-RJW100 crystal structure, the ligand hydroxyl group contacts a network of water molecules deep in the ligand binding pocket (Figure 1B). This water network coordinates a small group of polar residues *(e.g.* Thr352, His390, and Arg393) in an otherwise predominantly hydrophobic pocket. The *endo* RJW100 diastereomer adopts a nearly identical pose and makes the same water-mediated contact with Thr352, supporting the idea that this interaction is a primary driver of ligand orientation^13^. Using both a RJW100 analog lacking a hydroxyl group and a LRH-1 Thr352Val mutation, we demonstrated that this interaction is required for RJW100-mediated activation of LRH-1^13^. As the basis for the current studies, we hypothesized that strengthening this and other polar interactions in the vicinity could anchor ligand conformation, enabling more predictable targeting of desired parts of the pocket. We designed, synthesized, and evaluated novel compounds around the hexahydropentalene scaffold with the primary aim of strengthening polar contacts in the deep part of the binding pocket (the deep part of the pocket is hereafter abbreviated “DPP”). This systematic, structure-guided approach enabled the discovery of an agonist more potent than RJW100 by two orders of magnitude. We present three crystal structures of LRH-1 bound to novel agonists, which depict the modified polar groups projecting into the DPP. The best new agonist modulates expression of LRH-1-controlled, anti-inflammatory genes in intestinal organoids, suggesting therapeutic potential for treating inflammatory bowel diseases. This breakthrough in LRH-1 agonist development is a crucial step in developing potential new treatments for metabolic and inflammatory diseases.

## RESULTS

### Locking the agonist in place with polar interactions

Our structural studies have revealed that highly similar LRH-1 synthetic agonists can bind unpredictably within the hydrophobic binding pocket, which has presented a challenge for improving agonist design in a rational manner^13^. We reasoned that strengthening contacts within the DPP may anchor synthetic compounds in a consistent orientation and improve potency. To evaluate this hypothesis, we synthesized RJW100 analogs with bulkier polar groups in place of the RJW100 hydroxyl (R^1^), aiming to displace bridging waters and to generate direct interactions with Thr352 or other nearby polar residues (Figure 1B). In parallel, we synthesized compounds designed to interact with other sites in the DPP by (1) modifying the external styrene (R^2^) to promote interactions with helix 3 or to fill a hydrophobic pocket in the vicinity or (2) incorporating hydrogen bond donors at the *meta* position of the internal styrene (R^3^) to promote hydrogen bonding with His390 (also via water displacement) (Figure 1B). To prepare this compound library, we utilized a diastereoselective variant of Whitby’s zirconecene-mediated Pauson-Khand-type cyclization^14^ This highly modular approach unites three readily available precursors (an enyne, an alkyne, and 1,1-dibromoheptane) to generate all-carbon bridgehead [3.3.0]-bicyclic systems with varying functionality at positions R^1^, R^2^, and R^3^ (Figure 2A). R^1^ was most conveniently varied through modification of the RJW100 alcohol to yield derivatives **1–8**, which were synthesized separately as both the *endo* (N) or *exo* (X) diastereomers (Figure 2B). Oxygen-linked analogs **3** and **5** were formed directly from the diastereomerically-appropriate parent alcohol. Nitrogen-linked analogs **1, 2, 4, 6**, and **8** were prepared through alcohol activation (mesylate **S4**) and substitution (azide **S2**, nitrile **S3**) (Figure 2B and online supplementary materials). Alteration of R^2^ was accomplished by introducing phenylacetylene derivatives as the alkyne in the cyclization step (Figure 2A), generating **9–15** (Figure 2B). R^3^ variants **16–23** were prepared using functionalized enyne starting materials. Detailed chemical syntheses of all intermediates and tested compounds are provided in the online supplementary materials.

**Figure 2.**
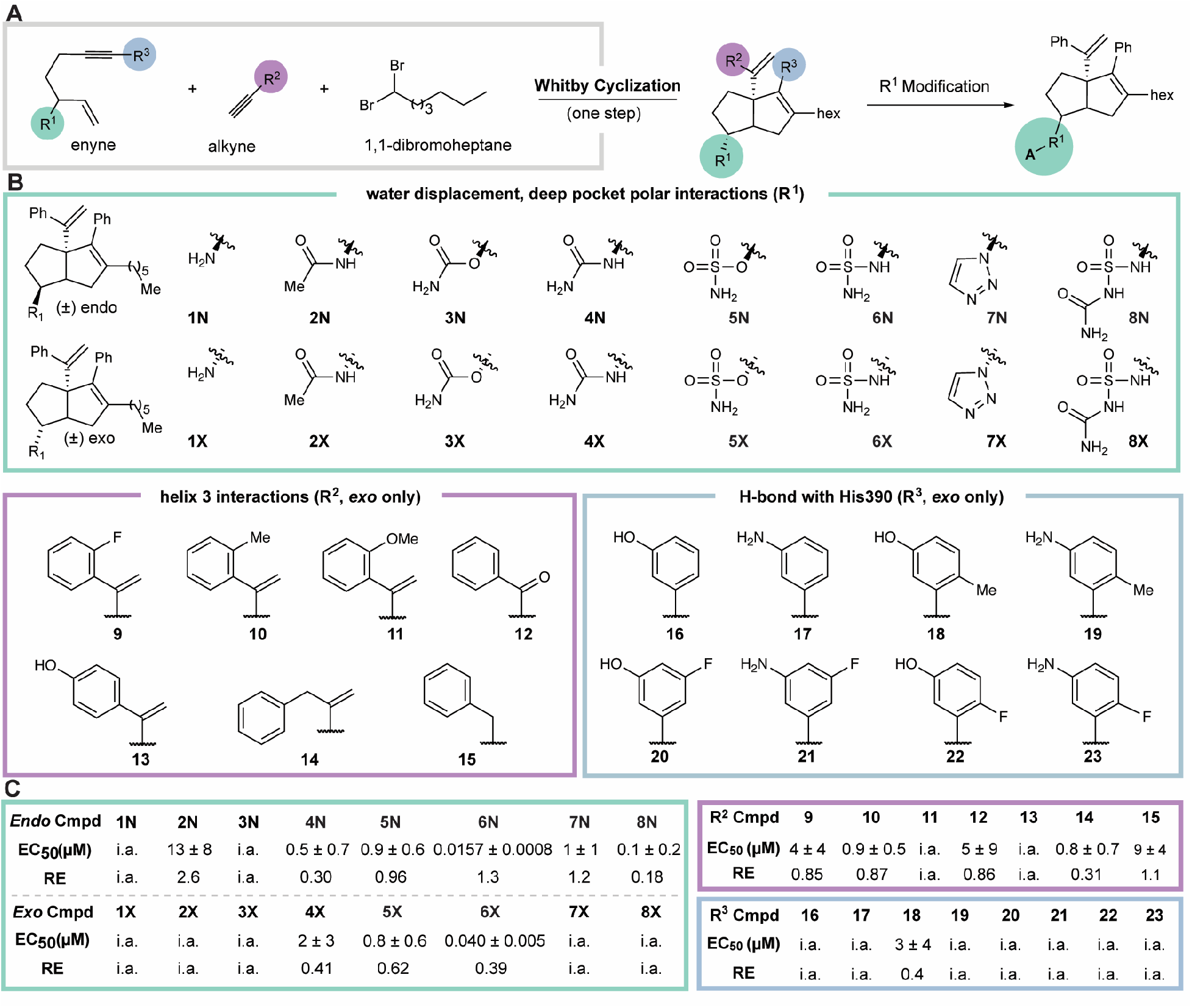
Synthesis of LRH-1-targeted compounds. A. Overview of synthetic strategy used to generate agonists based on modification of the [3.3.0]-bicyclic hexahydropentalene scaffold. B. Modifications to the scaffold evaluated in this study, grouped by site of modification by colored boxes. C. Summary of EC_5_0 and efficacy relative to RJW100 (RE). RE was calculated as described in the methods section. RJW100 RE = 1.0 and EC_50_ = 1.5 +/− 0.4 μM. The abbreviation “i.a.” refers to inactive compounds for which EC_50_ values could not be calculated.

### Discovery of the first low nanomolar LRH-1 agonist

We evaluated the new compounds using differential scanning fluorimetry (DSF), since entropic gain from displacement of buried water molecules or favorable energetics from bond formation would result in global stabilization of the LRH-1-agonist complex. DSF assays were paired with cellular luciferase reporter assays to determine effects on LRH-1 transcriptional activity. Luciferase data are summarized in Figure 2C and dose-response curves are shown in Figure S1.

As previously observed^13,15^, RJW100 stabilizes the LRH-1 ligand binding domain (LBD) by around 3 °C relative to a PL ligand in DSF assays (Figure 3A). While the R^2^- and R^3^-modified compounds (**9-23**) destabilize the receptor relative to RJW100 (Figure 3A) and tend to be poor activators (Figure 2C, S1), certain R^1^ modifications are highly stabilizing, with T_m_ values 3-8 °C higher than RJW100 (Figure 3A and online supplementary material). There is a striking correlation between potency in luciferase reporter assays and LRH-1 stabilization by DSF for the R^1^-modified compounds, where lower EC_50s_ are associated with higher Tm values (Pearson correlation coefficient = −0.71; p = 0.0009, Figure 3B). This correlation provides a direct link between cellular activity and receptor stabilization and suggests that improved potency is due to specific polar interactions mediated by the R^1^ group.

**Figure 3.**
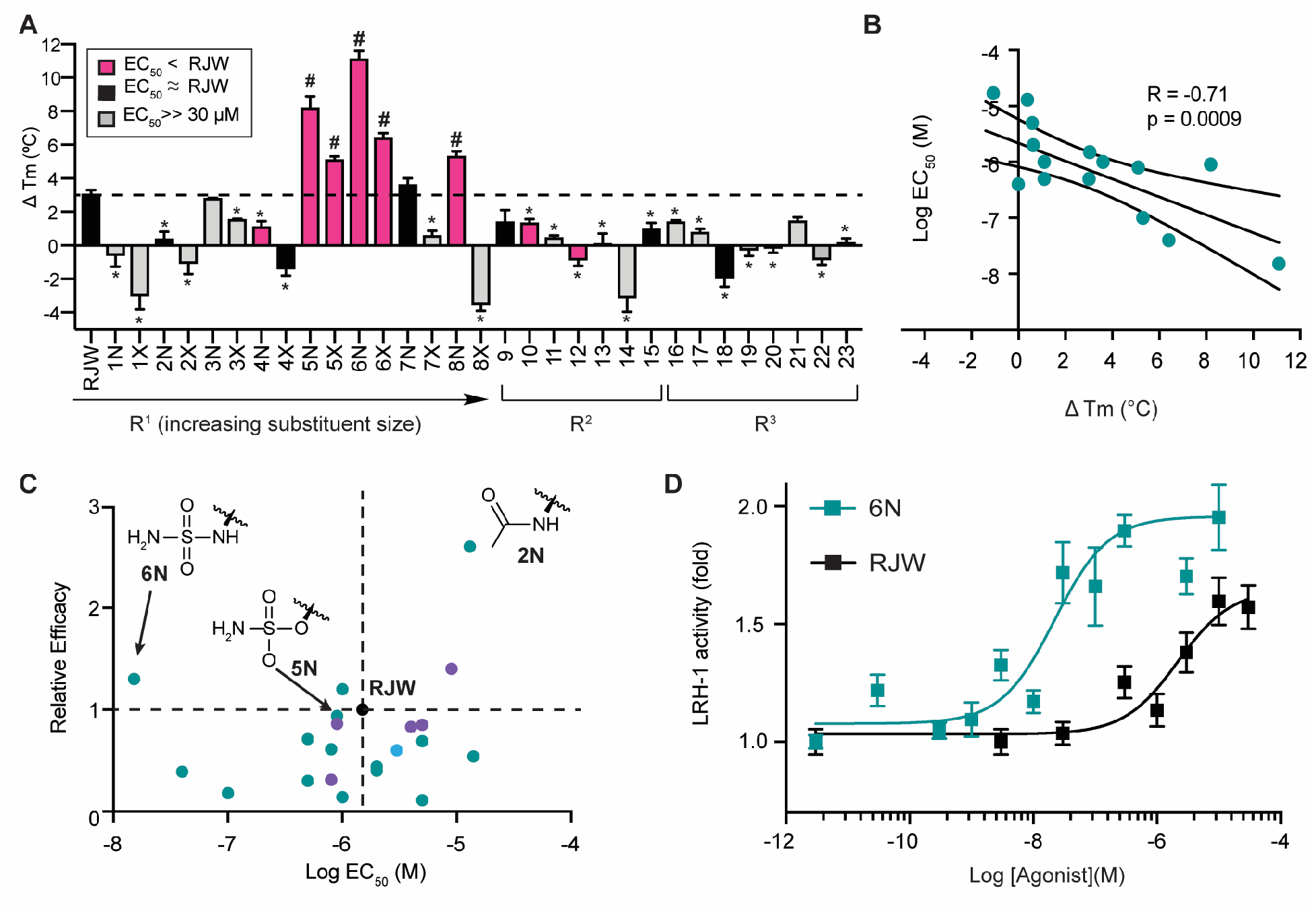
Optimization of R^1^ modification improves potency by two orders of magnitude. A. DSF assays demonstrate that the site of modification, R^1^ substituent size, and stereochemistry affect global LRH-1 stabilization. Colored bars represent EC_50s_ relative to RJW100 as indicated in the legend. Each bar represents three experiments conducted in triplicate. *, p< 0.05 for Tm decrease *versus* RJW100. #, p< 0.05 Tm increase *versus* RJW100. Significance was assessed by one-way ANOVA followed by Dunnett’s multiple comparison’s test. *Dotted line* indicates the Tm change induced by RJW100 relative to the phospholipid agonist, DLPC. B. Scatter plot showing the correlation between Tm. shift in DSF assay (x-axis) and EC_50_ from luciferase reporter assays (y-axis) for the R^1^- modified compounds. Data were analyzed by linear regression (curved lines are the 95% confidence interval. C. Scatter plot comparing potency (EC_50_) and efficacy relative to RJW100 (relative efficacy, RE) for all compounds for which EC_50_ values could be calculated. Dots are color-coded by site of modification (as indicated in Figure 2). The black dot is RJW100. The EC_50_ values and efficacies of compounds **2N, 5N**, and **6N** are indicated. Relative efficacy was calculated as described in the methods section. D. Dose response curves comparing **6N** and RJW100 in luciferase reporter assays. Each point represents the mean ± SEM for three experiments conducted in triplicate.

The R^1^ modifications are diverse, ranging from small to large polar groups, including hydrogen bond donors and acceptors and *endo* and *exo* diastereomers (Figure 2B). Both size and stereochemistry of the R^1^ group are important for activity. Mid-sized polar groups, mainly tetrahedral in geometry, tend to increase potency relative to RJW100 (Figure 2C). The close relationship between R^1^ size, agonist potency, and LRH-1 stabilization is evident looking at DSF results, where a strong peak in stabilization occurs for compounds **5 – 6** and **8N** (Figure 3A). Another strong trend among the data is that *endo* diastereomers are better activators (and more stabilizing) than the corresponding *exo* diastereomers (as seen for the triazoles **7**, sulfamides **6**, and acetamides **2**, Figure 2–3). While the compounds display a wide range of potencies and efficacies, the *endo* sulfamide (**6N**) stands out as being the most potent (Figure 3C). With an EC_50_ of 15.7 ± 0.8 nM, **6N** is two orders of magnitude more potent than RJW100 (Figure 3D). This is the first discovery of a low-nanomolar LRH-1 modulator, representing a leap forward in developing agonists for this challenging target.

### DPP contacts drive LRH-1 activation by 6N

The improved potency of **6N** is particularly striking considering that a very similar, highly stabilizing compound (**5N**) is not much more potent or effective for transcriptional activation than RJW100 (Figure 3C). The dramatic increase in potency for **6N** relative to **5N** is driven by replacement of oxygen with nitrogen in the R^1^ linker, as this is the only difference between the two compounds. Remarkably, this effect appears to be generalizable: a nitrogen-containing linker improves potency relative to an oxygen linker for several sets of compounds that differ only at this site (Figure 4A). The NH linker also contributes to selectivity for LRH-1 over its closest homolog, Steroidogenic Factor-1 (SF-1). Compound **6N** is a weaker activator of SF-1 than LRH-1, and **2N** (the *endo* acetamide) displays no activity against SF-1 while strongly activating LRH-1 (Figure 4C). In contrast, **5N** and RJW100 equally activate both receptors (Figure 4C).

**Figure 4.**
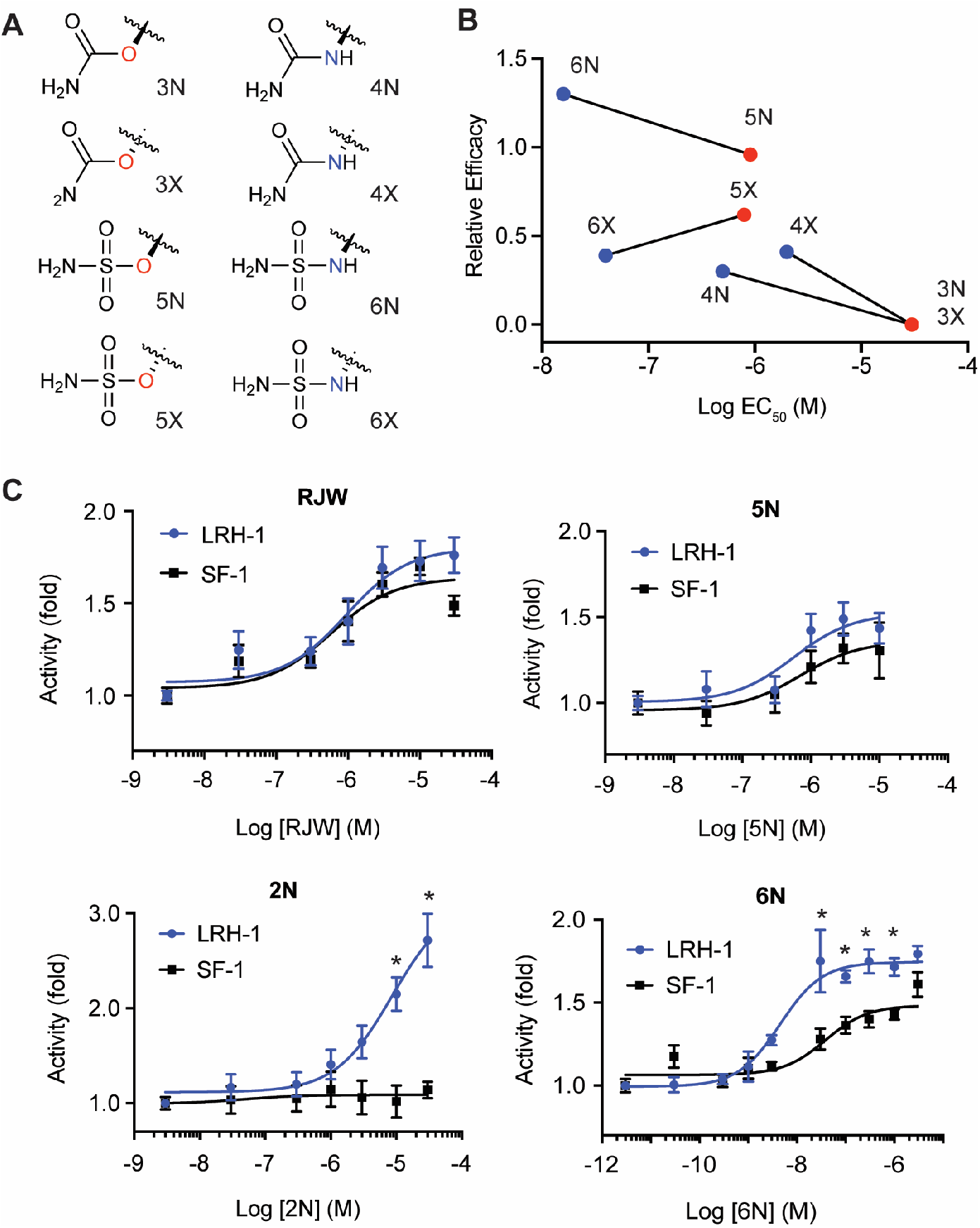
A hydrogen-bond donating nitrogen linker in the R^1^ group improves potency and selectivity. A-B. Comparison of potencies and efficacies for four sets of compounds that are identical except for the presence of a R^1^ linker containing an oxygen (red dots) or nitrogen (blue dots). Compounds with no activity at doses up to 30 μM are (**3N** and **3X**) are plotted as having EC_50s_ of 30 μM and Relative Efficacy of 0 for illustrative purposes. C. Dose response curves comparing activation of LRH-1 and SF-1 by select compounds. Significance of differences in activities of each compound for LRH-1 versus SF-1 was determined by two-way ANOVA followed by Sidak’s multiple comparison’s test. *, p < 0.05.

To investigate the role of the R^1^ linker in agonist activity and to gain insights into mechanisms underlying the potency of **6N**, we determined the X-ray crystal structure of **6N** bound to the LRH-1 LBD at a resolution of 2.23 Å (Figure 5A, Table S1). For comparison and to delineate the function of the NH-containing linker, we also determined structures of LRH-1 bound to **2N** (with an NH-linker, 2.2 Å) and **5N** (with an oxygen linker, 2.0 Å) (Table S1). The complexes were crystallized with a fragment of the coactivator protein, Transcriptional Intermediary Factor 2 (Tif2), which is bound at the AF-2 activation function surface (AFS) at the interface between helices 3, 4, and the activation function helix (AF-H, Figure 5A). Overall protein conformation does not differ greatly and is similar to the LRH-1-RJW100 structure (root mean square deviations are within 0.2 Å). The ligands are well-defined by the electron density, with the exception of the alkyl “tails” (Figure 5B). Disorder in the tail is also seen in the *endo* RJW100 structure^12^ and may be a general feature of *endo* agonists with this scaffold.

**Figure 5.**
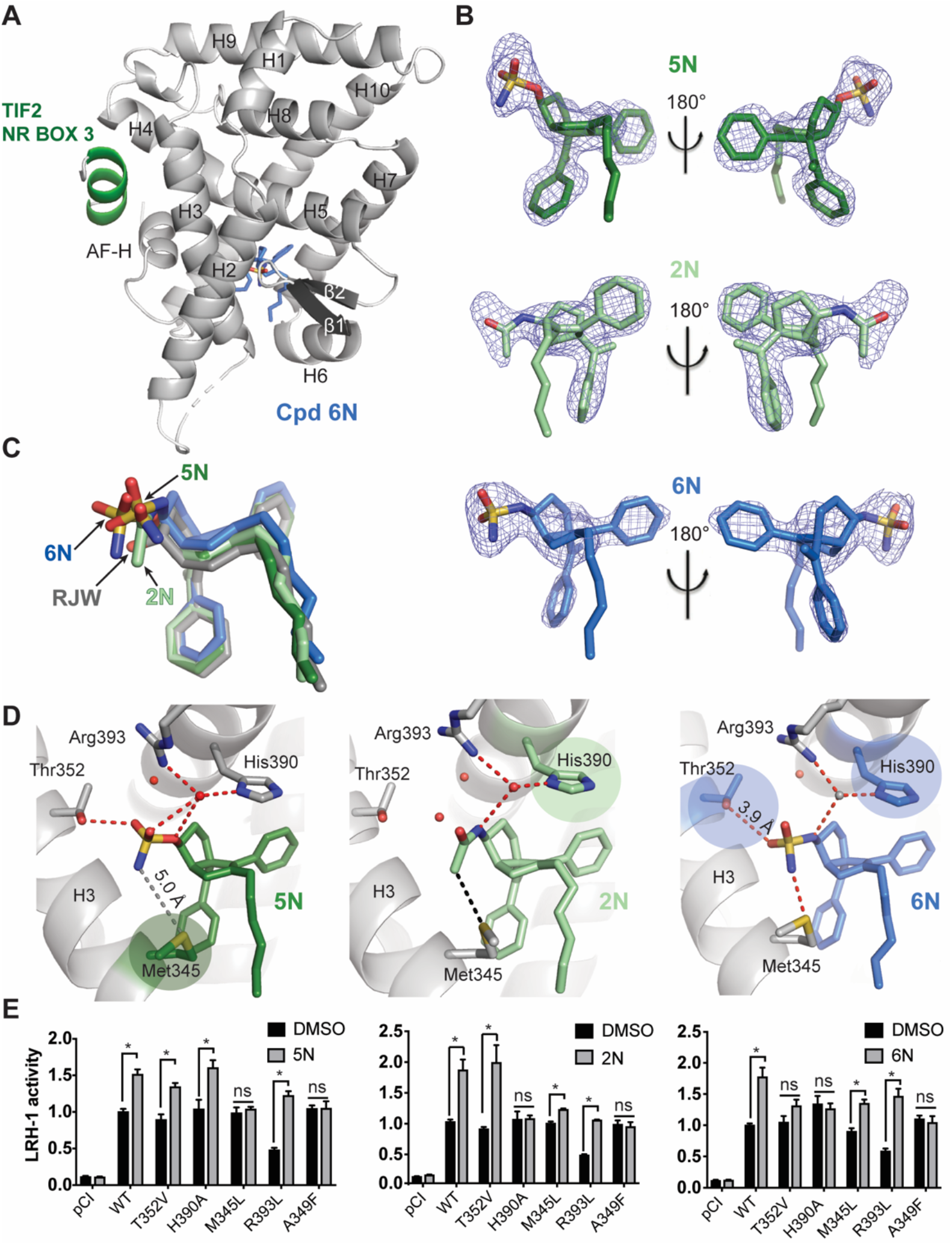
Crystal structures of LRH-1 bound to novel agonists. A. Overall structure of the LRH-1 LBD (grey) bound to **6N** (blue sticks). Tif2 is shown in green. The dotted line indicates a disordered region that could not be modeled. B. Omit maps for **2N, 5N**, and **6N**. Maps are F_o_-F_c_, contoured at 2.5σ. C. Superposition of ligands from the crystal structures showing a consistent position of the cores of the modified agonists compared to RJW100. D. Close view of LRH-1 binding pocket with either **5N, 2N**, or **6N** bound showing a subset of interactions made by the R^1^ groups. Colored circles highlight interactions that are important for LRH-1 activation by each agonist. Red spheres are water molecules (the grey sphere in the LRH-1-**6N** structure is a water molecule typically present in the LRH-1 pocket that could not be modeled due to poor crystallographic order). Red dotted lines indicate hydrogen bonds, and black dotted lines indicate hydrophobic contacts. The interaction indicated by the grey dotted line in the LRH-1-**5N** structure is outside of hydrogen-bonding distance in the structure but important for activity in mutagenesis studies. E. Luciferase reporter assays showing how the interactions made by the agonists affect LRH-1 activity. The A349F mutation occludes the DPP and was used as a negative control. Each bar represents the mean ± SEM for three independent experiments conducted in triplicate. Cells were treated with 10 μM **2N**, 10 μM **5N**, or 0.3 μM **6N** for 24 hours (concentrations chosen based on agonist EC_50_ toward wild-type LRH-1). *, p < 0.05 by two-way ANOVA followed by Dunnett’s multiple comparisons test. PDB codes for the structures are as follows: LRH-1-**5N**, 6OQX; LRH-1-**6N**, 6OQY; LRH-1-**2N**, 6OR1.

One of the main goals for these studies was to develop ligands that bind with consistent positions of the bicyclic cores. These structures demonstrate that this strategy was successful. Superposition of RJW100, **2N 5N**, and **6N** from the crystal structures shows nearly identical conformation of the agonists’ cores and phenyl groups, with slight variation in the positions of the R^1^ headgroups (Figure 5C). All three headgroups protrude into the DPP, filling space typically occupied by one or more water molecules and making several polar contacts (Figure 5D).

While the binding modes of the three agonists are similar, mutagenesis studies show that they activate LRH-1 through different mechanisms (Figure 5E). The first major difference is with the Thr352 interaction. Both **5N** and **6N** directly interact with Thr352, but the differential impact of a Thr352Val mutation shows that this interaction only contributes to agonist-mediated LRH-1 activity in the case of **6N** (Figure 5E). Compound **2N** is not well-positioned to interact with the water coordinating Thr352 due to the planar geometry of the R^1^ acetamide group, and the Thr352Val mutation has no effect on LRH-1 activity (Figure 5E). The agonists also demonstrate a differential reliance on the interaction with M345: **5N** is unable to activate a Met345L LRH-1 mutant, but **6N** and **2N** activate it significantly above basal levels (Figure 5E).

We were particularly interested in how interactions made by the NH linker contribute to agonist activity. All three agonists are positioned to make water-mediated hydrogen bonds with LRH-1 residue His390 via the R^1^ linkers (Figure 5D). In the case of **6N**, we were unable to model the bridging water molecule seen in the other two structures (and in other published LRH-1 LBD structures^13, 15–19^) due to weak electron density. The weak density for the water molecule is likely a consequence of poor crystallographic order, since very few waters could be modeled in this structure (24 total, unusual for a 2.2 Å structure). However, luciferase reporter assays with LRH-transcriptional activity (Figure 5E). Compound **2N** is also unable to activate the LRH-1 His390Ala mutant, supporting the idea that a productive water-mediated interaction with His390 is made by the NH-linker, (Figure 5E). Compound **5N**, with an oxygen linker, interacts with His390 with both the linker and sulfonyl oxygens (Figure 5D). However, **5N** does not utilize the His390 interaction for activation, since mutating His390 to alanine has no effect on its ability to activate LRH-1 (Figure 5E). Therefore, while **5N** and **6N** make very similar contacts, the presence of a hydrogen bond donor in the R^1^ linker is uniquely able to drive activation of LRH-1 via His390. This provides a potential mechanism through which a nitrogen linker increases agonist potency.

### Compound 6N stabilizes the AFS, strengthens allosteric signaling, and promotes coactivator recruitment

To investigate how **6N** alters LRH-1 dynamics to drive receptor activation, we determined its effects of LRH-1 conformation in solution using hydrogen-deuterium exchange (HDX) mass spectrometry. The most significant changes occur at sites involved in ligand-driven recruitment of coregulators: (1) the activation function surface (AFS) and (2) the helix 6/ β-sheet area (AF-B) involved in allosteric signaling to the AFS^17–18^. Relative to RJW100, **6N** destabilizes N-terminal portion of helix 7 and stabilizes the loop between helices 6 and 7 (Figure 6A). Rigidification of the loop between these helices may induce pressure to unwind helix 7, which could explain this pattern of motion. In addition to these changes near AF-B, **6N** strongly stabilizes a portion of helix 4 near the AFS (Figure 6A). Compound **5N** stabilizes the same region of helix 4, but it also destabilizes helix 12 (Figure 6B).

**Figure 6.**
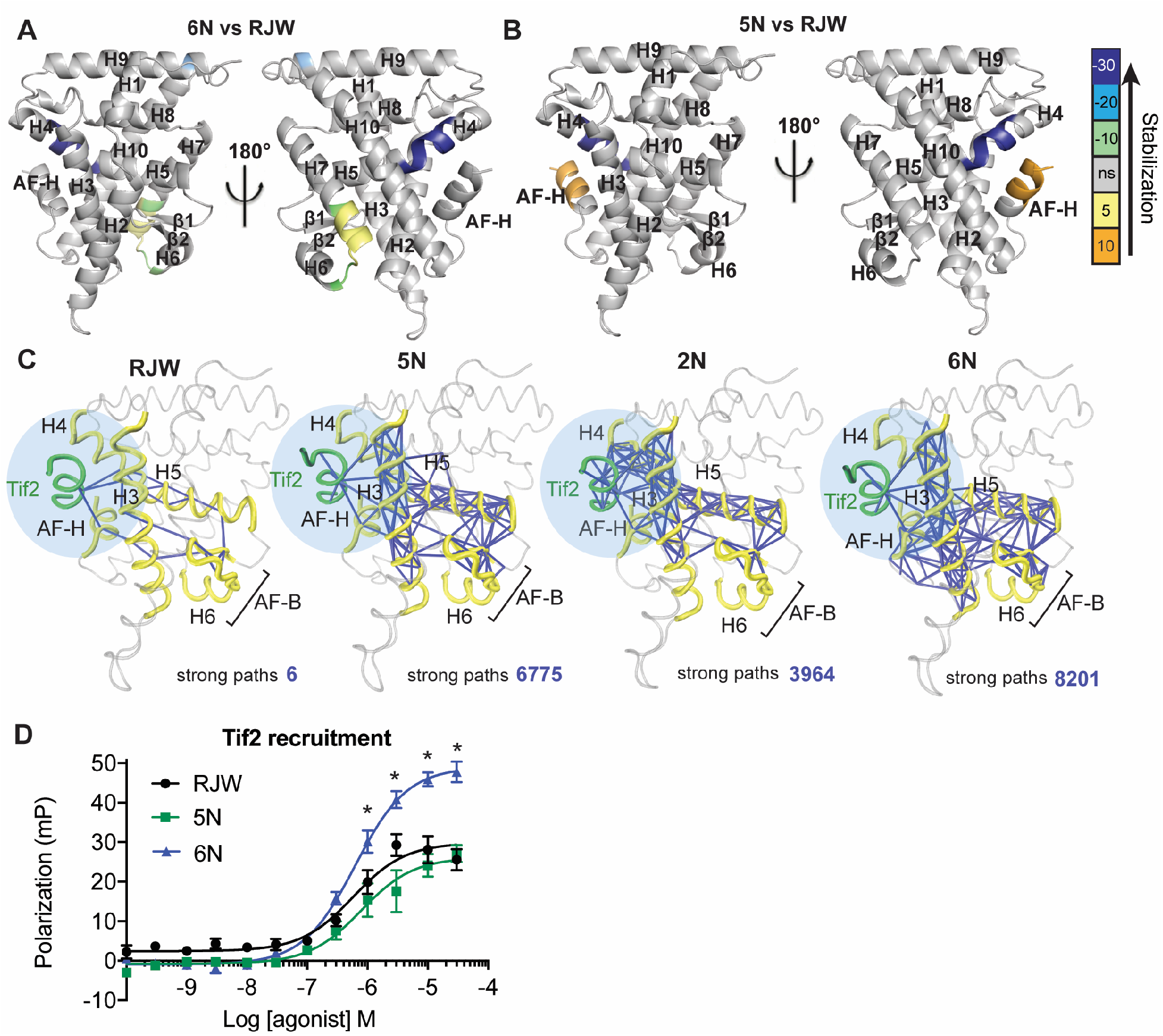
Compound 6N promotes allosteric communication to the AFS and coactivator recruitment. A-B. Differential HDX comparing **5N** to RJW100 (A) or **6N** to RJW100 (B). Color bar indicates the percent difference in deuterium uptake when **5N** or **6N** is bound compared to RJW100. A positive number indicates more deuterium exchange, indicating relative destabilization. A negative number indicates relative stabilization. C. MDS results showing the strongest suboptimal paths (blue lines) between AF-B and the Tif2 coactivator (green) when the indicated agonists are bound. The AFS is highlighted in light blue. D. Compound **6N** promotes recruitment of the Tif2 coactivator to purified LRH-1 LBD in a fluorescence polarization-based binding assay. Each point represents the mean +/− SEM for three independent experiments conducted in triplicate. *, p< 0.05 by two-way ANOVA followed by Sidak’s multiple comparison’s test.

Since **6N** alters LRH-1 conformation at AF-B and the AFS, we hypothesized that it increases communication between these two sites. To quantify the predicted strength of agonist-driven communication between AF-B and the AFS, we conducted 1 μs molecular dynamics simulations (MDS) using the crystal structures as starting models. Correlated motions of residues within a protein facilitate allosteric coupling between distant sites^20–22^. Communication paths can traverse thousands of possible routes through the receptor, and the chains of residues with the strongest patterns of correlated motion—the optimal path and a subset of suboptimal paths— are thought to convey the most information^23–24^. We therefore constructed dynamical networks of LRH-1-agonist complexes, using calculated covariance to weight the strength of communication between pairs of residues. The resulting covariance matrices were used to identify the strongest suboptimal paths facilitating communication between AF-B and Tif2 coactivator (bound at the AFS). The number of strong paths markedly increases when **2N, 5N**, or **6N** are bound compared to RJW100, with **6N** exhibiting the strongest communication between these sites (Figure 6C). There are also significant differences in the directionality of the paths promoted by each agonist. Although all paths traverse helix 5, indicating that correlated motion is induced in this region, compounds **2N, 5N**, and **6N** also induce strong communication along helix 3. Compound **6N** also induces highly interconnected communication within the AFS and the Tif2 coactivator, including significant involvement of the AF-H. This important helix in the AFS is notably excluded from the paths when the other agonists are bound (Figure 6C).

The stabilization of the AFS by **6N** is associated with enhanced coactivator recruitment. In a fluorescence-polarization based coregulator binding assay, RJW100, **5N**, and **6N** dose-dependently recruit fluorescein-labeled Tif2 peptide to LRH-1 and exhibit similar EC_50s_ (50% of maximum Tif2 binding occurs with ~600-700 nM agonist, Figure 6D). Each curve reaches a well-defined plateau that indicates the maximum response with saturating concentrations of agonist; however, curve maxima are lower for RJW100 and **5N** than **6N** by 50–60%, which is characteristic of partial agonists. Although the endogenous ligand has not been identified for comparison, **6N** behaves more like a full agonist than **5N** or RJW100 in this assay. Therefore, we have elucidated a novel mechanism of action utilized by **6N**, whereby specific interactions by the sulfamide and R^1^ linker promote allosteric signaling to the AFS, stabilizing the site of coactivator interaction and increasing Tif2 association.

### Compound 6N promotes expression of intestinal epithelial steroidogenic genes in humanized LRH-1 mouse enteroids

The discovery of the first highly potent LRH-1 agonist provides the opportunity to elucidate ligand-regulated transcriptional pathways controlled by this receptor. LRH-1 controls local steroid hormone production in the gut epithelium^25–26^, and overexpression of LRH-1 reduces inflammatory damage in immunologic mouse models of enterocolitis^10^. These findings suggest therapeutic potential for LRH-1 agonists in inflammatory bowel diseases (IBD). The recent development of methods to culture organoids of intestinal crypts (enteroids, Figure 7A) has provided an excellent research tool for drug discovery for IBD^27^. When stimulated with inflammatory cytokines, enteroids mimic features of gut epithelia in IBD^10,27^. To investigate antiinflammatory properties of **6N**, we measured the effects of the new agonist on gene expression in humanized LRH-1 mouse enteroids in the context of Tumor Necrosis Factor alpha (TNF-α)-induced inflammation. Expression of human LRH-1 in the enteroids was verified by qRT-PCR (Figure 7B). Treatment with 1 *μ*M **6N** significantly increased mRNA expression of the steroidogenic enzymes Cyp11a1 and Cyp11b1, which are Lrh-1 transcriptional targets (Figure 7C). There was a concomitant increase in expression of the anti-inflammatory cytokine IL-10 and a decrease in expression of the inflammatory cytokines IL-1β, and TNF-α (Figure 7D-E). These data suggest a role for **6N** in reducing inflammation in the gut via upregulation of steroidogenesis. Although the involvement of LRH-1 in IBD is clear from gain- and loss-of function studies^10^, the finding that epithelial steroidogenesis can be stimulated by an agonist demonstrates the tremendous potential for LRH-1 as a drug target for this disease.

**Figure 7.**
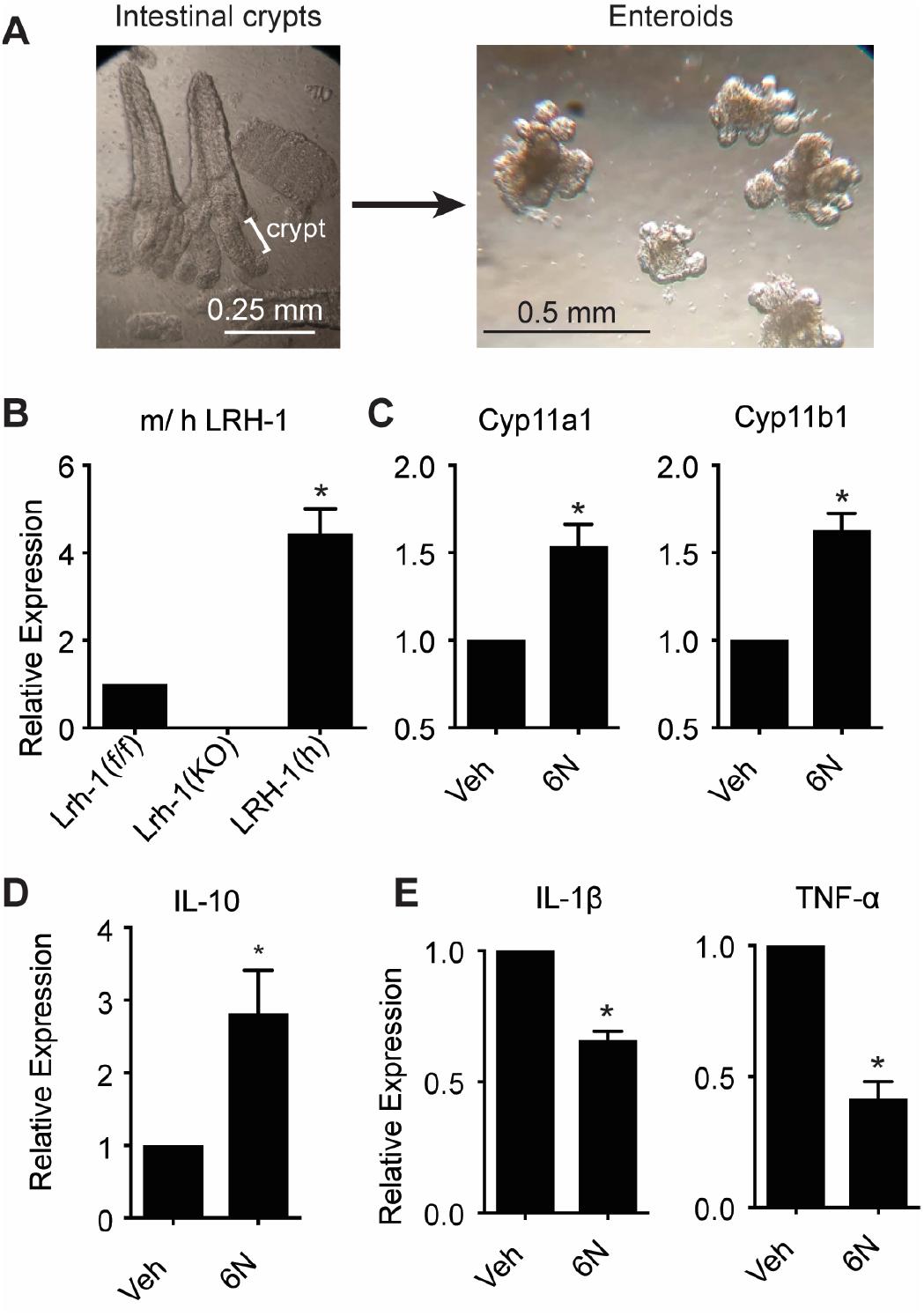
Compound 6N induces intestinal epithelial steroidogenesis in humanized LRH-1 mouse enteroids. A. Enteroids were generated through isolation and culture of intestinal crypts from mice expressing human LRH-1. B. mouse or human (m/h) LRH-1 mRNA expression in the intestinal enteroids of Lrh-1^f/f^, Lrh-1^KO^ and LRH-1^h^ mouse line. C. Compound **6N** induces mRNA expression of steroidogenic enzyme Cyp11a1 and Cyp11b1. D. Compound **6N** induces anti-inflammatory cytokine IL-10. E. Compound **6N** reduces inflammatory cytokine IL-1β (*Left*) and TNFa (*Right*). Error bars represent the SEM from 3-4 biological replicates. *, p < 0.05 (paired student’s t-test).

## DISCUSSION AND CONCLUSION

While the therapeutic potential of LRH-1 is widely recognized, this receptor has been difficult to target with synthetic modulators. Agonists with the hexahydropentalene scaffold^11–12^ (such as RJW100) are promising and have been used in several studies to probe LRH-1 biology^28–32^. However, we have shown that small modifications to this scaffold can greatly affect binding mode^13^. By exploiting a novel polar interaction in the LRH-1 DPP, we have overcome this challenge and have made substantial progress in agonist development. Systematic variation of three sites on the RJW100 scaffold has revealed a robust structure-activity relationship. The modifications to the styrene sites that we examined (R^2^ and R^3^) do not significantly improve performance and often ablate activity; however, modifications at R^1^ increase potency in transactivation assays (Figure 2C, S1). The increased potency is associated with global receptor stabilization by DSF promoted by tetrahedral, polar R^1^ substituents with *endo* stereochemistry (Figure 3). In addition, the composition of the R^1^ group, particularly the linker, is critical for activity. This is exemplified through the comparison of **5N** and **6N**, which differ only at the R^1^ linker. Compound 6N utilizes interactions with both Thr352 and His390 to activate LRH-1, the latter of which is likely mediated by the linker nitrogen (Figure 5D). This novel binding mode leads to a distinct mechanism of action for **6N** compared to similar, less potent compounds, inducing conformational changes at AF-B, stabilization of the AFS, and increasing coactivator association (Figure 6). Results from MDS support the idea that **6N** promotes very strong allostery to the AFS, evidenced in the strong communication between the AF-B and the AFS predicted to occur when **6N** is bound compared to less potent agonists (Figure 6).

With three separate crystal structures, we demonstrated that polar modifications at the RJW100 R^1^ group do not cause major repositioning of the scaffold (Figure 5), supporting our hypothesis that this polar group acts as an important anchor point. This finding was not only key to the success of the current study, but it will also greatly benefit future work. The ability to anchor the scaffold consistently provides an opportunity to tune for additional desired effects, such as solubility or selectivity. Moreover, the trajectory of the alkyl “tails” of these molecules is amenable for introduction of modifications that could engage residues near the mouth of the pocket in a PL-like manner^17, 19^. Initial studies in this vein have been fruitful, leading to the discovery of highly active compounds^33^. Finally, the establishment of a predictable binding mode may open avenues for antagonist design; for example, by modifying the scaffold to promote displacement of the AF-H and recruitment of corepressors. This approach has been successful for other nuclear receptors^34–35^ and could generate LRH-1 antagonists useful as therapeutics for certain cancers in which LRH-1 is aberrantly active^36–42^. This is an active area of research in our laboratory.

In conclusion, a systematic, structure-guided approach has resulted in the discovery of the first low nanomolar LRH-1 agonist and elucidated a novel mechanism of action. This agonist has great potential as a tool to uncover novel aspects of LRH-1 biology and as a therapeutic for IBD and obesity-associated metabolic diseases. Equally important, the discovery of elements that stabilize the orientation of the hexahydropentalene scaffold and drive activation of LRH-1 is invaluable for understanding ligand-regulation of this receptor and for future modulator design.

## EXPERIMENTAL SECTION

### General Chemical Methods

All reactions were carried out in oven-dried glassware, equipped with a stir bar and under a nitrogen atmosphere with dry solvents under anhydrous conditions, unless otherwise noted. Solvents used in anhydrous reactions were purified by passing over activated alumina and storing under argon. Yields refer to chromatographically and spectroscopically (^1^H NMR) homogenous materials, unless otherwise stated. Reagents were purchased at the highest commercial quality and used without further purification, unless otherwise stated. n-Butyllithium (n-BuLi) was used as a 1.6 M or a 2.5 M solution in hexanes (Aldrich), was stored at 4°C and titrated prior to use. Organic solutions were concentrated under reduced pressure on a rotary evaporator using a water bath. Chromatographic purification of products was accomplished using forced-flow chromatography on 230-400 mesh silica gel. Thin-layer chromatography (TLC) was performed on 250μm SiliCycle silica gel F-254 plates. Visualization of the developed chromatogram was performed by fluorescence quenching or by staining using KMnO4, p-anisaldehyde, or ninhydrin stains. ^1^H and ^13^C NMR spectra were obtained from the Emory University NMR facility and recorded on a Bruker Avance III HD 600 equipped with cryo-probe (600 MHz), INOVA 600 (600 MHz), INOVA 500 (500 MHz), INOVA 400 (400 MHz), VNMR 400 (400 MHz), or Mercury 300 (300 MHz), and are internally referenced to residual protio solvent signals. Data for ^1^H NMR are reported as follows: chemical shift (ppm), multiplicity (s = singlet, d = doublet, t = triplet, q = quartet, m = multiplet, dd = doublet of doublets, dt = doublet of triplets, ddd= doublet of doublet of doublets, dtd= doublet of triplet of doublets, b = broad, etc.), coupling constant (Hz), integration, and assignment, when applicable. Data for decoupled ^13^C NMR are reported in terms of chemical shift and multiplicity when applicable. High Resolution mass spectra (HRMS) were obtained from the Emory University Mass Spectral facility. Gas Chromatography Mass Spectrometry (GC-MS) was performed on an Agilent 5977A mass spectrometer with an Agilent 7890A gas chromatography inlet. Liquid Chromatography Mass Spectrometry (LC-MS) was used to obtain low-resolution mass spectra (LRMS) and was performed on an Agilent 6120 mass spectrometer with an Agilent 1220 Infinity liquid chromatography inlet. Purity of all tested compounds was determined by HPLC analysis, using the methods given below (as indicated for each compound).

Method A: A linear gradient using water and 0.1 % formic acid (FA) (Solvent A) and MeCN and 0.1% FA (Solvent B); t = 0 min, 30% B, t = 4 min, 99% B (held for 1 min), then 50% B for 1 min, was employed on an Agilent Poroshell 120 EC-C18 2.7 micron, 3.0 mm × 50 mm column (flow rate 1 mL/min) or an Agilent Zorbax SB-C18 1.8 micron, 2.1 mm × 50 mm column (flow rate 0.8 mL/min). The UV detection was set to 254 nm. The LC column was maintained at ambient temperature.

Method B: A linear gradient using water and 0.1 % formic acid (FA) (Solvent A) and MeCN and 0.1% FA (Solvent B); t = 0 min, 70% B, t = 4 min, 99% B (held for 1 min), then 50% B for 1 min, was employed on an Agilent Poroshell 120 EC-C18 2.7 micron, 3.0 mm × 50 mm column (flow rate 1 mL/min) or an Agilent Zorbax SB-C18 1.8 micron, 2.1 mm × 50 mm column (flow rate 0.8 mL/min). The UV detection was set to 254 nm. The LC column was maintained at ambient temperature.

Method C: An isocratic method using 75% MeCN, 35% water, and 0.1 % FA was employed on an Agilent Poroshell 120 EC-C18 2.7 micron, 3.0 mm × 50 mm column (flow rate 1 mL/min) or an Agilent Zorbax SB-C18 1.8 micron, 2.1 mm × 50 mm column (flow rate 0.8 mL/min). The UV detection was set to 254 nm. The LC column was maintained at ambient temperature.

Method D: An isocratic method using 85% MeCN, 15% water, and 0.1% FA was employed on an Agilent Poroshell 120 EC-C18 2.7 micron, 3.0 mm × 50 mm column (flow rate 1 mL/min) or an Agilent Zorbax SB-C C18 1.8 micron, 2.1 mm × 50 mm column (flow rate 0.8 mL/min). The UV detection was set to 254 nm. The LC column was maintained at ambient temperature.

### Chemical Synthesis of 2N, 5N, and 6N

The synthesis and characterization of key target compounds 2N, 5N, and 6N are outlined below. Detailed synthetic procedures and characterization data for all new compounds are provided in the supplemental section. *(Exo)5-hexyl-4-phenyl-3a-(1-pheylvinyl)-1,2,3,3,3a,6,6a-hexahydropoentalen-1-yl methanesulfonate* (S4X): A solution of RJW100 *exo* (122.5 mg, 0.3 mmol, 1.0 equiv) in dichloromethane was treated with methanesulfonyl chloride (5.0 equiv), then triethylamine (5.0 equiv) The reaction mixture was allowed to stir 1 h before concentrating and purifying on silica in 30% EtOAc/hexanes eluent. **S4X**: 139 mg, >99% yield. ^1^H NMR (500 MHz, CDCl_3_) δ 7.37 – 7.25 (m, 8H), 7.27 – 7.19 (m, 2H), 5.11 (d, *J* = 1.3 Hz, 1H), 5.01 (d, *J* = 1.3 Hz, 1H), 4.83 (d, *J* = 4.0 Hz, 1H), 2.95 (s, 3H), 2.63 (d, *J* = 9.2 Hz, 1H), 2.41 (dd, *J* = 17.4, 9.5 Hz, 1H), 2.14 (dd, *J* = 17.5, 2.0 Hz, 1H), 2.11 – 1.98 (m, 4H), 1.90 – 1.75 (m, 2H), 1.40 – 1.31 (m, 2H), 1.32 – 1.17 (m, 6H), 0. 87 (t, *J* = 7.1 Hz, 3H).

*Endo 5-hexyl-4-phenyl-3a-(1-phenylvinyl)-1,2,3,3a,6,6a-hexahydropentalen-1-amine* (**S3N**): A solution of **S4X** (139.8 mg, 0.12 mmol, 1.0 equiv) in DMF was treated with sodium azide (10.0 equiv) and the reaction was stirred 16 h at 80 °C behind a blast shield. The solution was allowed to cool to room temperature and poured over water and extracted with EtOAc three times. The combined organic layers were washed with water and brine, dried over MgSO_4_, and concentrated. The reaction mixture was purified on silica in 0-10% EtOAc/hexanes eluent. **S3N**: 117.6 mg, 95% yield. 1H NMR (600 MHz, CDCl_3_) δ 7.36 – 7.26 (m, 8H), 7.23 – 7.18 (m, 2H), 5.10 (d, *J* = 1.3 Hz, 1H), 4.94 (d, *J* = 1.3 Hz, 1H), 3.87 (ddd, *J* = 10.5, 8.8, 5.9 Hz, 1H), 2.62 – 2.51 (m, 2H), 2.16 – 2.01 (m, 4H), 1.97 – 1.88 (m, 1H), 1.79 (ddd, *J* = 12.4, 5.9, 1.8 Hz, 1H), 1.71 (td, *J* = 12.4, 5.2 Hz, 1H), 1.67 – 1.59 (m, 1H), 1.40 (p, *J* = 7.5 Hz, 2H), 1.31 – 1.19 (m, 5H), 0.87 (t, *J* = 7.2 Hz, 3H). *Endo 5-hexyl-4-phenyl-3a-(1-phenylvinyl)-1,2,3,3a,6,6a-hexahydropentalen-1-amine* (**1N**): Under nitrogen, a solution of S3N (54 mg, 0.13 mmol, 1.0 equiv) in anhydrous Et_2_O was cooled to 0 °C and treated dropwise with LiAlH_4_ (4.0M in Et_2_O, 10.0 equiv). The reaction was stirred at ambient temperature until the reaction was complete by TLC (ca. 1 h). The reaction was cooled to 0 °C, diluted with anhydrous Et_2_O, and slowly treated with water (1mL/g LiAlH_4_). Excess 4 M NaOH was added slowly and the solution was extracted with EtOAc three times. The combined organic layers were washed with Rochelle’s salt and brine, dried over MgSO_4_, and concentrated. The crude oil was purified by silica gel chromatography in 50% EtOAc/Hexanes eluent (1% triethylamine) to afford the title compounds as a colorless oil. 1N: 47.9 mg, 95% yield. Purity was established by Method C: t_r_ = 0.302 min, 98.6%. ^1^H NMR (600 MHz, CDCl_3_) δ 7.37 – 7.19 (m, 10H), 5.08 (d, *J* = 1.4 Hz, 1H), 4.94 (d, *J* = 1.5 Hz, 1H), 3.30 (ddd, *J* = 11.0, 8.8, 5.7 Hz, 1H), 2.48 (d, *J* = 17.4 Hz, 1H), 2.42 (t, *J* = 9.0 Hz, 1H), 2.12 – 2.00 (m, 2H), 1.83 – 1.78 (m, 1H), 1.73 – 1.68 (m, 2H), 1.46 – 1.37 (m, 2H), 1.35 – 1.20 (m, 8H), 0.88 (t, *J* = 7.1 Hz, 3H).

*N-((endo)-5-hexyl-4-phenyl-3a-(1-phenylvinyl)-1,2,3,3a,6,6a-hexahydropentalen-1-yl)acetamide* (**2N**) A solution of **1N** (23 mg, 0.06 mmol, 1.0 equiv) in DCM was cooled to 0 °C and treated with acetyl chloride (1.5 equiv) and triethylamine (3.0 equiv), then stirred for 1 h. The solution was diluted with water and extracted with DCM three times. The combined organic layers were washed with water and brine, dried with Na_2_SO_4_, filtered, and concentrated. The crude oil was purified on silica gel in 35% EtOAc/Hexanes eluent to afford the title compound as a colorless oil. **2N**: 21.1 mg, 83% yield. Purity was established by Method D: t_R_ = 1.00 min, 96.3 %. ^1^H NMR (500 MHz, CDCl_3_) δ 7.35 – 7.27 (m, 5H), 7.25 – 7.22 (m, 5H), 5.35 (d, *J* = 8.1 Hz, 1H), 5.06 (d, *J* = 1.5 Hz, 1H), 5.02 (d, *J* = 1.5 Hz, 1H), 4.25 (dtd, *J* = 10.5, 8.6, 6.2 Hz, 1H), 2.66 (ddd, *J* = 16.9, 8.4, 1.6 Hz, 1H), 2.14 – 2.00 (m, 4H), 1.99 (s, 3H), 1.87 (dtd, *J* = 11.7, 6.0, 2.3 Hz, 1H), 1.76 (td, *J* = 12.2, 11.7, 5.8 Hz, 1H), 1.66 (ddd, *J* = 12.7, 5.9, 2.3 Hz, 1H), 1.43 – 1.26 (m, 1H), 1.30 – 1.18 (m, 8H), 0.87 (t, *J* = 7.0 Hz, 3H). ^13^C NMR (126 MHz, CDCl_3_) δ 169.3, 154.6, 143.6, 141.7, 138.9, 137.2, 129.6, 127.9, 127.8, 127.7, 126.9, 126.6, 114.8, 69.0, 59.5, 54.4, 40.9, 33.0, 32.1, 31.6, 29.8, 29.4, 27.8, 26.0, 23.6, 22.6, 14.1. LRMS (ESI, APCI) *m/z:* calc’d for C_30_H_39_NO [M+H]^+^ 430.3, found 430.3 *(endo)-5-hexyl-4-phenyl-3a-(1-phenylvinyl)-1,2,3,3a,6,6a-hexahydropentalen-1-yl sulfamate* (**5N**): A 1M solution of sulfamoyl chloride (2.5 equiv) in DMA was cooled to 0°C. A solution of the appropriate RJW100 alcohol isomer *(endo* (224.3 mg, 0.6 mmol 1.0 equiv) in DMA was added slowly, followed by triethylamine (excess, ca. 5 equiv); the resulting solution was stirred for one hour. The solution was then diluted with water and extracted with EtOAc three times. The combined organic layers were washed with water and brine, dried with MgSO_4_, filtered, and concentrated. The crude oil was purified by silica gel chromatography in 20% EtOAc/Hexanes eluent (with 0.5% triethylamine), to afford the title compound as a clear oil. **5N**: 182 mg, 67% yield. Purity was established by Method D: tR = 1.15 min, 95.3%. ^1^H NMR (500 MHz, CDCl_3_) δ 7.35 – 7.24 (m, 8H), 7.23 – 7.15 (m, 2H), 5.11 (s, 1H), 4.92 (s, 1H), 4.87 (td, *J* = 9.1, 5.2 Hz, 1H), 4.64 (s, 2H), 2.71 (d, *J* = 9.0 Hz, 1H), 2.60 (d, *J* = 17.5 Hz, 1H), 2.17 (dd, *J* = 17.7, 9.3 Hz, 1H), 2.10 – 2.01 (m, 3H), 1.92 – 1.83 (m, 1H), 1.83 – 1.76 (m, 1H), 1.68 (td, *J* = 12.6, 5.6 Hz, 1H), 1.45 – 1.35 (m, 2H), 1.32 – 1.16 (m, 6H), 0.86 (t, *J* = 7.1 Hz, 3H). ^13^C NMR (126 MHz, CDCl_3_) δ 153.8, 143.5, 143.2, 138.5, 136.5, 129.8, 127.9, 127.7, 127.6, 127.0, 126.8, 115.7, 84.1, 68.2, 47.1, 34.9, 31.6, 31.2, 30.5, 29.8, 29.4, 27.7, 22.6, 14.1. LRMS (ESI, APCI) *m/z:* calc’d for C_28_H_36_NO_3_S [M-H]-465.3, found 465.4

*endo 5-hexyl-4-phenyl-3a-(1-phenylvinyl)-1,2,3,3a,6,6a-hexahydropentalen-1-yl sulfamide* (**6N**) A solution of **1N** (30 mg, 0.08 mmol, 1.1 equiv) in DCM was treated with triethylamine (2.0 equiv.) and solution of 2-oxo-1,3-oxazolidine-3-sulfonyl chloride (0.5 M in DCM, 1.0 equiv) (prepared according to the procedure of Borghese *et al*)^43^. The reaction was stirred at room temperature for 3 h then concentrated. The residue was treated with ammonia (0.5 M in dioxane, 1.5 equiv) and triethylamine (3.0 equiv). The solution was heated in a sealed tube at 85°C for 16 h behind a blast shield. After cooling to ambient temperature, the reaction was diluted with 3:3:94 MeOH:Et_3_N:EtOAc and passed through a pad of silica. The eluent was concentrated, and the crude oil was purified on silica in 20-30% EtOAc/hexanes eluent to afford the title compound as a colorless oil. **6N**: 21.6 mg, 60% yield. Purity was established by Method C: t_R_ = 2.0 min, 96.6 %. ^1^H NMR (600 MHz, CDCl_3_) δ 7.33 – 7.23 (m, 8H), 7.20 – 7.17 (m, 2H), 5.09 (d, *J* = 1.3 Hz, 1H), 4.96 (d, *J* = 1.3 Hz, 1H), 4.44 (s, 2H), 4.36 (d, *J* = 8.0 Hz, 1H), 3.84 – 3.77 (m, 1H), 2.62 (td, *J* = 8.9, 2.0 Hz, 1H), 2.38 (dd, *J* = 17.5, 2.0 Hz, 1H), 2.20 – 2.13 (m, 1H), 2.08 – 2.04 (m, 2H), 2.00 – 1. 95 (m, 1H), 1.74 – 1.70 (m, 2H), 1.50 – 1.43 (m, 1H), 1.42 – 1.16 (m, 8H), 0.86 (t, *J* = 7.1 Hz, 3H). ^13^C NMR (126 MHz, CDCl_3_) δ 154.1, 143.6, 142.8, 139.3, 136.6, 129.6, 127.8, 127.7, 126.9, 126.8, 115.5, 68.8, 57.2, 47.4, 35.4, 32.3, 32.0, 31.6, 29.8, 29.5, 27.9, 22.6, 14.1. LRMS (ESI, APCI) *m/z:* calc’d for C_28_H_37_N_2_O_2_S [M+H]^+^ 465.7, found 464.8

### Biology: materials and reagents

pCI empty vector was purchased from Promega. The SHP-luc and *Renilla* reporters, as well as pCI LRH-1, have been previously described^17^. The vector for His-tagged tobacco etch virus (TEV) was a gift from John Tesmer (University of Texas at Austin). The pMSC7 (LIC-HIS) vector was provided by John Sondek (University of North Carolina at Chapel Hill). The Tif2 NR Box 3 peptide was purchased from RS Synthesis. DNA oligonucleotide primers were synthesized by Integrated DNA Technologies.

### Protein purification

Purification of human LRH-1 ligand binding domain (residues 300-537) in a pMCSG7 expression vector was performed as described^13^. Briefly, protein was expressed in BL21 PLysS *E. coli,* using 1 mM IPTG for 4 hours (30°C) to induce expression. Protein was purified by nickel affinity chromatography. For DSF assays, protein eluted from the nickel column was exchanged with DLPC (5-fold molar excess overnight at 4 °C), followed by repurification by size exclusion to remove displaced lipids. The assay buffer was 20 mM Tris-HCl, pH 7.5, 150 mM sodium chloride, and 5% glycerol. Cleaved LRH-1 was then incubated with ligands overnight at 4 °C prior to repurification by size exclusion, using the same assay buffer as for DSF. Protein used for crystallography was prepared as for coregulator recruitment, except that it was sized into a buffer of 100 mM ammonium acetate (pH 7.5), 150 mM sodium chloride, 1 mM DTT, 1 mM EDTA, and 2 mM CHAPS.

### Differential scanning fluorimetry (DSF)

DSF assays were conducted on a StepOne Plus thermocycler as previously described^13,15^. Briefly, aliquots of purified LRH-1 LBD protein (0.2 mg/ ml) were incubated with saturating concentrations of ligand overnight at 4 °C. Protein-ligand complexes were heated in the presence of SYPRO orange dye at a rate of 0.5 degree/ minute. Complexes were excited at 488 nm, and fluorescence emissions at each degree Celsius were measured using the ROX filter (~600 nm). Tm values were calculated using the Bolzmann equation in GraphPad Prism, v7.

### Crystallography

Compounds **5N, 6N**, or **2N** were incubated with purified LRH-1 LBD (His tag removed) at 5-fold molar excess overnight at 4°C. The complexes were re-purified by size exclusion chromatography into the crystallization buffer (see above). Protein was concentrated to 5-6 mg/ ml and combined with a peptide from human Tif2 NR box 3 (H3N-KENALLRYLLDKDDT-CO2) at four-fold molar excess. Crystals were generated by hanging drop vapor diffusion at 18 °C, using a crystallant of 0.05 M sodium acetate (pH 4.6), 5-11% PEG 4000, and 0-10% glycerol. Crystals of **2N** with LRH-1 were generated by microseeding, using RJW100-LRH-1 crystals as the seed stocks (crystals used for seeding were grown as described)^13^.

### Structure Determination

Crystals were flash-frozen in liquid nitrogen, using a cryoprotectant of crystallant plus 30% glycerol. Diffraction data were collected remotely from Argonne National Laboratory, Southeast Regional Collaborative Access Team, Beamline 22ID. Data were processed and scaled using HKL2000^44^. Structures were phased by molecular replacement using Phenix^45^,with PBD 5L11 used as the search model. The structure was refined using phenix.refine^45^ and Coot^46^, with some additional refinement done using the PDB Redo web server^47^.

### Tissue culture

Hela cells were purchased from Atlantic Type Culture Collection and cultured in phenol red-free MEMα media supplemented with 10% charcoal-dextran-stripped fetal bovine serum. Cells were maintained under standard culture conditions.

### Reporter gene assays

Hela cells were reverse-transfected with three vectors: (1) full-length, human LRH-1 in a pCI vector, (2) a firefly reporter (pGL3 Basic) with a portion of the SHP promoter cloned upstream of the firefly luciferase gene, and (3) a constitutively active vector expressing *Renilla* luciferase under control of the CMV promoter. To study SF-1 activity, cells were transfected with the same constructs, except that full-length SF-1 (in a pcDNA3.1 vector) was overexpressed instead of LRH-1, with empty pcDNA3.1 used as the negative control. Transfections utilized the Fugene HD transfection reagent at a ratio of 5 μl per 2 μg DNA. To perform the reverse transfections, cells were trypsinized, combined with the transfection mixture, and plated at densities of 7,500 cells per well in white-walled 96-well plates. The following day, cells were treated with each compound (or DMSO control) for 24 hours. In most cases, six points in the concentration range of 0.03 – 30 μM were used (exceptions noted in figures), with a final DMSO concentration of 0.3% in all wells. Luciferase expression was measured using the DualGlo Kit (Promega). Firefly luciferase signal was normalized to *Renilla* luciferase signal in each well. EC_50_ values were calculated using three-parameter curve-fitting (GraphPad Prism, v.7). Assays were conducted in triplicate with at least two independent biological replicates. Experiments involving SF-1 activation were conducted in an identical manner, except full-length human SF-1 (in a pcDNA3.1+ vector) was overexpressed instead of LRH-1. Significance of differences in luminescence signal for LRH-1 *versus* SF-1 promoted by particular agonists was determined using two-way ANOVA followed by Sidak’s multiple comparisons test.

### Calculation of Relative Efficacy (RE)

This value was calculated from curve-fitting to data from luciferase reporter assays. To compare the maximum activities of the new compounds to RJW100, we used the formula (Max_cpd_ – Min_cpd_) / (Max_RJW100_ – Min_RJW100_), where “Max” and “Min” denote the dose response curve maximum and minimum, respectively. A RE of 0 indicates a completely inactive compound, a value of 1 indicates equal activity to RJW100, and values above 1 indicate greater activity.

### Mutagenesis

Mutations were introduced to LRH-1 in the pCI vector using the Quikchange Lightning site-directed mutagenesis kit (Ambion). Constructs were sequenced prior to use in reporter gene assays as described above.

### Model Construction for Molecular Dynamics Simulations

Four LRH-1 LBD complexes were prepared for molecular dynamics simulations. 1) LRH-1-Tif2-RJW100 (PDB 5L11), 2) LRH-1-Tif2-**5N**. 3LRH-1-Tif2-**2N**, LRH-1-Tif2-**6N**. For consistency, all structures contained LRH-1 residues 300–540. Missing residues (i.e., that could not be modeled in the structures) were added to the models used in the simulations.

### Molecular Dynamics Simulations

The complexes were solvated in an octahedral box of TIP3P water with a 10-Å buffer around the protein complex. Na^+^ and Cl^-^ ions were added to neutralize the protein and achieve physiological buffer conditions. All systems were set up using the xleap tool in AmberTools17^48^ with the ff14SB forcefield^49^. Parameters for the agonist ligands **6N, 2N** and **5N** were obtained using Antechamber^50^ also in AmberTools17. All minimizations and simulations were performed with Amber16^48^. Systems were minimized with 5000 steps of steepest decent followed by 5000 steps of conjugate gradient minimization with 500-kcal/moloÅ^2^ restraints on all solute atoms. Restraints were removed excluding the atoms in both the ligand and the Tif2 peptide, and the previous minimization was repeated. This minimization was repeated with restraints lowered to 100-kcal/moloÅ^2^. Finally, all restraints were removed for a last minimization step. The systems were heated from 0 to 300 K using a 100-ps run with constant volume periodic boundaries and 5-kcal/moloÅ^2^ restraints on all protein and ligand atoms. MD equilibration was performed for 12 ns with 10-kcal/moloÅ^2^ restraints on Tif2 peptide and ligand atoms using the NPT ensemble.

Restraints were reduced to 1 kcal/moloÅ^2^ for an additional 10 ns of MD equilibration. Then, restraints were removed, and 1000-ns production simulations were performed for each system in the NPT ensemble. A 2-fs time step was used with all bonds between heavy atoms and hydrogens fixed with the SHAKE algorithm^51^. A cutoff distance of 10 Å was used to evaluate long-range electrostatics with particle mesh Ewald and for van der Waals forces. Fifty thousand evenly spaced frames were taken from each simulation for analysis, using the CPPTRAJ module^52^ of AmberTools. The NetworkView plugin^20^ in VMD^53^ and the Carma program^54^ were used to produce dynamic networks for each system. In brief, networks are constructed by defining all protein C-α atoms as nodes, using Cartesian covariance to measure communication within the network. Pairs of nodes that reside within a 4.5-Å cutoff for 75% of the simulation are connected via an edge. Edge weights are inversely proportional to the covariance between the nodes. Networks were constructed using 500 ns of the MDS trajectories, to enable direct comparison with our previous LRH-1-RJW MDS^15^. Suboptimal paths between the AF-B and Tif2 peptide were identified using the Floyd-Warshall algorithm^55^. Suboptimal path analyses were performed using Carma and the subopt program in NetworkView. Cross-correlation matrices for C-α atoms in each system were computed with Carma.

### Coregulator Recruitment Assays

Synthetic agonists were titrated in the presence of purified LRH-1 LBD protein (2 μM) and a fluorescein (FAM)-labeled peptide corresponding to the Tif2 NR box 3 (FAM-H3N-PVSPKKKENALLRYLLDKDDT-CO_2_-) (50 nM). Protein and probe concentrations were determined from preliminary experiments titrating LRH-1 protein with no ligand added in the presence of FAM-Tif2 (2 μM was slightly above the Tif2 Kd in these experiments). Tif2 binding was detected by fluorescence polarization, using a BioTek Neo plate reader. Assays were conducted three times in triplicate, using two separate protein preparations. Significance of differences in Tif2 association at each dose was determined using two-way ANOVA followed by Tukey’s multiple comparisons test.

### Hydrogen-deuterium exchange mass spectrometry

Following cleavage of the His tag from purified LRH-1 LBD with TEV protease as described above, the protein was further purified by size exclusion chromatography into a buffer of phosphate buffered saline (pH 7.5) plus 5% glycerol. Protein purity exceeded 98% by Coomassie staining. Protein-ligand complexes were prepared by adding each ligand at 5-fold molar excess to 2 mg/ml protein and incubating overnight at 4 °C. Complexes were centrifuged to remove any aggregates prior to analysis by HDX-MS. HDX-MS was conducted using Waters’ UPLC HDX system coupled with a Q-Tof Premier mass spectrometer (Waters Corp, Milford, MA). Protein-ligand complexes were diluted 1:7 (v/v) into labeling buffer (protein buffer containing D_2_O instead of water) via an autosampler. Labeling took place at 20°C for time periods of 0, 10, 100, 1,000, and 10,000 seconds prior to quenching in with equal volume of precooled quenching buffer (100 mM phosphate, 0.5 M tris)2-carboxyethl)phosphine, 0.8% formic acid, and 2% acetonitrile, pH 2.5, 1°C). After quenching, samples were applied to a Waters enzymate pepsin column (2.1 × 30 mm). Peptides from the pepsin column were separated in-line on a Waters Acuity UPLC BEH C18 column (1.7 μM, 1.0 × 100 mm) at a flow of 40 μl/ min for 12 minutes (8-40% linear gradient, mobile phase: 0.1% formic acid in acetonitrile) at 1°C. The mass spectrometer was operated with the electrospray ionization source in positive ion mode, and the data were acquired in elevated-energy mass spectrometry mode. For internal calibration, a reference lock-mass of Glu-Fibrinopeptide (Sigma-Aldrich, St Louis, MO) was acquired along with each sample data collection. Peptides were identified by comparison to human LRH-1 protein sequence using the ProteinLynx Global SERVER (version 3.02). HDX data were processed in DynamX (version 3.0). Mass assignment for each peptide at 0 seconds of exchange was checked manually, and any assignment with a mass deviation > 0.2 Da was removed. Hydrogen-deuterium exchange protection was quantified by comparison of hydrogen exchange profiles at different time points. Peptide coverage was 99.2% for this experiment (Figure S4).

### Humanized LRH-1 mouse intestinal enteroid culture

The study protocol was approved by the Animal Care and Use Committee of Baylor College of Medicine and was in accordance with the Guide for the Care and Use of Laboratory Animals [DHHS publication no. (NIH) 85-23, revised 1985, Office of Science and Health Reports, DRR/NIH, Bethesda, MD 20205].

The humanized LRH-1 allele (LRH-1^h^) is obtained on a mouse line with a human LRH-1 transgene using the Rosa26-loxP-STOP-loxP strategy to allow villin-cre mediated expression of human LRH-1 (LRH-1^ΔΔ^) in enterocytes with knockout of the endogenous mLrh-1 (Lrh-1^f/f^). Intestinal crypt culture (enteroids) were derived from Lrh1^f/f^, Lrh-1^KO^ (Lrh1^f/f^;Villin-Cre^+^), and LRH-1^h^ (Lrh1^f/f^;hLRH1^ΔΔ^;Villin-Cre^+^) male mice (6-8 weeks old). Briefly, the small intestine was isolated and flushed with ice-cold phosphate-buffered saline (PBS), opened longitudinally, then cut into 1–2 mm pieces. Intestinal fragments were incubated in an EDTA (4 mM) containing solution at 4 °C for 60 min on a tube rocker. The intestinal fragment suspension was fractionated by vertical shaking manually and crypt-containing fractions passed through a 70-μm cell strainer for plating in Matrigel. Crypt-Matrigel suspension was allowed to polymerize at 37 °C for 15 min. Intestinal organoids were grown in base culture media (Advanced DMEM/F12 media, HEPES, GlutaMax, penicillin, and streptomycin) supplemented with growth factors (EGF, Noggin, R-spondin, R&D Systems), B27 (Life Technologies), N2 (Life Technologies), and N-acetyl cysteine (NAC, Sigma). Intestinal enteroids were passaged every 3 days. Established LRH-1^h^ enteroids were treated with mouse TNF-α overnight to provoke inflammatory changes, then treated with vehicle (DMSO) or compound 6N (1 μM) overnight. Following treatment, enteroid tissues were harvested for real time PCR.

### RNA isolation and PCR

Intestinal enteroids were washed in ice-cold PBS and suspended in Trizol solution (Sigma). RNA was isolated with RNeasy^®^ spin columns (Qiagen). DNAse-treated total RNA was used to generate cDNA using Superscript II (Quanta). Sybr green-based qPCR (Kapa Biosystems) was performed on a Roche LightCycler^®^ 480 II with primers as shown below. The ΔΔCt method was used for calculating gene expression fold changes using Rplp0 (ribosomal protein, large, P0, known as 36B4) as reference. Primer sequences were as follows:

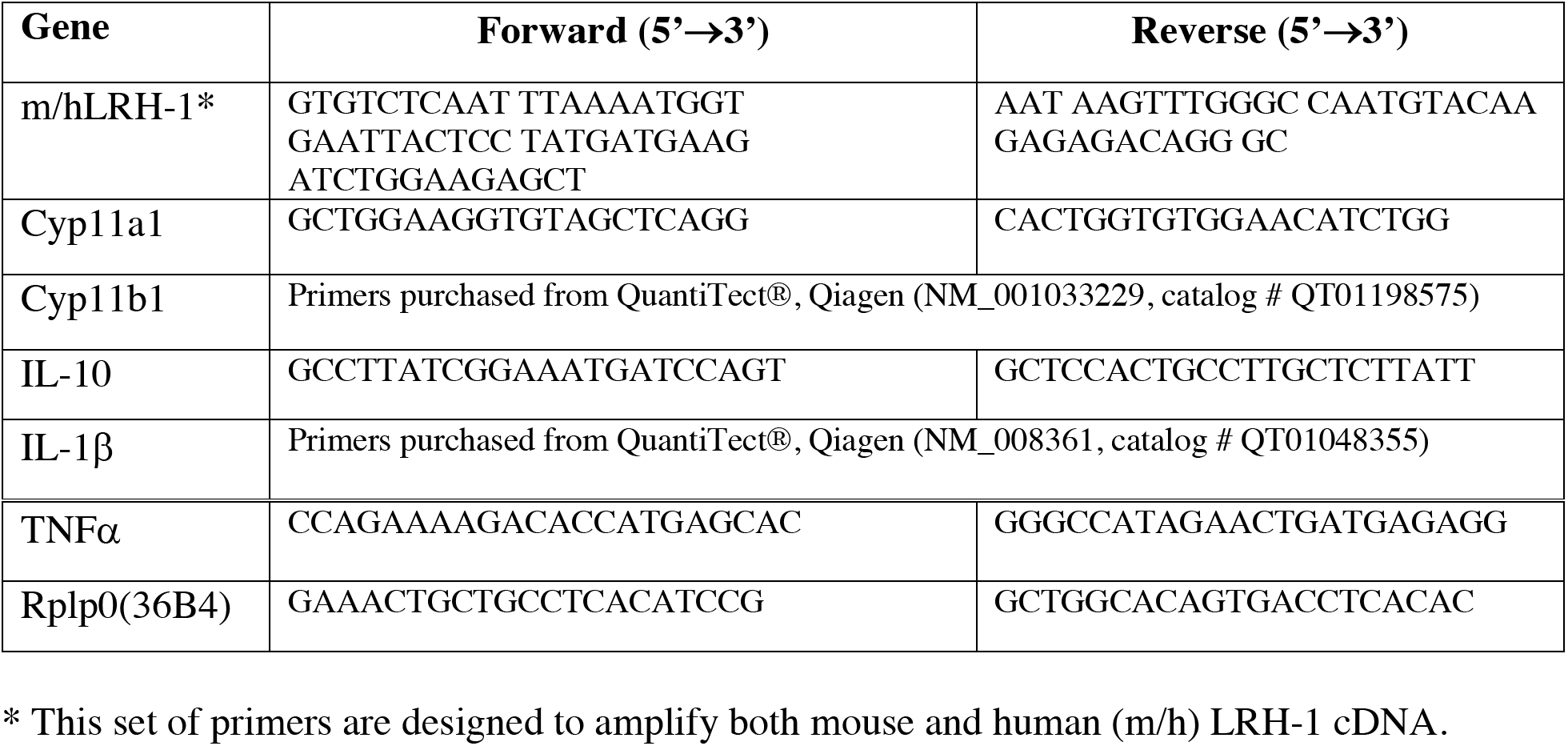

## ASSOCIATED CONTENT

### Supporting Information

Supplemental figures and tables showing dose-response curves for luciferase reporter assays, HDX deuterium uptake, crystal data collections and refinement statistics, detailed chemical syntheses are provided as a part of the supporting information.

Present Addresses

### Author Contributions

All authors contributed to this manuscript and approve the final version.

### Funding Sources

This work was supported in part by the National Institutes of Health under the following awards: T32GM008602 (S.G.M.), F31DK111171 (S.G.M.), R01DK095750 (E.A.O.), R01DK114213 (E.A.O., N.T.J., J.W.C), P30DK056338 (Texas Medical Center Digestive Disease Center pilot, H. W.). The work was also supported by an Emory Catalyst Award (E.A.O., N.T.J), NASPGHAN transition award (H.W.), and USDA ARS 3092-5-001-057 (D.D.M.).

## ACKNOWLEDGMENT

We thank the beamline staff at Argonne National Laboratory, South East Regional Collaborative Team, Beamline 22ID for support during remote data collection. We are also grateful to the Emory HDX Core, particularly Dr. Wei Dang and Dr. Renhao Li, for assistance with HDX experiments. We thank Michael Dugan for assistance with chemical synthesis.

## ABBREVIATIONS

LRH-1: Liver Receptor Homolog-1;
DLPC: dilauroylphosphatidylcholine;
DPP: deep polar portion of the LRH-1 binding pocket;
DSF: differential scanning fluorimetry;
RE: relative efficacy;
i.a.: inactive;
PL: phospholipid;
Tif2: transcriptional intermediary factor 2;
AFS: activation function surface;
AF-H: activation function helix;
SF-1: steroidogenic factor-1;
LBD: ligand binding domain;
HDX: hydrogen-deuterium exchange;
AF-B: helix 6/**β** sheet;
MDS: molecular dynamics simulation;
qRT-PCR: quantitative real time polymerase chain reaction;
veh: vehicle;
h: hour;
NR: nuclear receptor;
DMSO: dimethylsulfoxide;
Cyp11a1: Cytochrome P450 family 11 subfamily a member 1;
Cyp11b1: Cytochrome P450 family 11 subfamily b member 1;

## PDB CODES

Authors will release the atomic coordinates and experimental data upon article publication. PDB IDs have been provided in figure legends and are as follows: LRH-1-5N, 6OQX; LRH-1-6N, 6OQY; LRH-1-2N, 6OR1.

## Table of Contents graphic

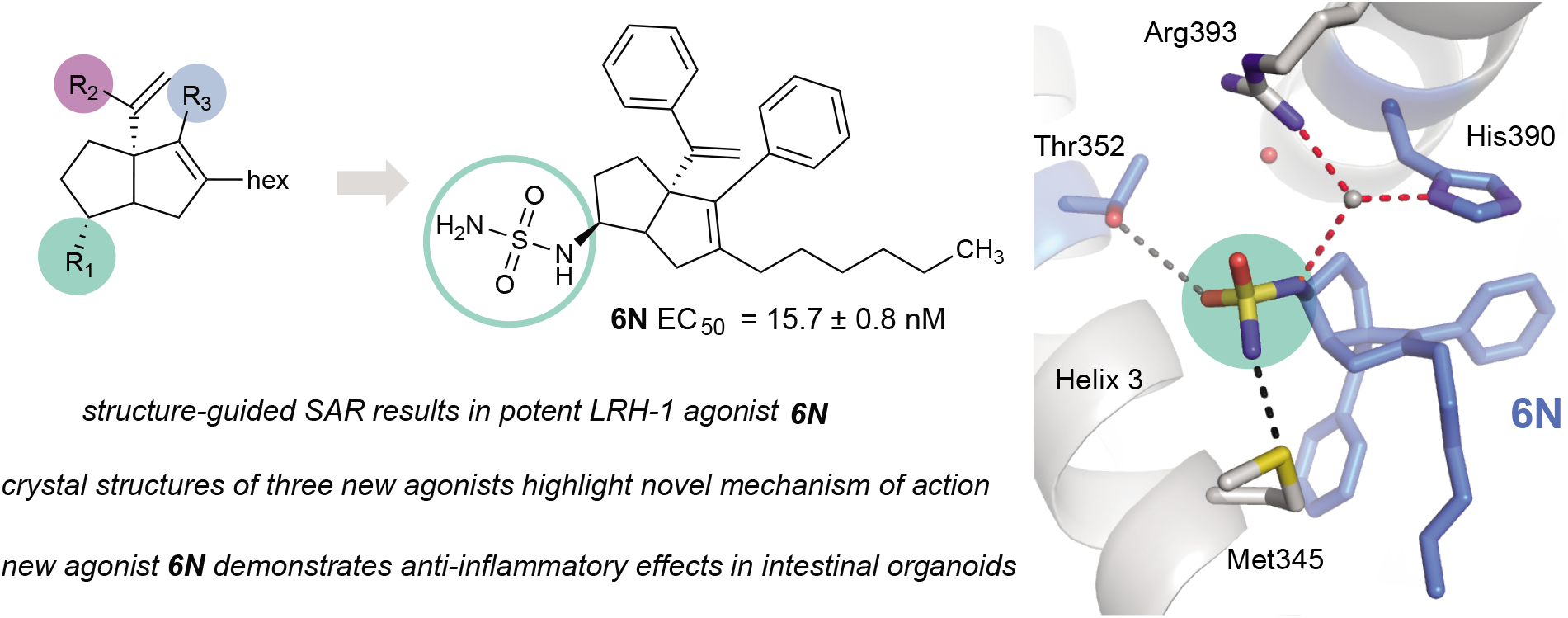

