## Supplemental for "Development of the first low nanomolar Liver Receptor Homolog-1 agonist through structure-guided design"

### Table of Contents

**I.** Additional biological data for tested compounds

a. Supplemental Figure 1: Luciferase curves (**1 – 23**) ............................................................ S1

b. Supplemental Figure 2: HDX peptide coverage and deuterium uptake plots (LRH-1 LBD with **RJW**, **5N**, or **6N**) ............................................................................................S2

c. Supplemental Table 1: X-ray data collection and refinement statistics…………............ S4

**II. General Information**

a. General chemical and synthetic information...................................................................... S6

b. Evaluation of Compound Purity.............................................................................................S7

### III. Detailed Syntheses and Characterization of Tested Compounds 1 – 23

a. R1: Hydroxyl modifications (**1 – 8, S1 – S4**) .......... S8

b. R2: Styrene modifications (**9 – 15**) .......... S19

c. R3: Internal styrene modifications (**16 – 23**) .......... S24

d. NMR spectra of key compounds **2N**, **5N**, and **6N** .......... S42

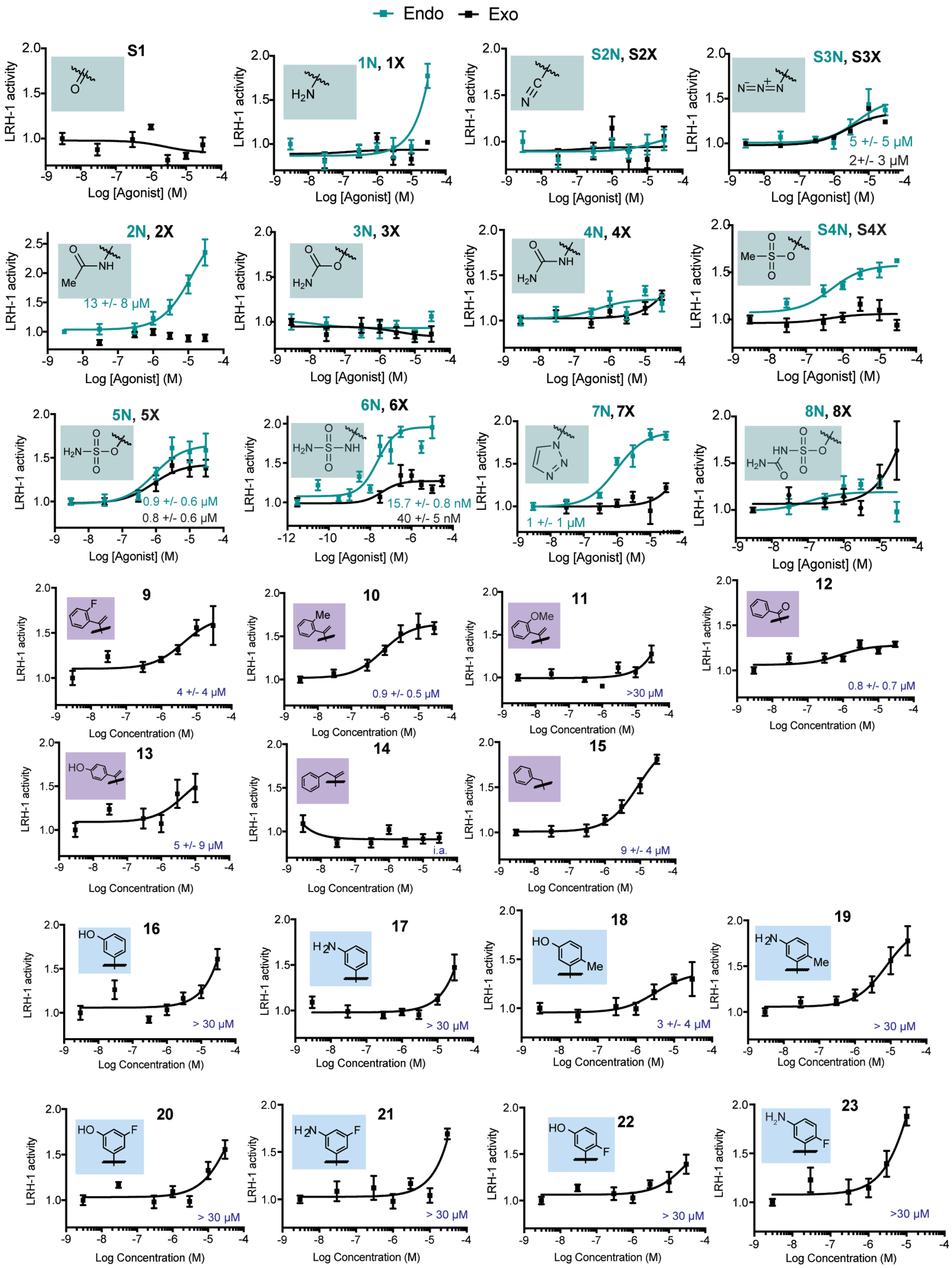


**Supplemental Figure 1.** Dose response curves from luciferase reporter assays in Hela cells. Each point represents the mean ± SEM for three experiments conducted in triplicate.

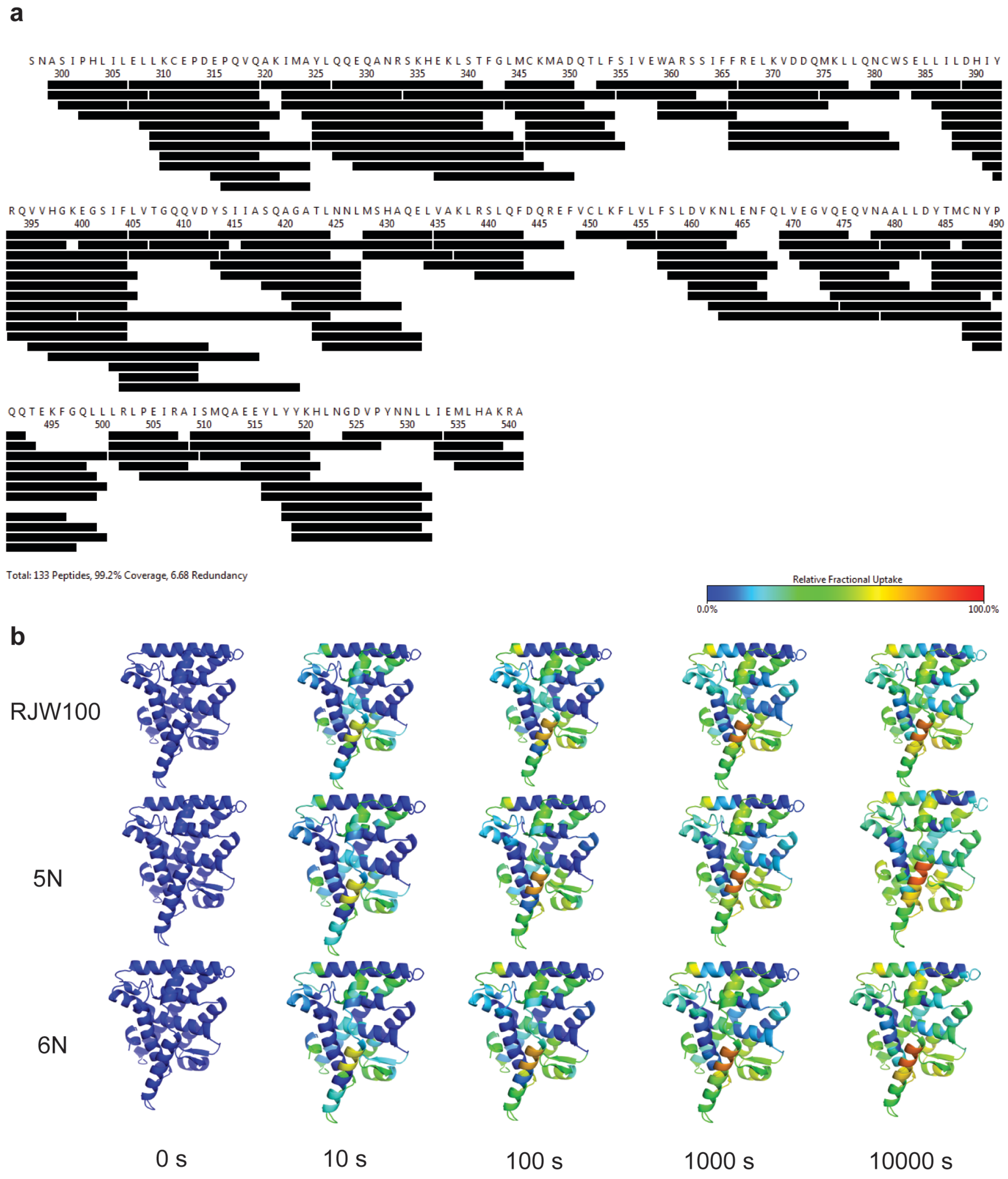


**Supplemental Figure 2.** HDX data for RJW100, 5N, and 6N. A. Map of peptide coverage. B. Fractional uptake of deuterium over time.

**Table S1: X-ray data collection and refinement statistics.**

| **Data collection** | LRH-1 - **5N** -Tif2 | LRH-1 - **6N** - Tif2 | LRH-1 - **2N** -Tif2 |
|---|---|---|---|
| Space group | $P4_32_12$ | $P4_32_12$ | $P4_32_12$ |
| Cell dimensions | | | |
| *a*, *b*, *c* (Å) | 46.5, 46.5, 221.0 | 46.7, 46.7, 218.0 | 46.7, 46.7, 222.7 |
| α,β,γ(°) | 90, 90, 90 | 90, 90, 90 | 90, 90, 90 |
| Resolution (Å) | 50 – 2.00 (2.07-2.00) | 50 – 2.23 (2.31-2.23) | 50 – 2.20 (2.28-2.20) |
| $R_{pim}$ | 0.06 (0.52) | 0.07 (0.46) | 0.04 (0.31) |
| $I$ / σ$I$ | 21.3 (1.72) | 8.9 (3.2) | 18.5 (1.6) |
| $CC_{1/2}$ in highest shell | 0.596 | 0.976 | 0.697 |
| Completeness (%) | 99.9 (100.0) | 97.3 (86.5) | 96.6 (87.9) |
| Redundancy | 11.2 (6.8) | 16.6 (12.5) | 21.1 (13.0) |

| **Refinement** | | | |
|---|---|---|---|
| Resolution (Å) | 2.00 | 2.23 | 2.20 |
| No. reflections | 17346 | 12206 | 13217 |
| $R_{work}$ / $R_{free}$ (%) | 20.6/ 24.5 | 23.2/ 26.9 | 20.0/ 23.9 |
| No. atoms | | | |
| Protein | 4038 | 4098 | 4077 |
| Water | 71 | 24 | 28 |
| Ligand | 68 | 69 | 69 |
| B-factors | | | |
| Protein | 44.8 | 60.5 | 59.3 |
| Ligand | 53.6 | 66.1 | 66.6 |
| Water | 44.0 | 52.7 | 50.7 |
| R.m.s. deviations | | | |

| | | | |
|---|---|---|---|
| Bond lengths (Å) | 0.002 | 0.002 | 0.002 |
| Bond angles (°) | 0.504 | 0.474 | 0.422 |
| Ramachandran favored (%) | 97.6 | 98.0 | 97.1 |
| Ramachandran outliers (%) | 0.4 | 0.0 | 0.0 |
| PDB accession code | 6OQX | 6OQY | 6OR1 |

Values in parenthesis indicate highest resolution shell

### II. a. General information

All reactions were carried out in oven-dried glassware, equipped with a stir bar and under a nitrogen atmosphere with dry solvents under anhydrous conditions, unless otherwise noted. Solvents used in anhydrous reactions were purified by passing over activated alumina and storing under argon. Yields refer to chromatographically and spectroscopically ($^1$H NMR) homogenous materials, unless otherwise stated. Reagents were purchased at the highest commercial quality and used without further purification, unless otherwise stated. n-Butyllithium (n-BuLi) was used as a 1.6 M or a 2.5 M solution in hexanes (Aldrich), was stored at 4°C and titrated prior to use. Organic solutions were concentrated under reduced pressure on a rotary evaporator using a water bath. Chromatographic purification of products was accomplished using forced-flow chromatography on 230-400 mesh silica gel. Preparative thin-layer chromatography (PTLC) separations were carried out on 1000μm SiliCycle silica gel F-254 plates. Thin-layer chromatography (TLC) was performed on 250μm SiliCycle silica gel F-254 plates. Visualization of the developed chromatogram was performed by fluorescence quenching or by staining using $KMnO_4$, p-anisaldehyde, or ninhydrin stains.

$^1$H and $^{13}$C NMR spectra were obtained from the Emory University NMR facility and recorded on a Bruker Avance III HD 600 equipped with cryo-probe (600 MHz), INOVA 600 (600 MHz), INOVA 500 (500 MHz), INOVA 400 (400 MHz), VNMR 400 (400 MHz), or Mercury 300 (300 MHz), and are internally referenced to residual protio solvent signals. Data for $^1$H NMR are reported as follows: chemical shift (ppm), multiplicity (s = singlet, d = doublet, t = triplet, q = quartet, m = multiplet, dd = doublet of doublets, dt = doublet of triplets, ddd= doublet of doublet of doublets, dtd= doublet of triplet of doublets, b = broad, etc.), coupling constant (Hz), integration, and assignment, when applicable. Data for decoupled $^{13}$C NMR are reported in terms of chemical shift and multiplicity when applicable. IR spectra were recorded on a Thermo Fisher Diamond- ATR and reported in terms of frequency of absorption ($cm^{-1}$). High Resolution mass spectra were obtained from the Emory University Mass Spectral facility. Gas Chromatography Mass Spectrometry (GC-MS) was performed on an Agilent 5977A mass spectrometer with an Agilent 7890A gas chromatography inlet. Liquid Chromatography Mass Spectrometry (LC-MS) was performed on an Agilent 6120 mass spectrometer with an Agilent 1220 Infinity liquid chromatography inlet. Preparative High-Pressure Liquid chromatography (Prep-HPLC) was performed on an Agilent 1200 Infinity Series chromatograph using an Agilent Prep-C18 30 x 250 mm 10 μm column, or an Agilent Prep-C18 21.2 x 100 mm, 5 μm column.

### II. b. Evaluation of Compound Purity.

Purity of all tested compounds was determined by HPLC analysis, using the methods given below (as indicated for each compound).

Method A: A linear gradient using water and 0.1 % formic acid (FA) (Solvent A) and MeCN and 0.1% FA (Solvent B); t = 0 min, 30% B, t = 4 min, 99% B (held for 1 min), then 50% B for 1 min, was employed on an Agilent Poroshell 120 EC-C18 2.7 micron, 3.0 mm x 50 mm column (flow rate 1 mL/min) or an Agilent Zorbax SB-C18 1.8 micron, 2.1 mm x 50 mm column (flow rate 0.8 mL/min). The UV detection was set to 254 nm. The LC column was maintained at ambient temperature.

Method B: A linear gradient using water and 0.1 % formic acid (FA) (Solvent A) and MeCN and 0.1% FA (Solvent B); t = 0 min, 70% B, t = 4 min, 99% B (held for 1 min), then 50% B for 1 min, was employed on an Agilent Poroshell 120 EC-C18 2.7 micron, 3.0 mm x 50 mm column (flow rate 1 mL/min) or an Agilent Zorbax SB-C18 1.8 micron, 2.1 mm x 50 mm column (flow rate 0.8 mL/min). The UV detection was set to 254 nm. The LC column was maintained at ambient temperature.

Method C: An isocratic method using 75% MeCN, 35% water, and 0.1 % FA was employed on an Agilent Poroshell 120 EC-C18 2.7 micron, 3.0 mm x 50 mm column (flow rate 1 mL/min) or an Agilent Zorbax SB-C18 1.8 micron, 2.1 mm x 50 mm column (flow rate 0.8 mL/min). The UV detection was set to 254 nm. The LC column was maintained at ambient temperature.

Method D: An isocratic method using 85% MeCN, 15% water, and 0.1% FA was employed on an Agilent Poroshell 120 EC-C18 2.7 micron, 3.0 mm x 50 mm column (flow rate 1 mL/min) or an Agilent Zorbax SB-C C18 1.8 micron, 2.1 mm x 50 mm column (flow rate 0.8 mL/min). The UV detection was set to 254 nm. The LC column was maintained at ambient temperature.

### II. Detailed Syntheses of Tested Compounds 1 – 23

#### a. Hydroxyl modifications 1 – 8

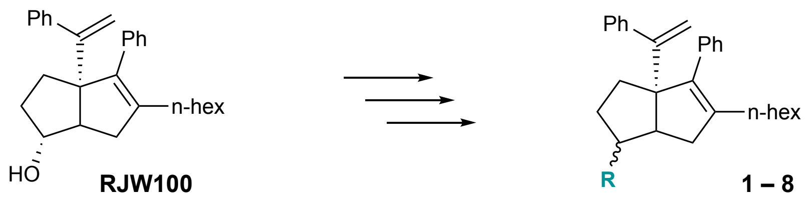


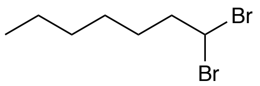


**1,1-dibromoheptane:** Under nitrogen, triphenylphosphite (11.4 mL, 40 mmol 1.1 equiv) was dissolved in DCM and cooled to -78ºC. Bromine (2.0 mL, 40 mmol 1.1 equiv) was added dropwise and stirred briefly. Heptanal (4.2 g, 37 mmol 1.0 equiv.) was then added dropwise in DCM and the reaction was allowed to come to room temperature over 3 hours. The reaction mixture was then filtered through silica and concentrated *in vacuo.* The crude oil was purified by silica gel chromatography in 100% hexanes to afford a clear, colorless oil (6.3 g, 66% yield). Spectral data is consistent with reported values.[1]

**$^1$H NMR** (600 MHz, $CDCl_3$) δ 5.67 (t, *J*=6.2 Hz, 1H), 2.38–2.33 (m, 2H), 1.51 (dd, *J*= 5.9, 3.3 Hz, 2H), 1.35–1.23 (m, 6H), 0.86 (t, *J*=6.9 Hz, 3H).

**RJW100 Synthesis**

Hexahydropentalene formation was accomplished through slight modification of Whitby's procedure.[2] Prior to cyclization, all non-volatile reagents were dried by azeotropic removal of water using benzene. A dry round bottom flask containing bis(cyclopentadienyl)zirconium(IV) dichloride (1.2 equiv) under nitrogen, was dissolved in anhydrous, degassed tetrahydrofuran (THF, 50 mL/mmol enyne) and cooled to -78 °C. The resulting solution was treated with n-BuLi (2.4 equiv.) and the light yellow solution was stirred for 30 minutes. A solution of ***tert*-butyldimethyl((7-phenylhept-1-en-6-yn-3-yl)oxy)silane** (1.0 equiv) in anhydrous, degassed THF (5 mL/mmol) was added. The resulting salmon-colored mixture was stirred at -78 °C for 30

[1] Hoffmann, R. W.; Bovicelli, P. *Synthesis* **1990**, 657-659.

[2] Whitby, R. J.; Stec, J.; Blind, R.D.; Dixon, S.; Leesnitzer, L. M.; Orband-Miller L. A.; Williams S.P.; Willson T.M.; Xu R.; Zuercher W. J.; Cai, F.; Ingraham H. A. *J. Med. Chem.* **2011**, *54* (7), 2266 – 2281.

minutes, the cooling bath removed, and the reaction mixture was allowed to warm to ambient temperature with stirring (2.5 hours total). The reaction mixture was then cooled to -78 °C and the required 1,1-dibromoheptane (1.1 equiv) was added as a solution in anhydrous THF (5 mL/mmol) followed by freshly prepared lithium diisopropylamide (LDA, 1.0 M, 1.1 equiv.). After 15 minutes, a freshly prepared solution of lithium phenylacetylide (3.6 equiv.) in anhydrous THF was added dropwise and the resulting rust-colored solution was stirred at -78 °C for 1.5 hours. The reaction was quenched with methanol and saturated aqueous sodium bicarbonate and allowed to warm to room temperature, affording a light yellow slurry. The slurry was poured onto water and extracted with ethyl acetate four times. The combined organic layers were washed with brine, dried with $MgSO_4$, and concentrated *in vacuo.* The resulting yellow oil was passed through a short plug of silica (20% EtOAc/Hexanes eluent) and concentrated. The crude product was dissolved in THF and treated with solid tetrabutylammonium fluoride hydrate (ca. 2.0 equiv.) and the resulting solution stirred at room temperature for 16 h. The reaction mixture was concentrated and the diastereomers were purified and separated by careful silica gel chromatography (5-20% EtOAc/hexanes eluent) to afford RJW100 *exo* and RJW100 *endo* in a 1.6:1 ratio, respectfully, as determined by characteristic $^1$H NMR signals. The spectral data reported are consistent with literature values.[1] (402.0 mg combined *exo* and *endo*, 58 %).

***Endo:*** **$^1$H NMR:** (400 MHz, $CDCl_3$) δ 7.38 – 7.12 (m, 10H), 5.05 (d, *J* = 1.4 Hz, 1H), 4.92 (d, *J* = 1.4 Hz, 1H), 4.16 (ddd, *J* = 9.0, 8.6, 5.4 Hz, 1H), 2.60 (dd, *J* = 17.3, 2.0 Hz, 1H), 2.46 (td, *J* = 8.7, 2.2 Hz, 1H), 2.13 – 1.95 (m, 3H), 1.84 (ddt, *J* = 10.1, 5.5, 4.6 Hz, 1H), 1.72 – 1.64 (m, 2H), 1.60 – 1.31 (m, 3H), 1.33 – 1.13 (m, 7H), 0.83 (t, *J* = 7.0 Hz, 3H).
***Exo:*** **$^1$H NMR:** (400 MHz, $CDCl_3$) δ 7.37 – 7.14 (m, 10H), 5.05 (d, *J* = 1.4 Hz, 1H), 4.97 (d, *J* = 1.4 Hz, 1H), 3.99 – 3.85 (m, 1H), 2.35 (dd, *J* = 16.6, 9.4 Hz, 1H), 2.27 (d, *J* = 9.4 Hz, 1H), 2.09 – 1.96 (m, 4H), 1.74 – 1.61 (m, 3H), 1.44 – 1.11 (m, 9H), 0.84 (t, *J* = 7.0 Hz, 3H).

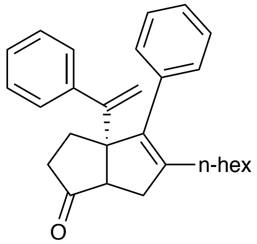


**5-hexyl-4-phenyl-3a-(1-phenylvinyl)-3,3a,6,6a-tetrahydropentalen-1(2H)-one (S1):** A solution of RJW100 (mixture of diastereomers (124.5 mg, 0.3 mmol 1.0 equiv) in acetonitrile was treated with N-methylmorpholine oxide (380.7 mg, 3.2 mmol, 10 equiv) and allowed to stir to homogeneity before the addition of tetrapropylammonium perruthenate (12.6 mg, 0.04 mmol, 0.1 equiv). The solution was stirred at room temperature until completion as determined by TLC (ca. 10 min). The solution was concentrated and subjected directly to silica gel chromatography in 10% EtOAc/Hexanes eluent to afford the title compound as a clear, colorless oil (118.8 mg, 95% yield).
**$^1$H NMR** (400 MHz, $CDCl_3$) δ 7.38 – 7.25 (m, 6H), 7.24 – 7.19 (m, 4H), 5.20 (d, *J* = 1.4 Hz, 1H), 5.09 (d, *J* = 1.4 Hz, 1H), 2.44 (d, *J* = 7.5 Hz, 1H), 2.34 – 2.23 (m, 2H), 2.15 – 1.95 (m, 5H), 1.89 (ddt, *J* = 16.5, 7.8, 1.1 Hz, 1H), 1.29 – 1.09 (m, 8H), 0.82 (t, *J* = 7.0 Hz, 3H).
**$^{13}$C NMR** (101 MHz, $CDCl_3$) δ 222.8, 153.2, 144.9, 142.5, 137.3, 136.6, 129.0, 128.2, 128.1, 127.6, 127.03, 126.96, 115.3, 110.0, 65.4, 55.5, 38.8, 37.5, 31.5, 30.0, 29.4, 28.3, 27.6, 22.5, 14.1.
**LRMS** (ESI, APCI) *m/z*: calc'd for $C_{28}H_{35}O$ $[M+H]^+$ 385.2, found 385.3

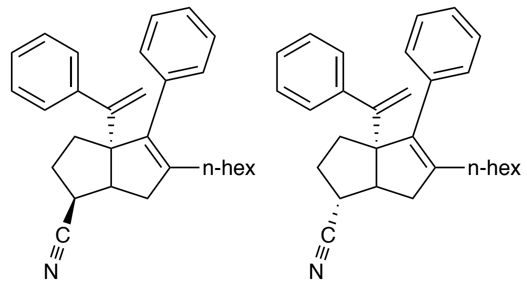


**(*endo* or *exo*)-5-hexyl-4-phenyl-3a-(1-phenylvinyl)-1,2,3,3a,6,6a-hexahydropentalene-1-carbonitrile (S2N, S2X):** A solution of sodium cyanide (10.0 equiv) in DMF was treated with **S4X** (99 mg, 0.2 mmol, 1.0 equiv) or **S4N** (50 mg, 0.1 mmol, 1.0 equiv) as a solution in DMF. The mixture was allowed to stir at 100 °C for about 16 h. The reaction was cooled to ambient temperature, diluted with water, and extracted with EtOAc three times. The combined organic layers were washed with water and brine, dried with $Na_2SO_4$, filtered, and concentrated. The crude oil was purified by silica gel chromatography in 5% EtOAc/Hexanes eluent to afford the title compound. (**S2N** *endo*: 21.9 mg, 26% yield; **S2X** *exo*: 19.9 mg, 47% yield) (Note: An appreciable amount of $E_2$ elimination product is typically also observed, despite optimization of reaction conditions.) (Note: inversion of stereochemistry). (Caution: inorganic cyanides must be handled carefully due to toxicity).

***S2N endo*** **$^1$H NMR** (600 MHz, $CDCl_3$) δ 7.34 – 7.23 (m, 8H), 7.22 – 7.19 (m, 2H), 5.08 (s, 1H), 4.98 (s, 1H), 2.91 – 2.84 (m, 1H), 2.60 (t, $J$ = 9.0 Hz, 1H), 2.53 (d, $J$ = 17.5 Hz, 1H), 2.30 (dd, $J$ = 17.6, 8.6 Hz, 1H), 2.15 – 1.97 (m, 3H), 1.83 – 1.75 (m, 2H), 1.74 – 1.66 (m, 1H), 1.39 (p, $J$ = 7.6 Hz, 2H), 1.31 – 1.17 (m, 6H), 0.84 (d, $J$ = 7.0 Hz, 3H).

***S2N endo*** **$^{13}$C NMR** (126 MHz, $CDCl_3$) δ 153.4, 143.23, 143.22, 137.9, 136.6, 129.6, 127.9, 127.8, 127.7, 127.1, 126.9, 121.1, 115.8, 69.8, 46.5, 39.1, 34.9, 34.7, 31.6, 30.6, 29.8, 29.5, 27.7, 22.6, 14.1.

***S2N endo*** **LRMS** (ESI, APCI) *m/z*: calc'd for $C_{29}H_{35}N$ $[M+H]^+$ 396.6 found 396.4

***S2X exo*** **$^1$H NMR** (600 MHz, $CDCl_3$) δ 7.38 – 7.20 (m, 10H), 5.09 (s, 1H), 5.08 (s, 1H), 2.71 (dd, $J$ = 7.9, 4.9 Hz, 1H), 2.53 (q, $J$ = 11.0, 5.9 Hz, 1H), 2.29 (dd, $J$ = 17.9, 8.4 Hz, 1H), 2.12 – 1.98 (m, 4H), 1.92 – 1.86 (m, 2H), 1.72 (dt, $J$ = 13.1, 5.2 Hz, 1H), 1.36 – 1.15 (m, 8H), 0.84 (t, $J$ = 7.0 Hz, 3H).

***S2X exo*** **$^{13}$C NMR** (75 MHz, $CDCl_3$) δ 153.1, 142.9, 141.5, 138.6, 136.6, 129.4, 128.0, 127.9, 127.8, 126.99, 126.95, 123.1, 115.4, 69.6, 51.8, 41.7, 37.5, 33.8, 31.6, 30.5, 29.7, 29.4, 27.8, 22.6, 14.1.

***S2X exo*** **LRMS** (ESI, APCI) *m/z*: calc'd for $C_{29}H_{36}N$ $[M+H]^+$ 398.3, found 398.3

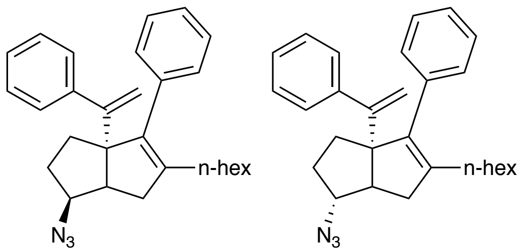


**(*endo* or *exo*)-5-hexyl-4-phenyl-3a-(1-phenylvinyl)-1,2,3,3a,6,6a-hexahydropentalen-1-amine (S3N, S3X):** A solution of **S4X** (139.8 mg, 0.3 mmol, 1.0 equiv) or **S4N** (57 mg, 0.12 mmol, 1.0 equiv) in DMF was treated with sodium azide (10.0 equiv) and the reaction was stirred 16 h at 80 °C behind a blast shield. The solution was allowed to cool to room temperature and poured over water and extracted with EtOAc three times. The combined organic layers were washed with water and brine, dried over $MgSO_4$, and concentrated. The reaction mixture was purified on silica in 0-10% EtOAc/hexanes eluent. (**S3N** *endo*: 117.6 mg, 95% yield; **S3X** *exo*: 45.6 mg, 90% yield) (Note: inversion of stereochemistry). (Warning: caution must be exercised when handling organic and inorganic azides for their toxicity and instability. Aqueous layers were basified and disposed of appropriately).

***S3N endo*** $^{1}$**H NMR** (600 MHz, $CDCl_3$) δ 7.36 – 7.26 (m, 8H), 7.23 – 7.18 (m, 2H), 5.10 (d, *J* = 1.3 Hz, 1H), 4.94 (d, *J* = 1.3 Hz, 1H), 3.87 (ddd, *J* = 10.5, 8.8, 5.9 Hz, 1H), 2.62 – 2.51 (m, 2H), 2.16 – 2.01 (m, 4H), 1.97 – 1.88 (m, 1H), 1.79 (ddd, *J* = 12.4, 5.9, 1.8 Hz, 1H), 1.71 (td, *J* = 12.4, 5.2 Hz, 1H), 1.67 – 1.59 (m, 1H), 1.40 (p, *J* = 7.5 Hz, 2H), 1.31 – 1.19 (m, 5H), 0.87 (t, *J* = 7.2 Hz, 3H).

***S3N endo*** $^{13}$**C NMR** (126 MHz, $CDCl_3$) δ 154.2, 143.8, 143.4, 138.5, 136.8, 129.8, 127.8, 127.7, 126.9, 126.7, 115.5, 69.1, 64.9, 47.9, 35.7, 32.5, 31.7, 30.2, 29.8, 29.5, 27.8, 22.6, 14.1.

***S3N endo*** **LRMS** (ESI) *m/z*: calc'd for $C_{28}H_{34}N_3$ 412.3 $[M+H]^+$, found 411.8

***S3X exo*** $^{1}$**H NMR** (500 MHz, $CDCl_3$) δ 7.36 – 7.24 (m, 8H), 7.21 (dt, *J* = 7.5, 1.4 Hz, 2H), 5.09 (s, 1H), 5.00 (s, 1H), 3.64 (s, 1H), 2.44 – 2.35 (m, 2H), 2.14 – 1.93 (m, 5H), 1.83 – 1.67 (m, 3H), 1.40 – 1.30 (m, 2H), 1.32 – 1.17 (m, 5H), 0.86 (d, *J* = 7.1 Hz, 3H).

***S3X exo*** $^{13}$**C NMR** (126 MHz, $CDCl_3$) δ 153.8, 143.8, 141.3, 139.1, 137.0, 129.6, 127.78, 127.77, 127.72, 126.80, 126.75, 115.3, 71.3, 69.3, 52.1, 41.1, 32.6, 31.6, 31.2, 29.7, 29.4, 27.8, 22.6, 14.1.

***S3X exo*** **LRMS** (ESI) *m/z*: calc'd for $C_{28}H_{34}N_3$ $[M+H]^+$ 412.3, found 412.3

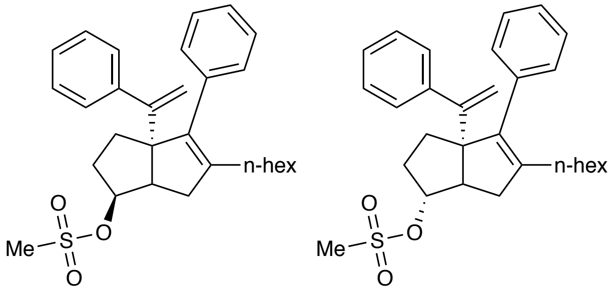


**(*endo* or *exo*) 5-hexyl-4-phenyl-3a-(1-phenylvinyl)-1,2,3,3a,6,6a-hexahydropentalen-1-yl methanesulfonate (S4N, S4X):** A solution of RJW100 *endo* (54.4 mg, 0.14 mmol, 1.0 equiv) or RJW100 *exo* (122.5 mg, 0.3 mmol, 1.0 equiv) in dichloromethane was treated with methanesulfonyl chloride (5.0 equiv), then triethylamine (5.0 equiv) The reaction mixture was allowed to stir 1 h before concentrating and purifying on silica in 30% EtOAc/hexanes eluent. (**S4N** *endo*: 62.1 mg, 95% yield; **S4X** *exo*: 139 mg, >99% yield)

***S4N endo*** $^{1}$**H NMR** (500 MHz, $CDCl_3$) δ 7.36 – 7.26 (m, 8H), 7.24 – 7.17 (m, 2H), 5.13 (d, *J* = 1.1 Hz, 1H), 5.04 – 4.92 (m, 1H), 4.95 (d, *J* = 1.2 Hz, 1H), 3.00 (s, 3H), 2.70 (t, *J* = 9.0, 1.8 Hz, 1H), 2.60 (d, *J* = 17.4 Hz, 1H), 2.17 (dd, *J* = 17.5, 9.1 Hz, 1H), 2.08 (ttd, *J* = 13.5, 6.8, 4.9 Hz, 4H), 1.92 – 1.76 (m, 3H), 1.72 (td, *J* = 12.5, 5.8 Hz, 1H), 1.40 (p, 2H), 1.33 – 1.18 (m, 4H), 0.88 (t, *J* = 7.2 Hz, 3H).
***S4N endo*** $^{13}$**C NMR** (126 MHz, $CDCl_3$) δ 153.7, 143.4, 143.2, 138.5, 136.5, 129.8, 127.9, 127.7, 127.6, 127.0, 126.8, 115.7, 82.9, 68.2, 47.4, 38.2, 34.9, 31.7, 31.1, 31.0, 29.8, 29.4, 27.8, 22.6, 14.1.
***S4N endo*** **LRMS** (ESI, APCI) *m/z*: calc'd for $C_{29}H_{33}$ $[M\text{-}CH_3O_3S]^+$ 369.2, found $[M\text{-}CH_3O_3S]^+$ 368.9.
***S4X exo*** $^{1}$**H NMR** (500 MHz, $CDCl_3$) δ 7.37 – 7.25 (m, 8H), 7.27 – 7.19 (m, 2H), 5.11 (d, *J* = 1.3 Hz, 1H), 5.01 (d, *J* = 1.3 Hz, 1H), 4.83 (d, *J* = 4.0 Hz, 1H), 2.95 (s, 3H), 2.63 (d, *J* = 9.2 Hz, 1H), 2.41 (dd, *J* = 17.4, 9.5 Hz, 1H), 2.14 (dd, *J* = 17.5, 2.0 Hz, 1H), 2.11 – 1.98 (m, 4H), 1.90 – 1.75 (m, 2H), 1.40 – 1.31 (m, 2H), 1.32 – 1.17 (m, 6H), 0.87 (t, *J* = 7.1 Hz, 3H).
***S4X exo*** $^{13}$**C NMR** (151 MHz, $CDCl_3$) δ 153.5, 143.6, 141.3, 138.8, 136.8, 132.8, 129.6, 127.83, 127.76, 127.6, 126.90, 126.86, 115.6, 92.1, 69.2, 53.0, 40.0, 38.7, 32.4, 32.1, 31.6, 29.6, 29.4, 27.8, 22.6, 14.1.
***S4X exo*** **LRMS** (ESI, APCI) *m/z*: calc'd for $C_{29}H_{33}$ $[M\text{-}CH_3O_3S]^+$ 369.2, found $[M\text{-}CH_3O_3S]^+$ 368.9

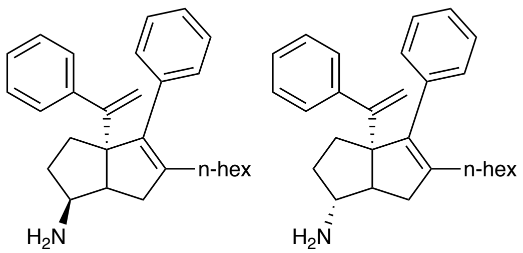


**(*endo* or *exo*)-5-hexyl-4-phenyl-3a-(1-phenylvinyl)-1,2,3,3a,6,6a-hexahydropentalen-1-amine (1N, 1X):** A solution of **S3N** *endo* (54 mg, 0.13 mmol, 1.0 equiv) or **S3X** *exo* (46 mg, 0.1 mmol, 1.0 equiv) in anhydrous $Et_2O$ was cooled to 0 °C and treated dropwise with $LiAlH_4$ (4.0M in $Et_2O$, 10.0 equiv). The reaction was stirred at ambient temperature for ca. 1 h, until the reaction was complete by TLC. The reaction was cooled to 0 °C, diluted with anhydrous $Et_2O$, and slowly treated with water (1mL/g $LiAlH_4$). Excess 4 M NaOH was added slowly and the solution was extracted with EtOAc three times. The combined organic layers were washed with Rochelle's salt and brine, dried over $MgSO_4$, and concentrated. The crude oil was purified by silica gel chromatography in 50% EtOAc/Hexanes eluent (1% triethylamine) to afford the title compounds as colorless oils. (**1N** *endo*: 47.9 mg, 95% yield; **1X** *exo*: 40.0 mg, 92% yield). Purity was established by Method C: *endo* $t_r$ = 0.302 min, 98.6%, *exo* $t_r$ = 0.290 min, 77.5%.
***1N endo*** $^{1}$**H NMR** (600 MHz, $CDCl_3$) δ 7.37 – 7.19 (m, 10H), 5.08 (d, *J* = 1.4 Hz, 1H), 4.94 (d, *J* = 1.5 Hz, 1H), 3.30 (ddd, *J* = 11.0, 8.8, 5.7 Hz, 1H), 2.48 (d, *J* = 17.4 Hz, 1H), 2.42 (t, *J* = 9.0 Hz, 1H), 2.12 – 2.00 (m, 2H), 1.83 – 1.78 (m, 1H), 1.73 – 1.68 (m, 2H), 1.46 – 1.37 (m, 2H), 1.35 – 1.20 (m, 8H), 0.88 (t, *J* = 7.1 Hz, 3H).
***1N endo*** $^{13}$**C NMR** (151 MHz, $CDCl_3$ δ 155.1, 144.2, 142.9, 139.4, 137.2, 129.8, 127.72, 127.66, 127.56, 126.6, 126.5, 115.0, 69.5, 55.3, 49.1, 34.6, 34.1, 33.3, 31.7, 29.9, 29.5, 28.0, 22.6, 14.1.
***1N endo*** **LRMS** (ESI, APCI) *m/z*: calc'd for $C_{28}H_{36}N$ $[M+H]^+$ 386.28, found 385.9

***1X exo*** **$^1$H NMR** (600 MHz, $CDCl_3$) δ 7.33 – 7.28 (m, 4H), 7.28 – 7.21 (m, 6H), 5.04 (d, *J* = 1.5 Hz, 1H), 5.03 (d, *J* = 1.4 Hz, 1H), 3.01 (dt, *J* = 5.4, 3.9 Hz, 1H), 2.32 – 2.26 (m, 1H), 2.08 – 2.02 (m, 4H), 1.79 – 1.72 (m, 1H), 1.65 – 1.60 (m, 1H), 1.42 – 1.17 (m, 10H), 0.85 (t, *J* = 7.2 Hz, 3H).
***1X exo*** **$^{13}$C NMR** (126 MHz, $CDCl_3$) δ 153.8, 143.4, 141.3, 139.0, 137.1, 129.5, 128.1, 127.8, 127.6, 126.71, 126.67, 114.9, 69.2, 61.2, 40.5, 32.1, 31.6, 30.3, 29.7, 29.4, 27.8, 22.6, 14.1.
***1X exo*** **LRMS** (ESI, APCI) *m/z*: calc'd for $C_{28}H_{36}N$ $[M+H]^+$ 386.28, found 385.9

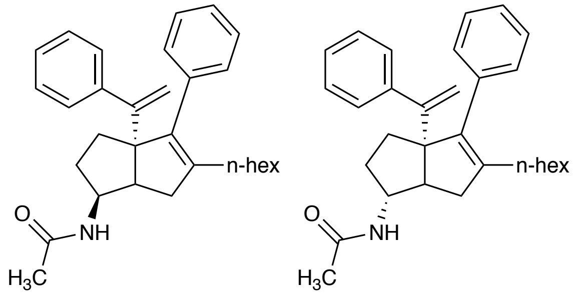


**N-((*endo* or *exo*)-5-hexyl-4-phenyl-3a-(1-phenylvinyl)-1,2,3,3a,6,6a-hexahydropentalen-1-yl)acetamide (2N, 2X):** A solution of **1N** (23 mg, 0.06 mmol, 1.0 equiv) or **1X** (8.4 mg, 0.02 mmol, 1.0 equiv) in DCM was cooled to 0 °C and treated with acetyl chloride (1.5 equiv) and triethylamine (3.0 equiv), then stirred for 1 h. The solution was diluted with water and extracted with DCM three times. The combined organic layers were washed with water and brine, dried with $Na_2SO_4$, filtered, and concentrated. The crude oil was purified on silica gel in 35% EtOAc/Hexanes eluent to afford the title compound as a colorless oil (**2N** *endo:* 21.1 mg, 83% yield; **2X** *exo:* 6.6 mg, 71% yield.). Purity was established by Method D: *endo* $t_R$ = 1.00 min, 96.3 %. *Exo* $t_R$ = 1.04 min, 96.4%.
***2N endo*** **$^1$H NMR** (500 MHz, $CDCl_3$) δ 7.35 – 7.27 (m, 5H), 7.25 – 7.22 (m, 5H), 5.35 (d, *J* = 8.1 Hz, 1H), 5.06 (d, *J* = 1.5 Hz, 1H), 5.02 (d, *J* = 1.5 Hz, 1H), 4.25 (dtd, *J* = 10.5, 8.6, 6.2 Hz, 1H), 2.66 (ddd, *J* = 16.9, 8.4, 1.6 Hz, 1H), 2.14 – 2.00 (m, 4H), 1.99 (s, 3H), 1.87 (dtd, *J* = 11.7, 6.0, 2.3 Hz, 1H), 1.76 (td, *J* = 12.2, 11.7, 5.8 Hz, 1H), 1.66 (ddd, *J* = 12.7, 5.9, 2.3 Hz, 1H), 1.43 – 1.26 (m, 1H), 1.30 – 1.18 (m, 8H), 0.87 (t, *J* = 7.0 Hz, 3H).
***2N endo*** **$^{13}$C NMR** (126 MHz, $CDCl_3$) δ 169.3, 154.6, 143.6, 141.7, 138.9, 137.2, 129.6, 127.9, 127.8, 127.7, 126.9, 126.6, 114.8, 69.0, 59.5, 54.4, 40.9, 33.0, 32.1, 31.6, 29.8, 29.4, 27.8, 26.0, 23.6, 22.6, 14.1.
***2N endo*** **LRMS** (ESI, APCI) *m/z*: calc'd for $C_{30}H_{39}NO$ $[M+H]^+$ 430.3, found 430.3
***2X exo*** **$^1$H NMR** (500 MHz, $CDCl_3$) δ 7.35 – 7.25 (m, 8H), 7.27 – 7.19 (m, 2H), 5.35 (d, *J* = 7.4 Hz, 1H), 5.07 (s, 2H), 3.96 (dd, *J* = 8.2, 4.0 Hz, 1H), 2.34 (dd, *J* = 17.2, 8.6 Hz, 1H), 2.25 (d, *J* = 7.1 Hz, 1H), 2.22 – 2.16 (m, 1H), 2.08 – 2.02 (m, 2H), 1.93 (s, 3H), 1.90 – 1.82 (m, 2H), 1.75 – 1.66 (m, 1H), 1.40 – 1.14 (m, 9H), 0.86 (t, *J* = 7.1 Hz, 3H).
***2X exo*** **$^{13}$C NMR** (126 MHz, $CDCl_3$) δ 169.5, 154.3, 143.5, 143.0, 141.3, 139.3, 136.0, 129.5, 128.0, 127.8, 127.7, 126.7, 115.0, 69.0, 53.1, 47.4, 35.2, 32.1, 31.7, 29.9, 29.6, 28.1, 23.3, 22.6, 14.1.
***2X exo*** **LRMS** (ESI, APCI) *m/z*: calc'd for $C_{30}H_{39}NO$ $[M+H]^+$ 430.3, found 430.3

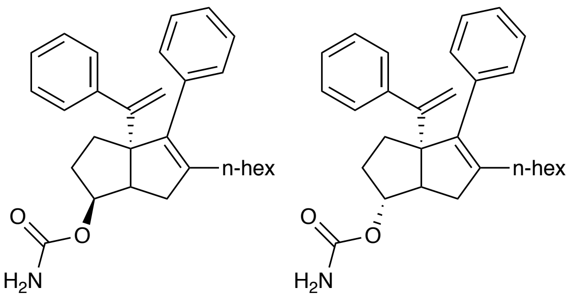

**(*endo* or *exo*)-5-hexyl-4-phenyl-3a-(1-phenylvinyl)-1,2,3,3a,6,6a-hexahydropentalen-1-yl carbamate (3N, 3X):** A solution of RJW100 *endo* (25 mg, 0.07 mmol, 1.0 equiv) or RJW100 *exo* (22 mg, 0.06 mmol, 1.0 equiv) in acetonitrile was cooled to -15 ºC and treated with chlorosulfonyl isocyanate (2.0 equiv) before stirring for 2 h. Concentrated hydrochloric acid (0.5 mL) was added slowly and the reaction was stirred 4 h. The solution was quenched with $NaHCO_3$, diluted with water, and extracted with EtOAc three times. The combined organic layers were rinsed with brine, dried with $Na_2SO_4$, filtered, and concentrated. The crude oil was purified by silica gel chromatography in 5-30% EtOAc/Hexanes eluent to afford the title compounds as yellow oils (**3N** *endo*: 20.4mg, 73% yield. **3X** *exo:* 16.6 mg, 68% yield). Purity was established by Method D: *endo* $t_R$ = 1.20 min, 93.3%. *exo* $t_R$ = 1.26 min, >99%.
***3N endo*** **$^1$H NMR** (400 MHz, $CDCl_3$) δ 7.34 – 7.23 (m, 5H), 7.24 – 7.18 (m, 5H), 5.04 (d, *J* = 1.4 Hz, 1H), 4.93 (d, *J* = 1.4 Hz, 1H), 4.91 – 4.87 (m, 1H), 4.54 (s, 2H), 2.66 (td, *J* = 8.9, 1.8 Hz, 1H), 2.32 (dd, *J* = 17.7, 1.9 Hz, 1H), 2.10 – 1.95 (m, 3H), 1.93 – 1.83 (m, 1H), 1.73 – 1.59 (m, 2H), 1.36 (p, *J* = 7.3 Hz, 2H), 1.29 – 1.16 (m, 7H), 0.84 (t, *J* = 7.0 Hz, 3H).
***3N endo*** **$^{13}$C NMR** (101 MHz, $CDCl_3$) δ 156.4, 154.3, 143.7, 143.3, 138.5, 136.9, 129.7, 127.8, 127.7, 127.6, 126.7, 126.6, 115.2, 68.54 47.0, 34.5, 31.7, 31.1, 30.1, 29.8, 29.4, 27.8, 22.6, 14.1.
***3N endo*** **LRMS** (ESI, APCI) *m/z*: calc'd for $C_{29}H_{39}NO_3$ $[M+H_2O]^-$ 449.3, 449.1
***3X exo*** **$^1$H NMR** (500 MHz, $CDCl_3$) δ 7.41 – 7.19 (m, 10H), 5.08 (d, *J* = 1.6 Hz, 1H), 5.02 (d, *J* = 1.6 Hz, 1H), 4.77 (dt, *J* = 4.2, 1.4 Hz, 1H), 4.56 (s, 2H), 2.42 (d, *J* = 8.1 Hz, 1H), 2.36 (dd, *J* = 16.3, 8.9 Hz, 1H), 2.19 (d, *J* = 17.0 Hz, 1H), 2.14 – 1.88 (m, 4H), 1.85 – 1.63 (m, 3H), 1.41 – 1.29 (m, 2H), 1.31 – 1.16 (m, 5H), 0.88 (t, *J* = 7.0 Hz, 3H).
***3X exo*** **$^{13}$C NMR** (126 MHz, $CDCl_3$) δ 156.6, 154.5, 143.9, 141.9, 138.5, 137.3, 129.6, 127.8, 127.7, 126.8, 126.7, 115.0, 85.6, 69.4, 53.0, 40.3, 32.4, 31.7, 31.5, 29.7, 29.4, 27.8, 22.6, 14.1.
***3X exo*** **LRMS** (ESI, APCI) *m/z*: calc'd for $C_{29}H_{39}NO_3$ $[M+H_2O]^-$ 449.3, found 449.3

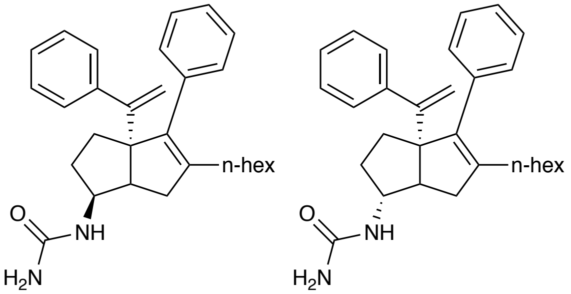

**1-((*endo* or *exo*)-5-hexyl-4-phenyl-3a-(1-phenylvinyl)-1,2,3,3a,6,6a-hexahydropentalen-1-yl)urea (4N, 4X):** A solution of **1N** (15 mg, 0.04 mmol, 1.0 equiv) or **1X** (11 mg, 0.03 mmol, 1.0 equiv) in water was treated with sodium cyanate (10.0 equiv) and 1M aqueous hydrochloric acid

(2.0 equiv). The reaction was heated to 90°C and stirred for approximately 72 hours before being cooled to room temperature, diluted with 3M aqueous NaOH and extracted with $Et_2O$ three times. The combined organic layers were washed with water and brine, dried with $Na_2SO_4$, filtered, and concentrated. The crude oil was purified on silica in 100% EtOAc eluent to afford the title compound as a white solid. (**4N** *endo*: 6.9 mg, 41% yield; **4X** *exo*: 4.5 mg, 37% yield). Purity was established by Method B: *endo* $t_R$ = 2.19 min, 96.5%. *exo* $t_R$ = 4.05 min, >99%.

***4N endo*** **$^1$H NMR** (600 MHz, $CDCl_3$) δ 7.34 – 7.19 (m, 10H), 5.06 (d, *J* = 1.4 Hz, 1H), 4.99 (d, *J* = 1.4 Hz, 1H), 4.48 (d, *J* = 7.7 Hz, 1H), 4.32 (s, 2H), 3.97 (s, 1H), 2.60 (t, *J* = 8.7 Hz, 1H), 2.22 (d, *J* = 17.3 Hz, 1H), 2.09 – 2.03 (m, 2H), 1.93 – 1.85 (m, 1H), 1.73 (td, *J* = 12.5, 5.7 Hz, 1H), 1.67 (ddd, *J* = 12.9, 6.1, 1.9 Hz, 1H), 1.39 – 1.31 (m, 2H), 1.29 – 1.17 (m, 8H), 0.86 (t, *J* = 7.1 Hz, 3H).

***4N endo*** **$^{13}$C NMR** (126 MHz, $CDCl_3$) δ 157.9, 154.4, 143.6, 143.2, 139.0, 136.8, 129.5, 127.9, 127.72, 127.68, 126.8, 126.7, 115.1, 69.2, 47.6, 35.1, 32.1, 31.9, 31.6, 29.9, 29.5, 28.0, 22.6, 14.1.

***4N endo*** **LRMS** (ESI, APCI) *m/z*: calc'd for $C_{29}H_{37}N_2O$ $[M+H]^+$ 429.7, found 428.9

***4X exo*** **$^1$H NMR** (600 MHz, $CDCl_3$) δ 7.34 – 7.19 (m, 10H), 5.06 (d, *J* = 1.3 Hz, 1H), 5.03 (d, *J* = 1.4 Hz, 1H), 4.42 (d, *J* = 7.4 Hz, 1H), 4.25 (s, 2H), 3.70 (s, 1H), 2.36 (dd, *J* = 17.2, 8.9 Hz, 1H), 2.22 – 2.17 (m, 2H), 2.05 (q, *J* = 7.0 Hz, 2H), 1.94 – 1.80 (m, 2H), 1.72 – 1.66 (m, 1H), 1.59 – 1.52 (m, 1H), 1.33 (q, *J* = 7.4 Hz, 2H), 1.28 – 1.17 (m, 6H), 0.85 (t, *J* = 7.2 Hz, 3H).

***4X exo*** **$^{13}$C NMR** (126 MHz, $CDCl_3$) δ 157.7, 154.5, 143.6, 141.5, 139.0, 137.2, 129.6, 127.83, 127.77, 127.7, 126.8, 126.7, 114.9, 69.0, 60.9, 54.4, 41.1, 32.8, 32.4, 31.6, 29.8, 29.7, 29.4, 27.8, 22.6, 14.1.

***4X exo*** **LRMS** (ESI, APCI) *m/z*: calc'd for $C_{29}H_{37}N_2O$ $[M+H]^+$ 429.7, found 428.9

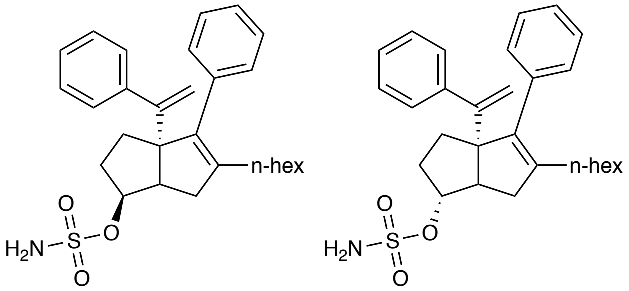


**(*endo* or *exo*)-5-hexyl-4-phenyl-3a-(1-phenylvinyl)-1,2,3,3a,6,6a-hexahydropentalen-1-yl sulfamate (5N, 5X):** A 1M solution of sulfamoyl chloride (2.5 equiv) in DMA was cooled to 0ºC. A solution of the appropriate RJW100 alcohol isomer (*endo* (224.3 mg, 0.6 mmol 1.0 equiv) or *exo* (26 mg, 0.07 mmol, 1.0 equiv)) in DMA was added slowly, followed by triethylamine (excess, ca. 5 equiv); the resulting solution was stirred for one hour. The solution was then diluted with water and extracted with EtOAc three times. The combined organic layers were washed with water and brine, dried with $MgSO_4$, filtered, and concentrated. The crude oil was purified by silica gel chromatography in 20% EtOAc/Hexanes eluent (with 0.5% triethylamine), to afford the title compound as a clear oil (**5N** *endo*: 182 mg, 67% yield. **5X** *exo*: 16.5 mg, 52% yield). Purity was established by Method D: *endo* $t_R$ = 1.15 min, 95.3%. *exo* $t_R$ = 0.97 min, 95.1%.

***5N endo*** **$^1$H NMR** (500 MHz, $CDCl_3$) δ 7.35 – 7.24 (m, 8H), 7.23 – 7.15 (m, 2H), 5.11 (s, 1H), 4.92 (s, 1H), 4.87 (td, *J* = 9.1, 5.2 Hz, 1H), 4.64 (s, 2H), 2.71 (d, *J* = 9.0 Hz, 1H), 2.60 (d, *J* = 17.5 Hz, 1H), 2.17 (dd, *J* = 17.7, 9.3 Hz, 1H), 2.10 – 2.01 (m, 3H), 1.92 – 1.83 (m, 1H), 1.83 – 1.76 (m, 1H), 1.68 (td, *J* = 12.6, 5.6 Hz, 1H), 1.45 – 1.35 (m, 2H), 1.32 – 1.16 (m, 6H), 0.86 (t, *J* = 7.1 Hz, 3H).

***5N endo*** **$^{13}$C NMR** (126 MHz, $CDCl_3$) δ 153.8, 143.5, 143.2, 138.5, 136.5, 129.8, 127.9, 127.7, 127.6, 127.0, 126.8, 115.7, 84.1, 68.2, 47.1, 34.9, 31.6, 31.2, 30.5, 29.8, 29.4, 27.7, 22.6, 14.1.
***5N endo*** **LRMS** (ESI, APCI) *m/z*: calc'd for $C_{28}H_{36}NO_3S$ $[M-H]^-$ 465.3, found 465.4
***5X exo*** **$^{1}$H NMR** (500 MHz, $CDCl_3$) δ 7.36 – 7.28 (m, 7H), 7.29 – 7.16 (m, 3H), 5.10 (d, *J* = 1.3 Hz, 1H), 5.00 (d, *J* = 1.3 Hz, 1H), 4.75 (d, *J* = 4.4 Hz, 1H), 4.62 (s, 2H), 2.68 (d, *J* = 9.1 Hz, 1H), 2.40 (dd, *J* = 18.1, 9.4 Hz, 1H), 2.19 – 2.01 (m, 6H), 1.88 – 1.73 (m, 2H), 1.38 – 1.16 (m, 7H), 0.87 (t, *J* = 7.1 Hz, 3H).
***5X exo*** **$^{13}$C NMR** (126 MHz, $CDCl_3$) δ 153.6, 143.7, 141.4, 138.8, 136.8, 129.6, 127.8, 127.8, 127.7, 126.89, 126.86, 115.6, 93.7, 69.3, 52.8, 40.1, 32.1, 32.0, 31.6, 29.7, 29.4, 27.8, 22.6, 14.1.
***5X exo*** **LRMS** (ESI, APCI) *m/z*: calc'd for $C_{28}H_{36}NO_3S$ $[M-H]^-$ 465.3, found 465.2

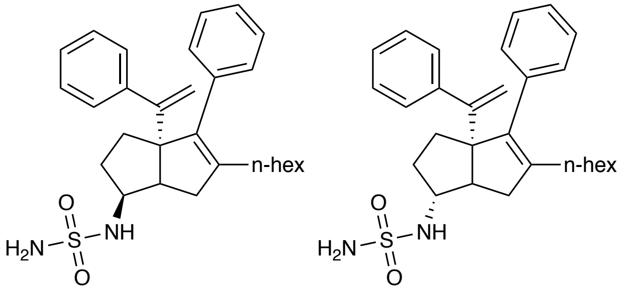


**(*endo* or *exo*)-5-hexyl-4-phenyl-3a-(1-phenylvinyl)-1,2,3,3a,6,6a-hexahydropentalen-1-yl sulfamide (6N, 6X):** A solution of **1N** (30 mg, 0.08 mmol, 1.1 equiv) or **1X** (12 mg, 0.03 mmol, 1.1 equiv) in DCM was treated with triethylamine (2.0 equiv.) and solution of 2-oxo-1,3-oxazolidine-3-sulfonyl chloride (0.5 M in DCM, 1.0 equiv) (prepared according to the procedure of Borghese et al.).[2] The reaction was stirred at room temperature for 3 h then concentrated. The residue was treated with ammonia (0.5 M in dioxane, 1.5 equiv) and triethylamine (3.0 equiv). The solution was heated in a sealed tube at 85°C for 16 h behind a blast shield. After cooling to ambient temperature, the reaction was diluted with 3:3:94 MeOH:$Et_3N$:EtOAc and passed through a pad of silica. The eluent was concentrated, and the crude oil was purified on silica in 20-30% EtOAc/hexanes eluent to afford the title compound as a colorless oil. (**6N** *endo*: 21.6 mg, 60% yield; **6X** *exo*: 5.4 mg, 36% yield) Purity was established by Method C: *endo* $t_R$ = 2.0 min, 96.6%. *exo* $t_R$ = 1.89 min, 79.6%.
***6N endo*** **$^{1}$H NMR** (600 MHz, $CDCl_3$) δ 7.33 – 7.23 (m, 8H), 7.20 – 7.17 (m, 2H), 5.09 (d, *J* = 1.3 Hz, 1H), 4.96 (d, *J* = 1.3 Hz, 1H), 4.44 (s, 2H), 4.36 (d, *J* = 8.0 Hz, 1H), 3.84 – 3.77 (m, 1H), 2.62 (td, *J* = 8.9, 2.0 Hz, 1H), 2.38 (dd, *J* = 17.5, 2.0 Hz, 1H), 2.20 – 2.13 (m, 1H), 2.08 – 2.04 (m, 2H), 2.00 – 1.95 (m, 1H), 1.74 – 1.70 (m, 2H), 1.50 – 1.43 (m, 1H), 1.42 – 1.16 (m, 8H), 0.86 (t, *J* = 7.1 Hz, 3H).
***6N endo*** **$^{13}$C NMR** (126 MHz, $CDCl_3$) δ 154.1, 143.6, 142.8, 139.3, 136.6, 129.6, 127.8, 127.7, 126.9, 126.8, 115.5, 68.8, 57.2, 47.4, 35.4, 32.3, 32.0, 31.6, 29.8, 29.5, 27.9, 22.6, 14.1.
***6N endo*** **LRMS** (ESI, APCI) *m/z*: calc'd for $C_{28}H_{37}N_2O_2S$ $[M+H]^+$ 465.7, found 464.8

[2] Borghese, A.; Antoine, L.; Van Hoeck, J. P.; Mockel, A.; Merschaert, A. *Org. Process Res. Dev.* **2006**, *10*, 770.

***6X exo*** $^{1}$**H NMR** (600 MHz, $CDCl_3$) δ 7.34 – 7.24 (m, 8H), 7.23 – 7.18 (m, 2H), 5.08 (d, *J* = 1.2 Hz, 1H), 5.01 (d, *J* = 1.2 Hz, 1H), 4.38 (s, 2H), 4.21 (d, *J* = 7.2 Hz, 1H), 3.58 – 3.52 (m, 1H), 2.40 (dd, *J* = 16.9, 8.9 Hz, 1H), 2.36 – 2.31 (m, 1H), 2.18 (d, *J* = 16.9 Hz, 1H), 2.05 (td, *J* = 7.5, 2.6 Hz, 2H), 1.99 – 1.85 (m, 2H), 1.76 – 1.69 (m, 2H), 1.38 – 1.30 (m, 1H), 1.30 – 1.15 (m, 7H), 0.86 (t, *J* = 7.2 Hz, 3H).
***6X exo*** $^{13}$**C NMR** (126 MHz, $CDCl_3$) δ 154.1, 143.5, 141.2, 139.3, 136.9, 129.6, 127.9, 127.7, 126.9, 126.8, 115.2, 68.8, 63.8, 54.0, 40.8, 32.8, 32.3, 31.6, 29.71, 29.69, 29.4, 27.8, 22.6, 14.1.
***6X exo*** **LRMS** (ESI, APCI) *m/z*: calc'd for $C_{29}H_{37}N_2O$ $[M+H]^+$ 465.7, found 464.8

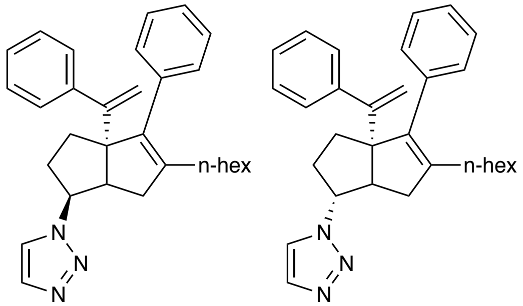


***(endo*** **or** ***exo)*** **5-hexyl-4-phenyl-3a-(1-phenylvinyl)-1,2,3,3a,6,6a-hexahydropentalen-1-yl)-1H-1,2,3-triazole (7N, 7X):** A solution of ascorbic acid (1.0 equiv), and potassium carbonate (6.0 equiv) in water was treated with copper sulfate pentahydrate (1.0 equiv) and stirred briefly. Trimethysilyl acetylene (6.0 equiv.) was added in MeOH before addition of **SN *endo*** (29 mg, 0.07 mmol, 1.0 equiv) or **S3X *exo*** (29 mg, 0.07 mmol, 1.0 equiv) in MeOH. The reaction mixture was stirred 16 h, diluted with water, and extracted with EtOAc three times. The combined organics were washed with brine and dried over $MgSO_4$ before concentration. The crude oil was purified by silica gel chromatography in 30% EtOAc/Hexanes eluent to afford the title compounds (**7N** *endo*: 19.0 mg, 62% yield, **7X** *exo*: 14.7 mg, 48% yield). Purity was established by: *endo* (Method B) $t_R$ = 2.88 min, 97.5%. *exo* (Method D) $t_R$ = 1.46 min, 78.0%.
***7N endo*** $^{1}$**H NMR** (600 MHz, $CDCl_3$) δ 7.70 (s, 1H), 7.49 (s, 1H), 7.36 – 7.25 (m, 8H), 7.24 – 7.21 (m, 2H), 5.14 (s, 1H), 5.00 (s, 1H), 4.96 (ddd, *J* = 11.5, 9.5, 6.7 Hz, 1H), 2.94 (td, *J* = 9.2, 1.9 Hz, 1H), 2.29 – 2.20 (m, 2H), 2.03 – 1.84 (m, 5H), 1.27 – 1.09 (m, 9H), 0.81 (t, *J* = 7.0 Hz, 3H).
***7N endo*** $^{13}$**C NMR** (126 MHz, $CDCl_3$) δ 153.8, 143.5, 138.4, 133.3, 129.7, 127.9, 127.8, 127.7, 127.0, 126.9, 122.9, 115.9, 69.1, 63.4, 48.6, 35.7, 32.6, 31.5, 29.7, 29.4, 29.3, 27.7, 22.6, 14.1.
***7N endo*** **LRMS** (ESI, APCI) *m/z*: calc'd for $C_{30}H_{38}N_3$ $[M+H]^+$ 440.3, found 440.4
***7X exo*** $^{1}$**H NMR** (600 MHz, $CDCl_3$) δ 7.66 (s, 1H), 7.43 (s, 1H), 7.37 – 7.20 (m, 10H), 5.13 (dd, *J* = 0.9 Hz, 1H), 5.09 (d, *J* = 0.9 Hz, 1H), 4.79 – 4.72 (m, 1H), 2.79 (dd, *J* = 8.6, 3.8 Hz, 1H), 2.41 (dd, *J* = 17.2, 8.6 Hz, 1H), 2.28 (d, *J* = 17.5 Hz, 1H), 2.18 – 1.98 (m, 4H), 1.85 – 1.77 (m, 1H), 1.40 – 1.33 (m, 2H), 1.29 – 1.16 (m, 7H), 0.85 (t, *J* = 7.1 Hz, 3H).
***7X exo*** $^{13}$**C NMR** (101 MHz, $CDCl_3$) δ 153.4, 143.2, 141.3, 139.2, 136.7, 129.6, 128.0, 127.0, 127.8, 127.1, 126.9, 121.7, 115.7, 110.0, 70.0, 69.3, 53.1, 41.2, 33.0, 32.9, 31.6, 29.8, 29.4, 27.8, 22.6, 14.1.
***7X exo*** **LRMS** (ESI, APCI) *m/z*: calc'd for $C_{30}H_{38}N_3$ $[M+H]^+$ 440.3, found 440.3

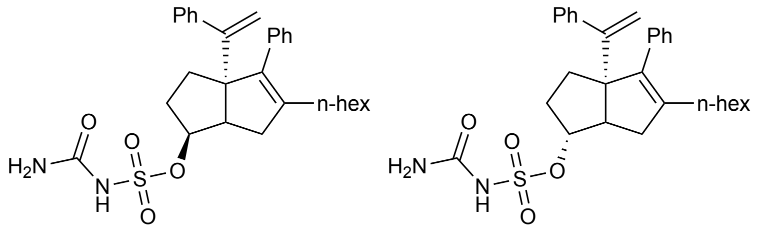

***(endo* or *exo)*** **5-hexyl-4-phenyl-3a-(1-phenylvinyl)-1,2,3,3a,6,6a-hexahydropentalen-1-yl carbamoylsulfamate (8N, 8X):** To a solution of sodium hydride (60% suspension in mineral oil, 2.0 equiv) at 0 °C was added either **5X** (49.7 mg, 0.11 mmol, 1.0 equiv) or **5N** (33.6 mg, 0.07 mmol, 1.0 equiv) in THF at 0 °C for 1 h and allowed to warm to room temperature. A solution of carbonyldiimidzaole (1.5 equiv) in THF was added to the reaction mixture at room temperature. The resulting solution was stirred for 1 h before slow addition of excess ammonia in methanol (7N). The solution was allowed to stir for an additional 3 h. The crude reaction mixture was dissolved in EtOAc, washed with brine, and dried in $MgSO_4$. The organic layer was concentrated, and the crude mixture was purified via preparatory HPLC (**8N** *endo*: 6.1 mg, 16% yield, **8X** *exo*: 2.5 mg, 5% yield). Purity was established by Method D: *endo* $t_R$ = 1.74 min, 96.2%. *exo* $t_R$ = 1.74 min, >99%.

***8N endo*** **$^1$H NMR** (400 MHz, $CDCl_3$) δ 7.35 (s, 1H), 7.33 – 7.20 (m, 8H), 7.19 – 7.13 (m, 2H), 5.08 (d, *J* = 1.2 Hz, 1H), 5.02 (td, *J* = 9.1, 6.0 Hz, 1H), 4.91 (d, *J* = 1.4 Hz, 1H), 3.74 (s, 2H), 2.67 (td, *J* = 9.0, 2.3 Hz, 1H), 2.58 (dd, *J* = 17.5, 2.1 Hz, 1H), 2.17 – 1.99 (m, 3H), 1.91 – 1.62 (m, 3H), 1.40 – 1.27 (m, 2H), 1.27 – 1.15 (m, 7H), 0.83 (t, *J* = 7.0 Hz, 3H).

***8N endo*** **LRMS** (ESI, APCI) *m/z*: calc'd for $C_{28}H_{33}$ $[M-CH_3N_2O_4S]^+$ 369.6, found 369.2. calc'd for $C_{29}H_{35}N_2O_4S$ $[M-H]^-$ 507.7, found 507.2.

***8X exo*** **$^1$H NMR** (400 MHz, $CDCl_3$) δ 7.32 – 7.11 (m, 11H), 4.99 (s, 1H), 4.93 (s, 1H), 4.69 (s, 1H), 3.60 – 3.48 (m, 2H), 2.60 (d, *J* = 9.1 Hz, 1H), 2.25 (dd, *J* = 17.4, 9.2 Hz, 1H), 2.11 – 1.90 (m, 3H), 1.75 – 1.54 (m, 2H), 1.32 – 1.07 (m, 10H), 0.81 (t, *J* = 7.1 Hz, 3H).

***8X exo*** **LRMS** (ESI, APCI) *m/z*: calc'd for $C_{28}H_{33}$ $[M-CH_3N_2O_4S]^+$ 369.6, found 369.2. calc'd for $C_{29}H_{35}N_2O_4S$ $[M-H]^-$ 507.7, found 507.2.

#### b. R2: Styrene Modifications 9 – 15

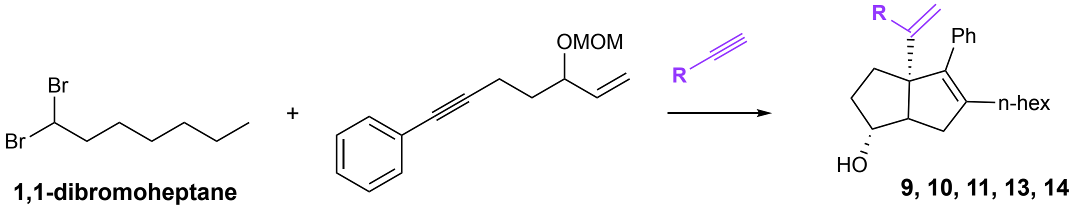


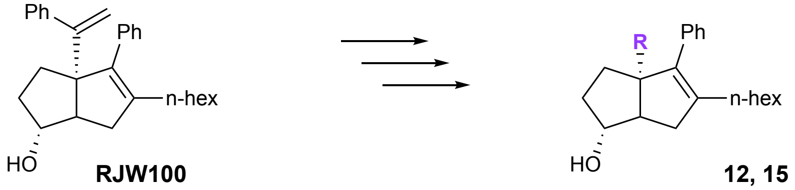


Hexahydropentalene formation was accomplished through slight modification of Whitby's procedure.[3] Prior to cyclization, all non-volatile reagents were dried by azeotropic removal of water using benzene. A dry round bottom flask containing bis(cyclopentadienyl)zirconium(IV) dichloride (1.2 equiv) under nitrogen, was dissolved in anhydrous, degassed tetrahydrofuran (THF, 50 mL/mmol enyne) and cooled to -78 °C. The resulting solution was treated with n-BuLi (2.4 equiv.) and the light yellow solution was stirred for 30 minutes. A solution of (5-(methoxymethoxy)hept-6-en-1-yn-1-yl)benzene (1.0 equiv) (prepared according to a literature procedure)[4] in anhydrous, degassed THF (5 mL/mmol) was added. The resulting salmon-colored mixture was stirred at -78 °C for 30 minutes, the cooling bath removed, and the reaction mixture was allowed to warm to ambient temperature with stirring (2.5 hours total). The reaction mixture was then cooled to -78 °C and the required 1,1-dibromoheptane (1.1 equiv) was added as a solution in anhydrous THF (5 mL/mmol) followed by freshly prepared lithium diisopropylamide (LDA, 1.0 M, 1.1 equiv.). After 15 minutes, a freshly prepared solution of lithium phenylacetylide (3.6 equiv.) in anhydrous THF was added dropwise and the resulting rust-colored solution was stirred at -78 °C for 1.5 hours. The reaction was quenched with methanol and saturated aqueous sodium bicarbonate and allowed to warm to room temperature, affording a light yellow slurry. The slurry was poured onto water and extracted with ethyl acetate four times. The combined organic layers were washed with brine, dried with $MgSO_4$, and concentrated *in vacuo.* The resulting yellow oil was passed through a short plug of silica (20% EtOAc/Hexanes eluent) and concentrated. The crude product was dissolved in acetonitrile and treated with concentrated aqueous hydrochloric acid (ca 5 equiv) and the resulting solution stirred at room temperature until completion of the reaction was detected (typically fewer than 10 minutes). The reaction mixture was concentrated and purified by silica gel chromatography (20% EtOAc/hexanes eluent) to afford the title compounds **9 – 15** as a 7:1 mixture of diastereomers, favoring the desired *exo*-isomer.

---

[3] Flynn, Autumn R.; Mays, Suzanne G.; Ortlund, Eric A.; and Jui, Nathan T. *ACS Med. Chem. Lett.* **2018** *9*(10), 1051–1056.

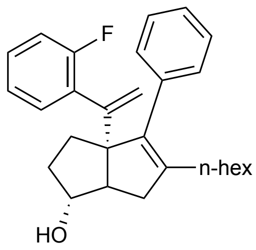


**(*exo)*-3a-(1-(2-fluorophenyl)vinyl)-5-hexyl-4-phenyl-1,2,3,3a,6,6a-hexahydropentalen-1-ol (9):** According to the general procedure, (5-(methoxymethoxy)hept-6-en-1-yn-1-yl)benzene (179.7 mg, 0.8 mmol) was reacted with 1-ethynyl-2-fluorobenzene (320 µL, 2.8 mmol). The crude oil was purified in 5-20% EtOAc/hexanes eluent to give the title compound (35.5 mg, 11% yield over two steps). Purity was established as the *exo* diastereomer by Method D: $t_R$ = 1.44 min, 98.3%.

**$^1$H NMR** (500 MHz, $CDCl_3$) δ 7.38 – 7.17 (m, 7H), 7.03 (dtt, *J* = 13.0, 7.5, 1.1 Hz, 2H), 5.27 (s, 1H), 5.11 (s, 1H), 3.92 (d, *J* = 4.3 Hz, 1H), 2.31 (d, *J* = 14.3 Hz, 2H), 2.15 – 1.95 (m, 4H), 1.76 – 1.57 (m, 4H), 1.40 – 1.15 (m, 7H), 0.86 (t, *J* = 7.1, 3H).

**$^{13}$C NMR** (126 MHz, $CDCl_3$) δ 160.6, 158.6, 147.9, 141.8, 138.3, 137.4, 130.4, 129.7, 128.4, 128.3, 127.7, 126.6, 123.2, 116.7, 115.4, 115.2, 82.0, 69.4, 56.8, 39.7, 34.7, 31.7, 31.2, 29.8, 29.4, 27.9, 22.6, 14.1.

**$^{19}$F NMR** (282 MHz, $CDCl_3$) δ -113.74.

**LRMS** (ESI, APCI) *m/z*: calc'd for $C_{28}H_{36}FO$ $[M+H]^+$ 405.3, found 405.9

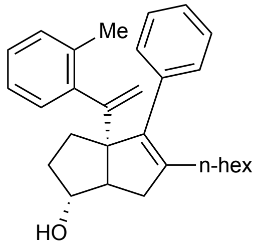


**(*exo)*-5-hexyl-4-phenyl-3a-(1-(o-tolyl)vinyl)-1,2,3,3a,6,6a-hexahydropentalen-1-ol (10):** According to the general procedure, (5-(methoxymethoxy)hept-6-en-1-yn-1-yl)benzene **(**144.4 mg, 0.6 mmol) was reacted with 1-ethynyl-2-methylbenzene (280 µL, 2.2 mmol). The crude oil was purified in 10% EtOAc/hexanes eluent to give the title compound (62.9 mg, 25% over two steps). Purity was established as the *exo* diastereomer by Method B: $t_R$ = 3.66 min, 98.3%.

**$^1$H NMR** (600 MHz, $CDCl_3$) δ 7.36 – 7.30 (m, 2H), 7.30 – 7.26 (m, 1H), 7.24 (d, *J* = 7.5 Hz, 2H), 7.18 (d, *J* = 7.5 Hz, 1H), 7.14 (t, *J* = 7.4 Hz, 1H), 7.04 (t, *J* = 7.4 Hz, 1H), 5.08 (s, 1H), 4.95 (s, 1H), 3.91 (s, 1H), 2.26 (s, 3H), 2.25 – 2.12 (m, 2H), 2.06 – 1.88 (m, 4H), 1.76 – 1.66 (m, 2H), 1.61 (dd, *J* = 11.9, 6.3 Hz, 1H), 1.42 – 1.28 (m, 3H), 1.28 – 1.22 (m, 1H), 1.22 – 1.15 (m, 4H), 0.85 (t, *J* = 7.2 Hz, 3H).

**$^{13}$C NMR** (126 MHz, $CDCl_3$) δ 152.3, 142.9, 141.7, 138.5, 137.7, 135.6, 130.1, 130.0, 127.6, 126.6, 126.5, 124.8, 115.5, 82.1, 74.7, 70.0, 55.7, 39.8, 34.6, 31.8, 31.7, 29.7, 29.3, 27.9, 22.6, 20.7, 14.1.

**LRMS** (ESI, APCI) *m/z*: calc'd for $C_{29}H_{39}O$ $[M+H]^+$ 401.3, found 401.0

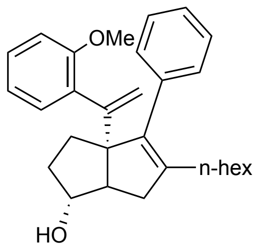


**(*exo*)-5-hexyl-3a-(1-(2-methoxyphenyl)vinyl)-4-phenyl-1,2,3,3a,6,6a-hexahydropentalen-1-ol (11):** According to the general procedure, (5-(methoxymethoxy)hept-6-en-1-yn-1-yl)benzene (146.5 mg, 0.6 mmol) was reacted with 1-ethynyl-2-methoxybenzene (300 µL, 2.3 mmol). The crude oil was purified in 10-20% EtOAc/hexanes eluent to give the title compound (12.2 mg, 5% yield over two steps). Purity was established as the *exo* diastereomer by Method B: $t_R$ = 2.95 min, 97.6%

**$^1$H NMR** (600 MHz, $CDCl_3$) δ 7.36 – 7.18 (m, 6H), 6.95 (d, *J* = 7.1 Hz, 1H), 6.88 – 6.81 (m, 2H), 5.18 (d, *J* = 1.8 Hz, 1H), 5.01 (d, *J* = 1.8 Hz, 1H), 3.86 (s, 1H), 3.75 (s, 3H), 2.50 (dd, *J* = 16.6, 9.1 Hz, 1H), 2.45 (d, *J* = 8.6 Hz, 1H), 2.08 (d, *J* = 16.6 Hz, 1H), 2.01 (t, *J* = 7.7 Hz, 2H), 1.81 – 1.72 (m, 2H), 1.65 – 1.58 (m, 2H), 1.58 – 1.53 (m, 1H), 1.36 – 1.29 (m, 2H), 1.27 – 1.20 (m, 2H), 1.21 – 1.14 (m, 2H), 0.84 (t, *J* = 7.2 Hz, 3H).

**$^{13}$C NMR** (151 MHz, $CDCl_3$) δ 172.5, 169.6, 156.4, 151.5, 140.5, 137.7, 132.6, 130.1, 129.8, 128.0, 127.5, 126.4, 120.4, 115.6, 110.8, 81.7, 69.3, 57.6, 55.6, 51.3, 39.6, 34.6, 31.6, 31.1, 29.6, 29.2, 28.0, 22.6, 14.1.

**LRMS** (ESI, APCI) *m/z*: calc'd for $C_{29}H_{39}O_2$ $[M+H]^+$ 417.3, found 417.9

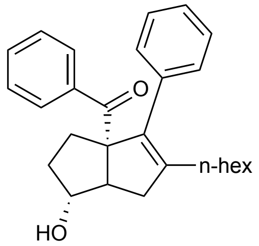


**(*exo*)-5-hexyl-1-hydroxy-4-phenyl-2,3,6,6a-tetrahydropentalen-3a(1*H*)-yl)(phenyl)methanone (12):**

A solution of RJW100 (11.2 mg, 0.03 mmol) in DCM was cooled to -78°C and treated with ozone until the solution was blue. At this point, the stream of ozone was stopped and the reaction was stirred until the blue color dissipated. DMS was added (11 µL, 0.15 mmol, 5.0 equiv) and briefly stirred. The reaction solution was concentrated, and the crude reaction mixture was purified on silica in 0-20% EtOAc/Hex to afford a clear, colorless oil (9.4 mg, 81% yield). Purity was established by Method D: $t_R$ = 1.43 min, 96.5%.

**$^1$H NMR** (600 MHz, $CDCl_3$) δ 7.86 (d, *J* = 7.7 Hz, 2H), 7.46 (t, *J* = 7.4 Hz, 1H), 7.35 (dd, *J* = 8.7, 6.7 Hz, 2H), 7.18 (d, *J* = 6.6 Hz, 3H), 6.87 (dd, *J* = 7.2, 2.3 Hz, 2H), 4.06 (s, 1H), 3.02 (dd, *J* = 17.5, 10.2 Hz, 1H), 2.90 (d, *J* = 11.2 Hz, 1H), 2.76 – 2.64 (m, 1H), 2.33 (dd, *J* = 17.7, 3.5 Hz, 1H), 2.09 (t, *J* = 7.8 Hz, 2H), 2.00 (d, *J* = 12.2 Hz, 1H), 1.85 – 1.66 (m, 2H), 1.49 – 1.36 (m, 2H), 1.29 – 1.15 (m, 6H), 0.85 (dd, *J* = 14.0, 7.2 Hz, 3H).

**$^{13}$C NMR** (101 MHz, $CDCl_3$) δ 203.4, 142.0, 140.0, 138.6, 136.2, 131.8, 129.0, 128.6, 128.1, 128.1, 127.0, 80.8, 76.3, 54.6, 40.5, 32.7, 31.6, 30.5, 29.4, 29.3, 27.8, 22.6, 21.6, 14.1.

**LRMS** (ESI, APCI) *m/z*: calc'd for $C_{27}H_{35}O_2$ $[M+H]^+$ 389.6, found 389.2

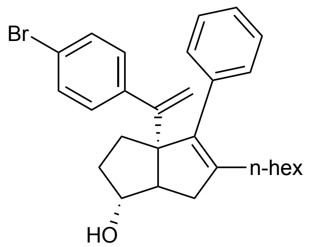


**(*exo)*-3a-(1-(4-bromophenyl)vinyl)-5-hexyl-4-phenyl-1,2,3,3a,6,6a-hexahydropentalen-1-ol (S9):** According to the general procedure, (5- (methoxymethoxy)hept-6-en-1-yn-1-yl)benzene (273.6 mg, 1.2 mmol) was reacted with 1-bromo-4-ethynylbenzene (776.3 mg, 4.3 mmol). The crude oil was purified in 10% EtOAc/hexanes eluent to give the title compound (105.0 mg, 19% yield over two steps).

**$^1$H NMR** (500 MHz, $CDCl_3$) δ 7.38 (d, *J* = 8.1 Hz, 2H), 7.34 – 7.27 (m, 3H), 7.23 (d, *J* = 8.1 Hz, 2H), 7.16 (d, *J* = 8.6 Hz, 2H), 5.06 (s, 1H), 5.01 (s, 1H), 3.96 (s, 1H), 2.38 (dd, *J* = 17.2, 9.4 Hz, 1H), 2.26 (d, *J* = 9.4 Hz, 1H), 2.13 – 1.98 (m, 4H), 1.72 – 1.65 (m, 3H), 1.33 (d, *J* = 7.5 Hz, 3H), 1.29 – 1.18 (m, 5H), 0.87 (t, *J* = 7.1 Hz, 3H).

**$^{13}$C NMR** (126 MHz, $CDCl_3$) δ 153.6, 143.1, 141.5, 138.9, 137.2, 130.8, 129.6, 129.4, 127.7, 126.7, 120.8, 115.5, 82.0, 69.2, 55.7, 40.3, 34.0, 32.1, 31.6, 29.7, 29.4, 27.8, 22.6, 14.1.

**LRMS** (ESI, APCI) *m/z*: calc'd for $C_{28}H_{36}BrO$ $[M+H]^+$ 465.2, found 465.7

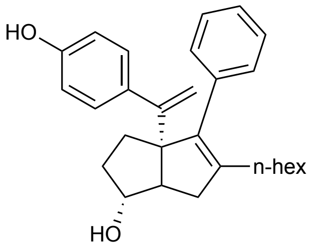


**(*exo*)-5-hexyl-3a-(1-(4-hydroxyphenyl)vinyl)-4-phenyl-1,2,3,3a,6,6a-hexahydropentalen-1-ol (13):** A solution of potassium hydroxide (13.5 mg, 0.24 mmol, 3.0 equiv.), tris(dibenzylideneacetone)dipalladium(0) (1.8 mg, 0.002 mmol, ca 0.03 equiv), and $^t$BuXPhos (2.8 mg, 0.007 mmol, ca 0.1 equiv) in 1,4-dioxane (ca 2 mL) in a reaction tube under nitrogen was treated with water and (*exo)*-3a-(1-(4-bromophenyl)vinyl)-5-hexyl-4-phenyl-1,2,3,3a,6,6a-hexahydropentalen-1-ol (**S9)** (30.7 mg, 0.07 mmol, 1.0 equiv) as a solution in dioxane (ca 1 mL). Water (ca 0.5 mL) was added, and the reaction mixture was heated to 80 °C for 16 hours. After reaction completion, the reaction was allowed to cool to ambient temperature before poured onto water and extracted with ethyl acetate three times. The combined organic layers were washed with water then brine, dried with $MgSO_4$, concentrated, and purified on silica in 20-50% EtOAc/hexanes eluent to afford the title compound. (6.2 mg, 26% yield). Note: for larger scale reactions palladium loading can be brought down to 0.01 equiv, and ligand to 0.04 equiv. Purity was established as the *exo* diastereomer by Method B: $t_R$ = 1.33 min, 95.6%.

**$^1$H NMR** (500 MHz, $CDCl_3$) δ 7.38 – 7.27 (m, 4H), 7.24 – 7.14 (m, 3H), 6.78 – 6.70 (m, 2H), 5.03 (d, *J* = 1.4 Hz, 1H), 4.96 (d, *J* = 1.5 Hz, 1H), 4.74 (s, 1H), 3.95 (s, 1H), 2.38 (dd, *J* = 16.9, 9.4 Hz, 1H), 2.28 (d, *J* = 9.3 Hz, 1H), 2.12 – 1.93 (m, 4H), 1.78 – 1.62 (m, 3H), 1.33 (dd, *J* = 14.0, 6.7 Hz, 3H), 1.31 – 1.17 (m, 5H), 0.86 (t, *J* = 7.1 Hz, 3H).

**$^{13}$C NMR** (126 MHz, $CDCl_3$) δ 154.6, 154.0, 141.1, 139.1, 137.4, 136.6, 129.7, 129.0, 127.6, 126.6, 114.5, 82.2, 69.5, 55.7, 40.3, 34.0, 32.0, 31.7, 29.7, 29.4, 27.8, 22.6, 14.1.

**LRMS** (ESI, APCI) *m/z*: calc'd for $C_{28}H_{37}O_2$ $[M+H]^+$ 403.3, found 403.9

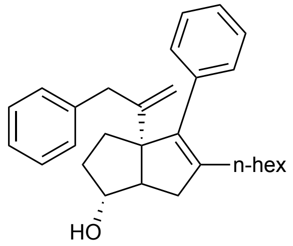


**(1*R*,3a*R*)-5-hexyl-4-phenyl-3a-(3-phenylprop-1-en-2-yl)-1,2,3,3a,6,6a-hexahydropentalen-1-ol (14):** According to the general procedure, (5-(methoxymethoxy)hept-6-en-1-yn-1-yl)benzene (57.0 mg, 0.25 mmol) was reacted with 3-phenyl-1-propyne (110 µL, 0.9 mmol) and purified in 5-20% EtOAc/hexanes eluent to afford the title compound (38.7 mg, 39% yield over two steps). Purity was established as the *exo* diastereomer by Method B: $t_R$ = 3.89 min, >99%.
**$^1$H NMR** (500 MHz, $CDCl_3$) δ 7.34 – 7.27 (m, 4H), 7.25 – 7.17 (m, 4H), 7.11 – 7.07 (m, 2H), 4.93 (d, *J* = 1.1 Hz, 1H), 4.50 (d, *J* = 1.4 Hz, 1H), 3.99 (s, 1H), 3.50 (d, *J* = 16.4 Hz, 1H), 3.39 (d, *J* = 16.4 Hz, 1H), 2.89 (dd, *J* = 17.2, 9.3 Hz, 1H), 2.39 (d, 1H), 2.25 (dd, *J* = 17.3, 2.0 Hz, 1H), 2.17 – 2.03 (m, 3H), 1.80 – 1.70 (m, 1H), 1.56 – 1.51 (m, 1H), 1.42 – 1.35 (m, 2H), 1.28 – 1.12 (m, 7H), 0.84 (t, *J* = 7.1 Hz, 3H).
**$^{13}$C NMR** (126 MHz, $CDCl_3$) δ 153.2, 140.9, 140.1, 139.1, 137.3, 129.7, 129.3, 128.3, 127.7, 126.5, 126.0, 95.5, 82.5, 70.2, 55.6, 40.7, 39.4, 34.2, 31.6, 30.2, 29.5, 29.2, 28.1, 22.6, 14.1.
**LRMS** (ESI, APCI) *m/z*: calc'd for $C_{29}H_{39}O$ $[M+H]^+$ 401.3, found 401.0

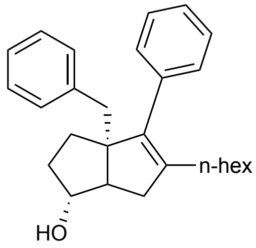


**(*exo*)-3a-benzyl-5-hexyl-4-phenyl-1,2,3,3a,6,6a-hexahydropentalen-1-ol (15):**
A solution of (**12**) (9.0 mg, 0.02 mmol, 1.0 equiv) in ethylene glycol was treated with hydrazine hydrate (0.5 mL, 0.16 mmol, 8 equiv) and heated to 100 °C for 1 hour. Potassium hydroxide (19 mg, 0.3 mmol, 15 equiv) was added and the reaction mixture was stirred at 150 °C for 48 h, at which point the reaction was allowed to come to ambient temperature, poured onto water and extracted with EtOAc three times. The combined organic layers were dried with $MgSO_4$, filtered, and concentrated. The crude reaction mixture was purified on silica in 10-20% EtOAc/Hexanes eluent (2.6 mg, 30% yield). Purity was established as the *exo* diastereomer by Method D: $t_{R.}$ = 1.35 min, 95.3%.
**$^1$H NMR** (600 MHz, $CDCl_3$) δ 7.37 – 7.33 (m, 3H), 7.30 – 7.27 (m, 1H), 7.24 – 7.22 (m, 1H), 7.21 – 7.17 (m, 3H), 7.16 – 7.11 (m, 2H), 3.83 (s, 1H), 2.93 (d, *J* = 13.6 Hz, 1H), 2.73 (d, *J* = 13.6 Hz, 1H), 2.38 (dd, *J* = 15.6, 8.4 Hz, 1H), 2.34 (d, *J* = 10.6 Hz, 1H), 2.05 – 1.99 (m, 1H), 1.89 – 1.83 (m, 2H), 1.74 – 1.67 (m, 3H), 1.49 – 1.44 (m, 1H), 1.29 – 1.18 (m, 5H), 1.12 (dt, *J* = 6.0, 4.2 Hz, 3H), 0.82 (td, *J* = 7.2, 0.6 Hz, 3H).
**$^{13}$C NMR** (101 MHz, $CDCl_3$) δ 140.9, 139.8, 139.4, 138.0, 130.5, 129.9, 127.9, 126.5, 126.0, 81.8, 64.3, 52.6, 44.0, 38.9, 33.8, 32.1, 31.6, 29.2, 29.0, 27.8, 22.6, 14.1.
**LRMS** (ESI, APCI) *m/z*: calc'd for $C_{26}H_{35}$ $[M-H_2O]^+$ 358.3, found 358.3

#### c. R3: Internal Styrene Modifications 16 – 23

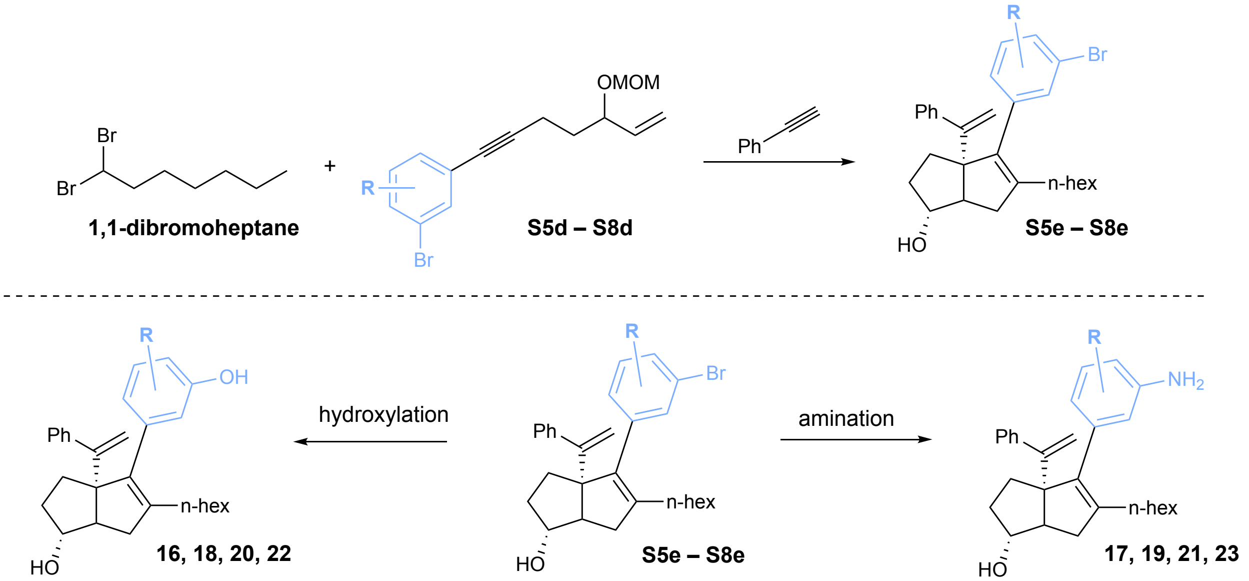


**General Sonogashira coupling procedure (S5a – S8a):**

A roundbottom flask equipped with magnetic stir bar was charged with bis(triphenylphosphine) palladium dichloride (0.01 equiv) and copper iodide (0.03 equiv). The flask was placed under nitrogen and triethylamine (1M with respect to aryl halide) was added via syringe. The solution was treated with iodobenzene (1.0 equiv), then sparged with nitrogen for 30 minutes. 4-pentyn-1-ol (1.2 equiv) was then added via syringe. The sparging needle was removed from the solution and replaced with a vent needle under positive nitrogen pressure. The solution was vigorously stirred at 60°C for 2 hours, at which point the reaction was complete by TLC. The reaction was cooled and precipitated with ether. The entire reaction was filtered over a plug of celite (eluted with ether). The filtrate was concentrated *in vacuo* to afford a rust-colored oil, which was purified on silica (10–30% EtOAc/hexanes eluent) to afford the title compounds.

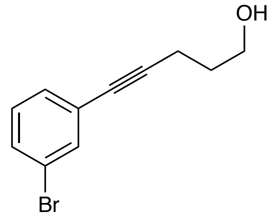


**5-(3-bromophenyl)pent-4-yn-1-ol (S5a):** According to the general procedure, 1-bromo-3-iodobenzene (3.6 g, 12.5 mmol) was reacted with 4-pentyn-1-ol (1.3 g, 15 mmol) to give the title compound as a yellow oil. (3.1 g, 92% yield).

**$^{1}$H NMR** (500 MHz, $CDCl_3$) δ 7.54 (t, *J* = 1.8 Hz, 1H), 7.40 (ddd, *J* = 8.1, 2.0, 1.0 Hz, 1H), 7.31 (dt, *J* = 7.7, 1.3 Hz, 1H), 7.15 (t, *J* = 7.9 Hz, 1H), 3.81 (t, *J* = 6.2 Hz, 2H), 2.54 (t, *J* = 7.0 Hz, 2H), 1.86 (tt, J = 6.9, 6.1 Hz, 2H), 1.59 (s, 1H).

**$^{13}$C NMR** (126 MHz, $CDCl_3$) δ 134.3, 130.8, 130.1, 129.6, 125.7, 122.0, 90.9, 79.7, 61.7, 31.2, 15.9.

**LRMS** (EI) *m/z:* calc'd for $C_{11}H_{11}BrO$ $[M]^+$ 238.0, found 238.0.

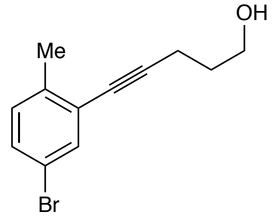


**5-(5-bromo-2-methylphenyl)pent-4-yn-1-ol (S6a):** According to the general procedure, 4-methyl-2-iodo-1-methylbenzene (2.5 mL, 17 mmol) was reacted with 4-pentyn-1-ol (2.2 mL, 22 mmol) to give the title compound as a yellow oil (3.6 g, 81% yield).

**$^{1}$H NMR** (600 MHz, $CDCl_3$) δ 7.48 (d, *J* = 2.2 Hz, 1H), 7.28 (dd, *J* = 8.2, 2.2 Hz, 1H), 7.03 (d, *J* = 8.2 Hz, 1H), 3.83 (t, *J* = 6.1 Hz, 2H), 2.58 (t, *J* = 6.9 Hz, 2H), 2.34 (s, 3H), 1.87 (p, *J* = 6.8 Hz, 2H), 1.55 (s, 1H).

**$^{13}$C NMR** (126 MHz, $CDCl_3$) δ 138.8, 134.3, 130.8, 130.6, 125.6, 118.6, 94.7, 78.7, 61.7, 31.4, 20.3, 16.1.

**LRMS** (EI) *m/z*: calc'd for $C_{12}H_{13}BrO$ $[M]^+$ 252.0, found 252.0

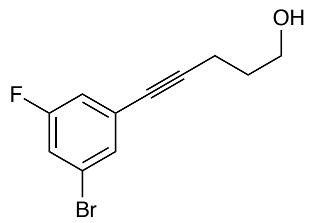


**5-(3-bromo-5-fluorophenyl)pent-4-yn-1-ol (S7a):** According to the general procedure, 1-bromo-3-fluoro-5-iodobenzene (4.6 g, 15 mmol) was reacted with 4-pentyn-1-ol (1.5 g, 18 mmol) to give the title compound as a yellow oil (3.5 g 90% yield).
**$^{1}$H NMR** (500 MHz, $CDCl_3$) δ 7.32 (t, *J* = 1.6 Hz, 1H), 7.16 (dt, *J* = 8.2, 2.1 Hz, 1H), 7.02 (ddd, *J* = 9.0, 2.4, 1.3 Hz, 1H), 3.79 (t, *J* = 6.2 Hz, 2H), 2.53 (t, *J* = 7.0 Hz, 2H), 1.85 (p, *J* = 6.6 Hz, 2H), 1.69 (s, 1H).
**$^{19}$F NMR** (282 MHz, $CDCl_3$) δ -111.19 (t, *J* = 8.7 Hz).
**$^{13}$C NMR** (126 MHz, $CDCl_3$) δ 162.1 (d, *J* = 250.6 Hz), 130.4 (d, *J* = 3.3 Hz), 127.0 (d, *J* = 10.3 Hz), 122.2 (d, *J* = 10.7 Hz), 118.7 (d, *J* = 24.6 Hz), 117.3 (d, *J* = 22.6 Hz), 92.2, 78.7 (d, *J* = 3.5 Hz), 61.5, 31.1, 15.9.
**LRMS** (EI) *m/z*: calc'd for $C_{11}H_9BrFO$ $[M]^+$ 256.0, found 256.1

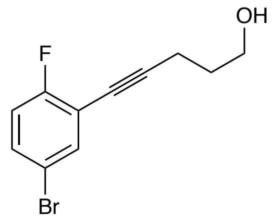


**5-(5-bromo-2-fluorophenyl)pent-4-yn-1-ol (S8a):** According to the general procedure, 4-bromo-1-fluoro-2-iodobenzene (2.6 mL, 20 mmol) was reacted with 4-pentyn-1-ol (2.2 mL, 22 mmol) to give the title compound as a yellow oil (3.9 g, 76% yield).
**$^{1}$H NMR** (500 MHz, $CDCl_3$) δ 7.51 (dd, *J* = 6.3, 2.5 Hz, 1H), 7.35 (ddd, *J* = 8.8, 4.6, 2.6 Hz, 1H), 6.93 (t, *J* = 8.8 Hz, 1H), 3.83 (t, *J* = 6.1 Hz, 2H), 2.59 (t, *J* = 6.9 Hz, 2H), 1.88 (tt, *J* = 6.7, 6.1 Hz, 2H), 1.39 (s, 1H).
**$^{13}$C NMR** (126 MHz, $CDCl_3$) δ 161.9 (d, *J* = 251.0 Hz), 135.9 (d, *J* = 1.9 Hz), 132.1 (d, *J* = 7.8 Hz), 117.0 (d, *J* = 22.8 Hz), 116.0 (d, *J* = 3.8 Hz), 114.3 (d, *J* = 17.3 Hz), 96.5 (d, *J* = 3.4 Hz), 73.2, 61.6, 31.1, 16.1.
**$^{19}$F NMR** (282 MHz, $CDCl_3$) δ -113.28 (ddd, *J* = 9.0, 6.3, 4.6 Hz).
**LRMS** (EI) *m/z*: calc'd for $C_{11}H_{10}BrFO$ $[M]^+$ 256.0, found 256.0

**General Swern Oxidation Procedure (S5b – S8b):** Under nitrogen, a solution of oxalyl chloride (1.1 equiv) in DCM (0.1 M with respect to alcohol) was cooled to -78 °C. A solution of dimethylsulfoxide (1.3 equiv) in DCM was added dropwise. After effervescence ceased (ca. 30 minutes), the required alcohol (1.0 equiv) was added dropwise in DCM. The reaction mixture was stirred at -78 °C for 1.5 h before the addition of triethylamine (2.5 equiv). The solution was allowed to warm to room temperature before the addition of saturated ammonium chloride (excess). The reaction mixture was then poured onto water and extracted with EtOAc, dried with $MgSO_4$, concentrated, and purified by silica gel chromatography.

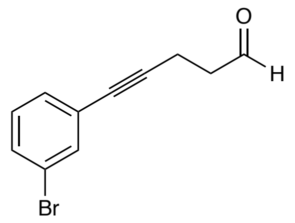


**5-(3-bromophenyl)pent-4-ynal (S5b):**
According to the general procedure, 5-(3-bromophenyl)pent-4-yn-1-ol (**S5a**) (4.2 g, 18 mmol) was reacted to give the title compound. The crude oil was purified on silica gel with 10-50% EtOAc/Hexanes eluent (3.4g, 81%).
**$^1$H NMR** (600 MHz, $CDCl_3$) δ 9.85 (t, $J$ = 1.1 Hz, 1H), 7.53 (t, $J$ = 1.8 Hz, 1H), 7.41 (ddd, $J$ = 8.0, 2.1, 1.0 Hz, 1H), 7.30 (dt, $J$ = 7.7, 1.4 Hz, 1H), 7.15 (t, $J$ = 7.9 Hz, 1H), 2.80 – 2.71 (m, 4H).
**$^{13}$C NMR** (126 MHz, $CDCl_3$) δ 200.1, 134.4, 131.1, 130.1, 129.7, 125.3, 122.0, 89.3, 80.0, 42.5, 12.6.
**LRMS** (ESI) *m/z*: calc'd for $C_{11}H_9BrO$ $[M]^+$ 236.0, found 236.0

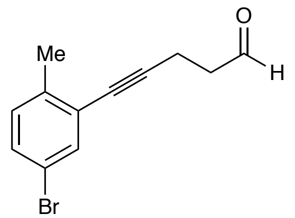


**5-(5-bromo-2-methylphenyl)pent-4-ynal (S6b):** According to the general procedure, 5-(5-bromo-2-methylphenyl)pent-4-yn-1-ol **(S6a**) (3.6 g, 14 mmol) was reacted to give the title compound. The crude oil was purified on silica gel with 10% EtOAc/Hex (0.6 g, 17%).
**$^1$H NMR** (500 MHz, $CDCl_3$) δ 9.86 (s, 1H), 7.48 (d, $J$ = 2.2 Hz, 1H), 7.29 (dd, $J$ = 8.2, 2.2 Hz, 1H), 7.04 (d, $J$ = 8.2 Hz, 1H), 2.78 (s, 4H), 2.33 (s, 3H).
**$^{13}$C NMR** (151 MHz, $CDCl_3$) δ 200.2, 172.5, 139.0, 134.3, 130.8, 125.1, 118.6, 93.0, 79.1, 42.7, 20.2, 12.8.
**LRMS** (EI) *m/z*: calc'd for $C_{12}H_{11}BrO$ $[M]^+$ 250.0, found 250.0

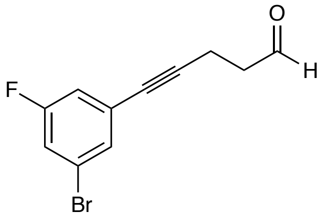


**5-(3-bromo-5-fluorophenyl)pent-4-ynal (S7b):** According to the general procedure, 5-(3-bromo-5-fluorophenyl)pent-4-yn-1-ol (**S7a**) (1.4 g, 5.5 mmol) was reacted to give the title compound. The crude oil was purified on silica gel with 10-20% EtOAc/Hex, (700 mg, 50%).
**$^1$H NMR** (500 MHz, $CDCl_3$) δ 9.81 (t, *J* = 1.0 Hz, 1H), 7.30 – 7.27 (m, 1H), 7.15 (ddd, *J* = 8.1, 2.4, 1.8 Hz, 1H), 6.99 (ddd, *J* = 9.1, 2.4, 1.3 Hz, 1H), 2.79 – 2.71 (m, 2H), 2.73 – 2.66 (m, 2H).
**$^{13}$C NMR** (126 MHz, $CDCl_3$) δ 199.8, 162.1 (d, *J* = 250.7 Hz), 130.5 (d, *J* = 3.3 Hz), 126.6 (d, *J* = 10.4 Hz), 122.2 (d, *J* = 10.5 Hz), 118.9 (d, *J* = 24.5 Hz), 117.3 (d, *J* = 22.7 Hz), 90.6, 78.9 (d, *J* = 3.5 Hz), 42.3, 12.5.
**$^{19}$F NMR** (282 MHz, $CDCl_3$) δ -111.00 (t, *J* = 8.6 Hz).
**LRMS (**EI) *m/z*: calc'd for $C_{11}H_9BrFO$ $[M]^+$ 254.0, found 254.0

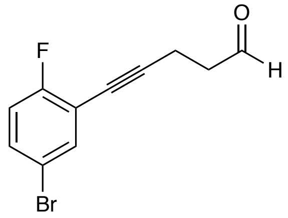


**5-(5-bromo-2-fluorophenyl)pent-4-ynal (S8b):** According to the general procedure, 5-(5-bromo-2-fluorophenyl)pent-4-yn-1-ol (**S8a**) (3.9 g, 15 mmol) was reacted to give the title compound. The crude oil was purified on silica gel with 10-20% EtOAc/Hex (2.4 g, 62%).
**$^1$H NMR** (500 MHz, $CDCl_3$) δ 9.84 (s, 1H), 7.49 (dd, *J* = 6.3, 2.5 Hz, 1H), 7.35 (ddd, *J* = 8.8, 4.6, 2.6 Hz, 1H), 6.93 (t, *J* = 8.8 Hz, 1H), 2.82 – 2.73 (m, 4H).
**$^{13}$C NMR** (126 MHz, $CDCl_3$) δ 199.9, 161.9 (d, *J* = 251.5 Hz), 135.9 (d, *J* = 1.8 Hz), 132.4 (d, *J* = 7.7 Hz), 117.0 (d, *J* = 22.5 Hz), 116.0 (d, *J* = 3.4 Hz), 113.9 (d, *J* = 20.5 Hz), 94.8 (d, *J* = 3.4 Hz), 73.5, 42.3, 12.8.
**$^{19}$F NMR** (282 MHz, $CDCl_3$) δ -113.04 (ddd, *J* = 8.9, 6.3, 4.5 Hz).
**LRMS** (EI) *m/z*: calc'd for $C_{11}H_8BrFO$ $[M]^+$ 254.0, found 254.0

**General Grignard Addition Procedure (S5c – S8c):** Under nitrogen, a solution of aldehyde **S2b – S5b** (1.0 equiv) in anhydrous THF (0.5 M) was cooled to -78 °C. The solution was treated with vinylmagnesium bromide (1.0M solution in THF, 1.5 equiv). The reaction was stirred and allowed to warm to room temperature over 16 h, then saturated ammonium chloride was added. The reaction mixture was poured onto water and extracted with EtOAc, dried with $MgSO_4$, and concentrated before purification on silica to give the title compounds.

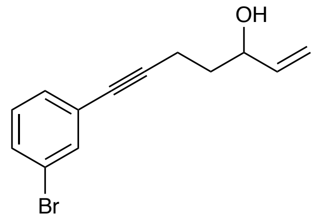


**7-(3-bromophenyl)hept-1-en-6-yn-3-ol (S5c):** According to the general procedure, 5-(3-bromophenyl)pent-4-ynal (**S5b**) (3.4 g, 14 mmol) was reacted with vinylmagnesium bromide (21 mL of a 1.0M solution in THF, 21 mmol). The crude oil was purified on silica gel with 5-10% EtOAc/Hexanes eluent, (2.0 g, 53% yield).

**$^1$H NMR** (600 MHz, $CDCl_3$) δ 7.53 (t, *J* = 1.8 Hz, 1H), 7.40 (ddd, *J* = 8.1, 2.1, 1.1 Hz, 1H), 7.30 (dt, *J* = 7.8, 1.3 Hz, 1H), 7.14 (t, *J* = 7.9 Hz, 1H), 5.90 (ddd, *J* = 17.2, 10.4, 6.1 Hz, 1H), 5.29 (dt, *J* = 17.3, 1.4 Hz, 1H), 5.16 (dt, *J* = 10.5, 1.3 Hz, 1H), 4.31 (qt, J = 6.0, 1.3 Hz, 1H), 2.72 – 2.33 (m, 2H), 1.96 – 1.69 (m, 2H).

**$^{13}$C NMR** (126 MHz, $CDCl_3$) δ 140.4, 134.3, 130.8, 130.1, 129.6, 125.8, 122.0, 115.3, 91.0, 79.7, 71.9, 35.4, 15.6.

**LRMS** (ESI) *m/z*: calc'd for $C_{13}H_{14}BrO$ $[M+H]^+$ 265.0, found 265.0

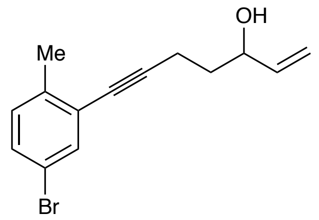


**7-(5-bromo-2-methylphenyl)hept-1-en-6-yn-3-ol (S6c):** According to the general procedure, 5-(5-bromo-2-methylphenyl)pent-4-ynal (**S6b**) (0.6 g, 2.4 mmol) was reacted with vinyl magnesium bromide (3.6 mL of a 1.0 M solution in THF, 3.6 mmol). The crude oil was purified on silica gel in 20% EtOAc/Hexanes eluent, (0.2 g, 32% yield).

**$^1$H NMR** (600 MHz, $CDCl_3$) δ 7.49 (d, *J* = 2.2 Hz, 1H), 7.28 (dd, *J* = 8.2, 2.2 Hz, 1H), 7.04 (d, *J* = 8.2 Hz, 1H), 5.95 – 5.87 (m, 1H), 5.30 (dt, *J* = 17.3, 1.4 Hz, 1H), 5.17 (dt, *J* = 10.6, 1.4 Hz, 1H), 4.34 (q, *J* = 6.4 Hz, 1H), 2.57 (qt, *J* = 17.1, 7.1 Hz, 2H), 2.35 (s, 3H), 1.84 (q, *J* = 6.9 Hz, 2H), 1.67 (s, 1H).

**$^{13}$C NMR** (126 MHz, $CDCl_3$) δ 140.4, 138.8, 134.3, 130.8, 130.6, 125.6, 118.6, 115.3, 94.8, 78.8, 72.0, 35.6, 20.3, 15.8.

**LRMS** (ESI) m/*z*: calc'd for $C_{14}H_{16}BrO$ $[M+H]^+$ 279.0 , found 279.0.

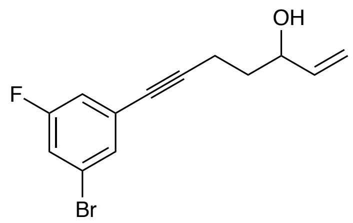

**7-(3-bromo-5-fluorophenyl)hept-1-en-6-yn-3-ol (S7c):** According to the general procedure, 5-(3-bromo-5-fluorophenyl)pent-4-yn-1-ol **(S7b**) (700 mg, 2.7 mmol) was reacted with vinylmagnesium bromide (5 mL of a 1.0M solution in THF, 5 mmol, 1.9 equiv). The crude oil was purified on silica gel in 20% EtOAc/Hexanes, (497 mg, 64% yield).

**$^{1}$H NMR** (500 MHz, $CDCl_3$) δ 7.33 (s, 1H), 7.17 (ddd, *J* = 8.2, 2.4, 1.8 Hz, 1H), 7.02 (ddd, *J* = 9.1, 2.5, 1.3 Hz, 1H), 5.90 (ddd, *J* = 17.1, 10.4, 6.1 Hz, 1H), 5.30 (dt, *J* = 17.2, 1.4 Hz, 1H), 5.18 (dt, *J* = 10.4, 1.3 Hz, 1H), 4.30 (d, *J* = 10.3 Hz, 1H), 2.60 – 2.45 (m, 3H), 1.85 – 1.78 (m, 3H), 1.66 (d, *J* = 4.3 Hz, 1H).

**$^{13}$C NMR** (126 MHz, $CDCl_3$) δ 140.3, 130.4 (d, *J* = 3.2 Hz), 122.2 (d, *J* = 10.7 Hz), 118.8 (d, *J* = 24.6 Hz), 117.3 (d, *J* = 22.6 Hz), 115.3, 92.3, 78.7, 71.9, 35.3, 15.6.

**$^{19}$F NMR** (282 MHz, $CDCl_3$) δ -111.22 (t, *J* = 8.6 Hz).

**LRMS** (ESI) *m/z*: calc'd for $C_{13}H_{12}BrFO$ $[M]^+$ 282.0, found 282.1

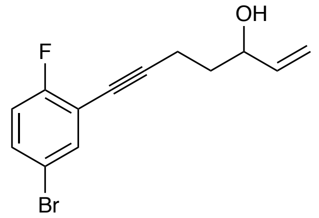

**7-(5-bromo-2-fluorophenyl)hept-1-en-6-yn-3-ol (S8c):** According to the general procedure, 5-(5-bromo-2-fluorophenyl)pent-4-ynal (**S8b**) (2.4 g, 9.3 mmol) was reacted with vinylmagnesium bromide (14 mL of a 1.0M solution in THF, 14 mmol). The crude oil was purified on silica gel in 20% EtOAc/Hexanes eluent, (1.7 g, 64% yield).

**$^{1}$H NMR** (600 MHz, $CDCl_3$) δ 7.51 (dd, *J* = 6.4, 2.4 Hz, 1H), 7.36 – 7.33 (m, 1H), 6.93 (t, *J* = 8.9, Hz, 1H), 5.90 (m, 1H), 5.31 (dq, *J* = 17.2, 1.2 Hz, 1H), 5.17 (dq, *J* = 10.4, 1.2 Hz, 1H), 4.34 (q, *J* = 6.5 Hz, 1H), 2.58 (qt, *J* = 17.2, 7.1 Hz, 3H), 1.87 – 1.80 (m, 2H), 1.67 (s, 1H).

**$^{13}$C NMR** (126 MHz, $CDCl_3$) δ 161.9 (d, *J* = 250.6 Hz), 140.3, 135.9 (d, *J* = 1.9 Hz), 132.1 (d, *J* = 7.8 Hz), 117.0 (d, *J* = 23.0 Hz), 116.0 (d, *J* = 3.6 Hz), 115.3, 114.4 (d, *J* = 17.3 Hz), 96.6 (d, *J* = 3.4 Hz), 73.2, 71.9, 35.3, 15.8.

**$^{19}$F NMR** (282 MHz, $CDCl_3$) δ -113.24 (ddd, *J* = 9.0, 6.3, 4.5 Hz).

**LRMS** (ESI) *m/z*: calc'd for $C_{13}H_{13}BrFO$ $[M+H]^+$ 283.0, found 283.0

**General procedure for methoxymethyl (MOM) ether alcohol protection (S5d–S8d):** The required enyne (**S5c – S8c**) (1.0 equiv.) was dissolved in DCM (0.5 M), followed by diisopropylethyl amine (1.25 equiv.) Chloromethyl methyl ether (1.5 equiv.) was added and the reaction mixture was stirred at 30 °C until completion was detected by TLC (typically 1-4 hours). The reaction mixture was cooled to room temperature, poured onto water, and extracted with DCM. The combined organic layers were washed with dilute HCl (1 M) and brine, dried with $MgSO_4$, filtered, and concentrated before purification on silica to give the title compounds.

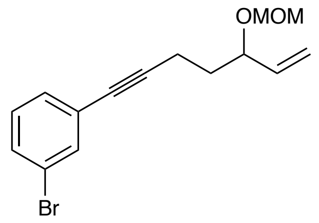


**1-bromo-3-(5-(methoxymethoxy)hept-6-en-1-yn-1-yl)benzene (S5d):** According to the general procedure, 7-(3-bromophenyl)hept-1-en-6-yn-3-ol (**S5c**) (2.0 g, 7.5 mmol) was reacted with chloromethyl methyl ether. The crude oil was purified in 5% EtOAc/hexanes eluent (2.1 g, 92% yield).
**$^1$H NMR** (600 MHz, $CDCl_3$) δ 7.53 (t, *J* = 1.8 Hz, 1H), 7.40 (m, 1H), 7.32 – 7.29 (m, 1H), 7.14 (t, *J* = 7.9 Hz, 1H), 5.70 (ddd, J = 17.8, 10.3, 7.6 Hz, 1H), 5.31 – 5.21 (m, 2H), 4.73 (d, J = 6.8 Hz, 1H), 4.57 (d, *J* = 6.7 Hz, 1H), 4.19 (td, *J* = 7.8, 5.2 Hz, 1H), 3.40 (s, 3H), 2.63 – 2.42 (m, 2H), 1.93 – 1.86 (m, 1H), 1.84 – 1.76 (m, 1H).
**$^{13}$C NMR** (126 MHz, $CDCl_3$) δ 137.6, 134.3, 130.8, 130.1, 129.6, 125.9, 122.0, 117.9, 93.8, 91.0, 79.6, 75.8, 55.5, 34.2, 15.6.
**LRMS** (ESI) *m/z*: calc'd for $C_{15}H_{16}BrO_2$ $[M-H]^+$ 307.0, found 307.0, calc'd for $C_{14}H_{14}BrO$ $[M-OCH_3]^+$ 277.0, found 277.1, calc'd for $C_{13}H_{12}BrO$ $[M-C_2H_5O]^+$ 263.0, found 263.0.

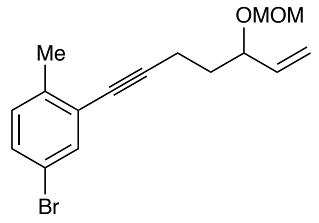


**4-bromo-2-(5-(methoxymethoxy)hept-6-en-1-yn-1-yl)-1-methylbenzene (S6d):** According to the general procedure, 7-(5-bromo-2-methylphenyl)hept-1-en-6-yn-3-ol (**S6c**) (0.2 g, 0.7 mmol) was reacted with chloromethyl methyl ether. The crude oil was purified in 2-10% EtOAc/hexanes eluent (0.18 g, 78% yield).
**$^1$H NMR** (600 MHz, $CDCl_3$) δ 7.48 (d, *J* = 2.2 Hz, 1H), 7.28 (dd, *J* = 8.2, 2.2 Hz, 1H), 7.04 (d, *J* = 8.2 Hz, 1H), 5.71 (m, 1H), 5.31 – 5.20 (m, 2H), 4.74 (d, *J* = 6.7 Hz, 1H), 4.57 (d, *J* = 6.7 Hz, 1H), 4.22 (td, *J* = 7.7, 5.3 Hz, 1H), 3.40 (s, 3H), 2.63 – 2.50 (m, 2H), 2.35 (s, 3H), 1.91 (dtd, *J* = 13.8, 7.6, 6.2 Hz, 1H), 1.81 (dtd, *J* = 13.2, 7.6, 5.4 Hz, 1H).
**$^{13}$C NMR** (126 MHz, $CDCl_3$) δ 138.8, 137.6, 134.3, 130.8, 130.6, 125.7, 118.6, 117.9, 94.9, 93.8, 78.6, 75.8, 55.5, 34.5, 20.2, 15.8.
**LRMS** (ESI) *m/z*: calc'd for $C_{16}H_{19}BrO_2$ $[M-H]^+$ 321.0, found 321.0, calc'd for $C_{15}H_{16}BrO$ $[M-OCH_3]^+$ 291.0, found 291.0, calc'd for $C_{14}H_{14}BrO$ $[M-C_2H_5O]^+$ 277.0, found 277.0.

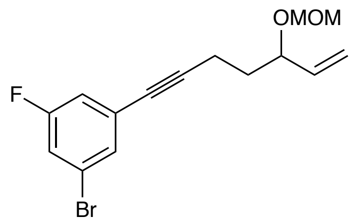

**1-bromo-3-fluoro-5-(5-(methoxymethoxy)hept-6-en-1-yn-1-yl)benzene (S7d):** According to the general procedure, 7-(3-bromo-5-fluorophenyl)hept-1-en-6-yn-3-ol (**S7c**) (497.1 mg, 1.8 mmol) was reacted with chloromethyl methyl ether. The crude oil was purified in 2-10% EtOAc/hexanes (270 mg, 47% yield).

**$^1$H NMR** (500 MHz, $CDCl_3$) δ 7.32 (s, 1H), 7.16 (ddd, *J* = 8.2, 2.4, 1.7 Hz, 1H), 7.02 (ddd, *J* = 9.1, 2.4, 1.3 Hz, 1H), 5.70 (ddd, *J* = 17.2, 10.3, 7.5 Hz, 1H), 5.28 (ddd, *J* = 17.3, 1.6, 1.0 Hz, 1H), 5.24 (ddd, *J* = 10.3, 1.6, 0.8 Hz, 1H), 4.73 (d, *J* = 6.7 Hz, 1H), 4.56 (d, *J* = 6.7 Hz, 1H), 3.40 (s, 3H), 2.58 – 2.42 (m, 2H), 1.88 (dtd, *J* = 13.9, 7.8, 6.1 Hz, 1H), 1.80 (dtd, *J* = 13.6, 7.7, 5.3 Hz, 1H).

**$^{13}$C NMR** (126 MHz, $CDCl_3$) δ 162.1 (d, *J* = 250.6 Hz), 137.5, 130.4 (d, *J* = 3.3 Hz), 127.1 (d, *J* = 10.2 Hz), 122.2 (d, *J* = 10.6 Hz), 118.7 (d, *J* = 24.6 Hz), 117.8, 117.3 (d, *J* = 22.6 Hz), 93.8, 92.3, 78.6 (d, *J* = 3.4 Hz), 75.7, 55.5, 34.1, 15.5.

**$^{19}$F NMR** (282 MHz, $CDCl_3$) δ -111.22 (t, *J* = 8.6 Hz).

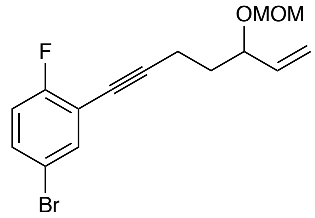

**4-bromo-1-fluoro-2-(5-(methoxymethoxy)hept-6-en-1-yn-1-yl)benzene (S8d):** According to the general procedure, 7-(5-bromo-2-fluorophenyl)hept-1-en-6-yn-3-ol (**S8c**) (1.7 g, 6 mmol) was reacted with chloromethyl methyl ether. The crude oil was purified in 2-10% EtOAc/hexanes eluent (1.6 g, 80% yield).

**$^1$H NMR** (600 MHz, $CDCl_3$) δ 7.50 (dd, *J* = 6.3, 2.5 Hz, 1H), 7.34 (m, 1H), 6.93 (t, *J* = 8.8 Hz, 1H), 5.70 (m, 1H), 5.31 – 5.21 (m, 2H), 4.73 (d, *J* = 6.7 Hz, 1H), 4.57 (d, *J* = 6.7 Hz, 1H), 4.20 (td, *J* = 7.8, 5.3 Hz, 1H), 3.40 (s, 3H), 2.63 – 2.50 (m, 2H), 1.94 – 1.87 (m, 1H), 1.85 – 1.78 (m, 1H).

**$^{13}$C NMR** (126 MHz, $CDCl_3$) δ 161.9 (d, *J* = 251.0 Hz), 137.6, 135.9 (d, *J* = 1.8 Hz), 132.1 (d, *J* = 7.8 Hz), 117.9, 117.0 (d, *J* = 22.6 Hz), 116.0 (d, *J* = 3.6 Hz), 114.5 (d, *J* = 17.4 Hz), 96.6 (d, *J* = 3.4 Hz), 93.9, 75.7, 73.1, 55.5, 34.1, 15.8.

**$^{19}$F NMR** (282 MHz, $CDCl_3$) δ -113.23 (ddd, *J* = 9.3, 6.4, 4.6 Hz).

**LRMS** (ESI) *m/z*: calc'd for $C_{15}H_{15}BrFO_2$ $[M\text{-}H]^+$ 325.0, found 325.0, calc'd for $C_{14}H_{13}BrFO$ $[M\text{-}OCH_3]^+$ 295.0, found 295.1, calc'd for $C_{13}H_{11}BrFO$ $[M\text{-}C_2H_5O]^+$ 281.0, found 281.0.

Hexahydropentalene formation was accomplished through slight modification of Whitby's procedure.[1] Prior to cyclization, all non-volatile reagents were dried by azeotropic removal of water using benzene. A dry round bottom flask containing bis(cyclopentadienyl)zirconium(IV) dichloride (1.2 equiv) under nitrogen, was dissolved in anhydrous, degassed tetrahydrofuran (THF, 50 mL/mmol enyne) and cooled to -78 °C. The resulting solution was treated with n-BuLi (2.4 equiv.) and the light yellow solution was stirred for 30 minutes. A solution of **S5d – S8d** (1.0 equiv) in anhydrous, degassed THF (5 mL/mmol) was added. The resulting salmon-colored mixture was stirred at -78 °C for 30 minutes, the cooling bath removed, and the reaction mixture was allowed to warm to ambient temperature with stirring (2.5 hours total). The reaction mixture was then cooled to -78 °C and the required 1,1-dibromoheptane (1.1 equiv) was added as a solution in anhydrous THF (5 mL/mmol) followed by freshly prepared lithium diisopropylamide (LDA, 1.0 M, 1.1 equiv.). After 15 minutes, a freshly prepared solution of lithium phenylacetylide (3.6 equiv.) in anhydrous THF was added dropwise and the resulting rust-colored solution was stirred at -78 °C for 1.5 hours. The reaction was quenched with methanol and saturated aqueous sodium bicarbonate and allowed to warm to room temperature, affording a light yellow slurry. The slurry was poured onto water and extracted with ethyl acetate four times. The combined organic layers were washed with brine, dried with $MgSO_4$, and concentrated *in vacuo.* The resulting yellow oil was passed through a short plug of silica (20% EtOAc/Hexanes eluent) and concentrated. The crude product was dissolved in acetonitrile and treated with concentrated aqueous hydrochloric acid (ca 5 equiv) and the resulting solution stirred at room temperature until completion of the reaction was detected (typically fewer than 10 minutes). The reaction mixture was concentrated and purified by silica gel chromatography (5-20% EtOAc/hexanes eluent) to afford the title compounds **S5e – S8e** as a 7:1 mixture of diastereomers, favoring the desired *exo*-isomer.

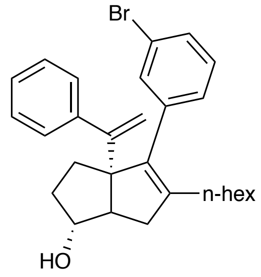


**(*exo*)-4-(3-bromophenyl)-5-hexyl-3a-(1-phenylvinyl)-1,2,3,3a,6,6a-hexahydropentalen-1-ol (S5e):** According to the general procedure, 1-bromo-3-(5-(methoxymethoxy)hept-6-en-1-yn-1-yl)benzene (**S5d**) (234 mg, 0.76 mmol) was reacted with 1,1-dibromoheptane (215 mg, 0.84 mmol) and phenylacetylide to afford the title compound (21.2 mg, 6% yield) over two steps.

**$^1$H NMR** (500 MHz, $CDCl_3$) δ 7.43 – 7.37 (m, 1H), 7.36 – 7.29 (m, 3H), 7.29 – 7.26 (m, 2H), 7.24 – 7.08 (m, 3H), 5.10 (d, *J* = 1.3 Hz, 1H), 5.00 (d, *J* = 1.2 Hz, 1H), 3.95 (s, 1H), 2.38 (dd, *J* = 17.0, 9.4 Hz, 1H), 2.30 (d, *J* = 9.5 Hz, 1H), 2.14 – 1.97 (m, 5H), 1.74 – 1.65 (m, 3H), 1.39 – 1.15 (m, 7H), 0.87 (t, *J* = 7.2 Hz, 3H).

**$^{13}$C NMR** (126 MHz, $CDCl_3$) δ 154.3, 144.0, 142.4, 139.6, 137.7, 132.5, 129.7, 129.2, 128.4, 127.8, 127.6, 126.8, 121.8, 115.3, 81.9, 69.3, 55.8, 40.3, 34.0, 32.1, 31.6, 29.7, 29.3, 27.7, 22.6, 14.1.

**LRMS** (ESI, APCI) *m/z*: calc'd for $C_{28}H_{36}BrO$ $[M+H]^+$ 465.2, found 465.7

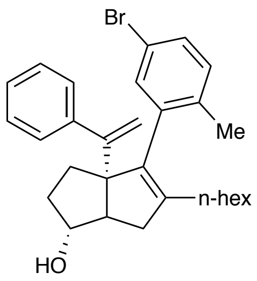


**(*exo*)-4-(5-bromo-2-methylphenyl)-5-hexyl-3a-(1-phenylvinyl)-1,2,3,3a,6,6a-hexahydropentalen-1-ol (S6e):** According to the general procedure, 4-bromo-2-(5-(methoxymethoxy)hept-6-en-1-yn-1-yl)-1-methylbenzene (**S6d**) (317.5 mg, 1 mmol) was reacted with 1,1-dibromoheptane (282.8 mg, 1.1 mmol) and phenylacetylide to afford the title compound (20.4 mg, 4% yield over two steps).

**$^{1}$H NMR** (500 MHz, $CDCl_3$) δ 7.39 – 7.34 (m, 2H), 7.32 – 7.27 (m, 3H), 7.26 – 7.22 (m, 2H), 7.11 (d, *J* = 8.2 Hz, 1H), 5.22 (d, *J* = 1.1 Hz, 1H), 4.85 (d, *J* = 1.1 Hz, 1H), 3.98 (dd, *J* = 3.8, 1.8 Hz, 1H), 2.63 (dd, *J* = 17.4, 10.0 Hz, 1H), 2.34 (ddt, *J* = 10.0, 3.0, 1.7 Hz, 1H), 2.15 (s, 3H), 2.09 – 1.99 (m, 3H), 1.87 (t, *J* = 7.9 Hz, 2H), 1.84 – 1.69 (m, 2H), 1.37 (td, *J* = 7.9, 5.1 Hz, 2H), 1.30 – 1.11 (m, 6H), 0.86 (t, *J* = 7.2 Hz, 3H).

**$^{13}$C NMR** (126 MHz, $CDCl_3$) δ 154.5, 141.4, 138.1, 136.4, 136.2, 133.2, 131.3, 129.7, 128.1, 127.8, 127.2, 126.8, 116.4, 81.3, 55.5, 40.3, 33.8, 33.0, 31.6, 30.1, 29.5, 27.1, 22.6, 19.5, 14.1.

**LRMS** (ESI, APCI) *m/z*: calc'd for $C_{29}H_{38}BrO$ $[M+H]^+$ 479.2, found 479.7

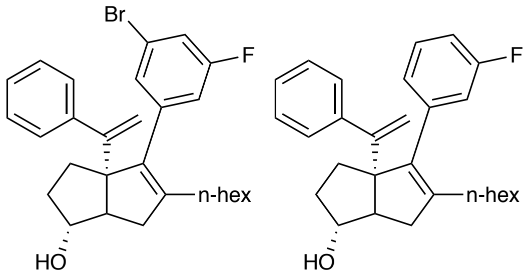


**(*exo*)-4-(3-bromo-5-fluorophenyl)-5-hexyl-3a-(1-phenylvinyl)-1,2,3,3a,6,6a-hexahydropentalen-1-ol (S7e):** According to the general procedure, 1-bromo-3-fluoro-5-(5-(methoxymethoxy)hept-6-en-1-yn-1-yl)benzene (**S7d**) (345.6 mg, 1 mmol) was reacted with 1,1-dibromoheptane (300.5 mg, 1.2 mmol) and phenylacetylide to afford both the desired compound and lithium-halogen exchange byproduct in an appreciable amount (72.4 mg combined, over two steps) This mixture was carried on without further purification. Spectral data are representative of both hydrodehalogenation (33%) and desired (67%) products.

**$^{1}$H NMR** (500 MHz, $CDCl_3$) δ 7.36 – 7.23 (m, 4H), 7.20 – 7.12 (m, 1H), 7.01 – 6.87 (m, 3H), 5.12 (d, *J* = 1.3 Hz, 0.5H), 5.10 (d, *J* = 1.4 Hz, 0.3H), 5.04 (d, *J* = 1.3 Hz, 0.6H), 5.02 (d, *J* = 1.4 Hz, 0.3H), 3.94 (s, 1H), 2.43 – 2.34 (m, 1H), 2.31 (dt, *J* = 9.4, 1.9 Hz, 1H), 2.15 – 2.07 (m, 1H), 2.06 – 1.99 (m, 2H), 1.72 – 1.60 (m, 3H), 1.36 (d, *J* = 7.8 Hz, 4H), 1.31 – 1.17 (m, 5H), 0.91 – 0.82 (m, 3H).

**$^{13}$C NMR** (126 MHz, $CDCl_3$) δ 154.4, 154.2, 143.7, 143.4, 129.1, 129.0, 128.5, 127.9, 127.8, 127.7, 127.6, 126.9, 126.8, 125.5, 117.4, 117.2, 116.5, 116.3, 115.52, 115.47, 115.4, 115.2, 113.6, 113.4, 81.9, 81.8, 69.3, 55.8, 40.3, 34.0, 32.1, 31.62, 31.59, 29.68, 29.65, 29.4, 29.3, 27.74, 27.67, 22.6, 14.1.
**$^{19}$F NMR** (282 MHz, $CDCl_3$) δ -111.56, -113.95.
**LRMS** (ESI, APCI) *m/z*: calc'd for $C_{28}H_{35}BrFO$ $[M+H]^+$ 483.2, found 483.1, calc'd for $C_{28}H_{36}FO$ $[M+H]^+$ 405.3, found 405.2

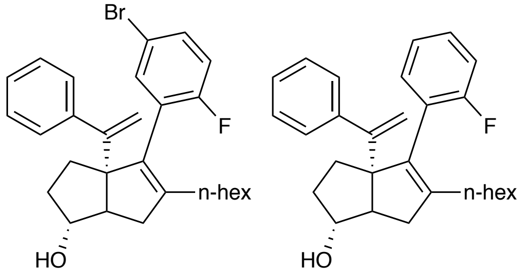


**(*exo*)-4-(5-amino-2-fluorophenyl)-5-hexyl-3a-(1-phenylvinyl)-1,2,3,3a,6,6a-hexahydropentalen-1-ol (S8e):** According to the general procedure, 4-bromo-1-fluoro-2-(5-(methoxymethoxy)hept-6-en-1-yn-1-yl)benzene (**S8d**) (417.6 mg, 1.8 mmol) was reacted with 1,1-dibromoheptane (371.6 mg, 1.4 mmol) and phenylacetylide, affording both the desired compound and lithium-halogen exchange byproduct in an appreciable amount (63.3 mg, combined, over two steps). This mixture was carried on without further purification. Spectral data are representative of both hydrodehalogenation (67%) and desired (33%) products.
**$^{1}$H NMR** (500 MHz, $CDCl_3$) δ 7.41 – 7.24 (m, 6H), 7.21 (td, *J* = 7.6, 1.7 Hz, 0.67H), 7.10 – 7.02 (m, 1H), 6.96 (t, *J* = 8.8 Hz, 0.33H), 5.12 (d, *J* = 1.3 Hz, 0.33H), 5.07 (d, *J* = 1.3 Hz, 0.67H), 4.90 (d, *J* = 1.3 Hz, 0.33H), 4.87 (d, *J* = 1.3 Hz, 0.67H), 4.11 – 3.90 (m, 1H), 2.54 (dd, *J* = 16.1, 8.7 Hz, 1H), 2.32 (d, *J* = 9.7 Hz, 1H), 2.12 – 2.02 (m, 2H), 1.94 (t, *J* = 7.7 Hz, 2H), 1.88 – 1.63 (m, 2H), 1.36 (q, *J* = 7.0, 6.4 Hz, 2H), 1.29 – 1.14 (m, 7H), 0.86 (q, *J* = 7.0 Hz, 3H).
**$^{13}$C NMR** (126 MHz, $CDCl_3$) δ 161.6, 160.8, 159.7, 158.8, 154.6, 154.4, 144.57, 144.55, 144.3, 143.3, 134.2, 134.1, 132.6, 131.8, 131.7, 131.5, 131.4, 128.5, 128.5, 127.9, 127.9, 127.5, 127.4, 126.8, 126.7, 124.5, 124.4, 123.08, 123.05, 117.1, 116.9, 115.74, 115.71, 115.5, 115.4, 115.3, 115.1, 82.0, 81.8, 69.69, 69.66, 55.4, 55.3, 40.6, 33.58, 33.56, 33.2, 31.61, 31.59, 30.00, 29.95, 29.3, 27.2, 27.2, 22.6, 14.1.
**$^{19}$F NMR** (282 MHz, $CDCl_3$) δ -113.58, -115.34.
**LRMS** (ESI, APCI) *m/z*: calc'd for $C_{28}H_{35}BrFO$ $[M+H]^+$ 483.2, found 483.1, calc'd for $C_{28}H_{36}FO$ $[M+H]^+$ 405.3, found 405.2.

**General Procedure for Hydroxyl Coupling (16, 18, 20, 22):** Potassium hydroxide (3.0 equiv.), tris(dibenzylideneacetone)dipalladium(0) (0.01 equiv.), and ᵗBuXPhos (0.04 equiv.) were placed in a reaction tube, which was evacuated and backfilled with nitrogen three times. The solids were then suspended in degassed 1,4-dioxane under nitrogen. The required brominated [3.3.0] bicycle (**S5e – S8e**) was added in 1,4-dioxane. Water (~10 equiv.) was added. The reaction mixture was heated to 80 °C and stirred for 16 hours. After stirring, the mixture was poured over water and extracted with EtOAc three times. The combined organic layers were washed with water and brine, dried with $MgSO_4$, concentrated, and purified on silica in 20% EtOAc/Hexanes eluent.

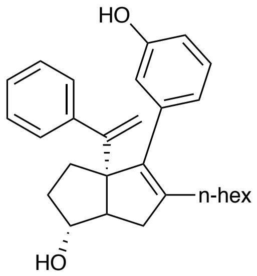


**(*exo*)-5-hexyl-4-(3-hydroxyphenyl)-3a-(1-phenylvinyl)-1,2,3,3a,6,6a-hexahydropentalen-1-ol (16):** (*exo*)-6-(3-bromophenyl)-5-hexyl-3-(methoxymethoxy)-6a-(1-phenylvinyl)-1,2,3,3a,4,6a-hexahydropentalen-1-ol (**S5e**) (9.1 mg, 0.02 mmol) was reacted and purified according to the general procedure to give the title compound (3.4 mg, 43% yield). Purity was established as the *exo* diastereomer by Method A: $t_R$ = 1.02 min, 89.2%.

**$^1$H NMR** (500 MHz, $CDCl_3$) δ 7.30 (d, *J* = 41.7 Hz, 5H), 7.17 (t, *J* = 7.8 Hz, 1H), 6.75 (t, *J* = 9.3 Hz, 2H), 6.69 (s, 1H), 5.07 (s, 1H), 5.02 (s, 1H), 4.88 (s, 1H), 3.94 (s, 1H), 2.35 (dd, *J* = 17.3, 8.0 Hz, 1H), 2.27 (d, *J* = 9.4 Hz, 1H), 2.05 (dt, *J* = 21.8, 6.9 Hz, 4H), 1.77 – 1.48 (m, 5H), 1.35 – 1.16 (m, 8H), 0.86 (t, *J* = 7.0 Hz, 3H).

**$^{13}$C NMR** (126 MHz, $CDCl_3$) δ 154.9, 154.6, 144.1, 141.4, 139.1, 138.6, 128.8, 127.72, 126.68, 122.4, 116.4, 115.1, 113.6, 82.1, 69.3, 55.8, 40.2, 34.0, 32.1, 31.7, 29.7, 29.4, 27.9, 27.8, 22.6, 14.1.

**LRMS** (ESI, APCI) *m/z*: calc'd for $C_{28}H_{37}O_2$ $[M+H]^+$ 403.3, found 403.8

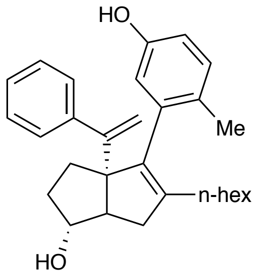


**(*exo*)-5-hexyl-4-(5-hydroxy-2-methylphenyl)-3a-(1-phenylvinyl)-1,2,3,3a,6,6a-hexahydropentalen-1-ol (18):** (*exo*)-4-(5-bromo-2-methylphenyl)-5-hexyl-3a-(1-phenylvinyl)-1,2,3,3a,6,6a-hexahydropentalen-1-ol (**S6e**) (10.2 mg, 0.02 mmol) was reacted and purified according to the general procedure to give the title compound (2.4 mg, 27% yield). Purity was established as the *exo* diastereomer by Method B: $t_R$ = 1.25 min, 97.6%.

**$^1$H NMR** (500 MHz, $CDCl_3$) δ 7.46 – 7.39 (m, 2H), 7.31 – 7.27 (m, 3H), 7.09 (d, *J* = 8.3 Hz, 1H), 6.67 (dd, *J* = 8.3, 2.8 Hz, 1H), 6.56 (d, *J* = 2.8 Hz, 1H), 5.20 (d, *J* = 1.1 Hz, 1H), 4.88 (d, *J* = 1.2

Hz, 1H), 4.52 (s, 1H), 3.99 (s, 1H), 3.49 (s, 3H), 2.64 (dd, $J$ = 17.3, 10.2 Hz, 1H), 2.33 (d, $J$ = 9.5 Hz, 1H), 2.10 – 1.99 (m, 3H), 1.94 – 1.77 (m, 3H), 1.71 (ddd, $J$ = 19.1, 13.3, 6.6 Hz, 2H), 1.42 – 1.12 (m, 7H), 0.85 (t, $J$ = 7.1 Hz, 3H).
**LRMS** (ESI, APCI) *m/z*: calc'd for $C_{29}H_{37}O_2$ [M+H] 417.3, found 416.9.

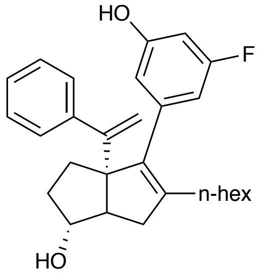


**(*exo)*-4-(3-fluoro-5-hydroxyphenyl)-5-hexyl-3a-(1-phenylvinyl)-1,2,3,3a,6,6a-hexahydropentalen-1-ol (20):** (*exo*)-4-(3-bromo-5-fluorophenyl)-5-hexyl-3a-(1-phenylvinyl)-1,2,3,3a,6,6a-hexahydropentalen-1-ol (**S7e**) (27.9 mg, 0.06 mmol) was reacted and purified according to the general procedure to give the title compound (5.9 mg, 24%). Purity was established as the *exo* diastereomer by Method B: $t_R$ = 1.14 min, 97.3%.
**$^1$H NMR** (500 MHz, $CDCl_3$) δ 7.35 – 7.28 (m, 2H), 7.26 (d, $J$ = 6.2 Hz, 3H), 6.55 – 6.45 (m, 3H), 5.09 (s, 1H), 5.04 (s, 1H), 4.94 (s, 1H), 3.93 (s, 1H), 2.35 (dd, $J$ = 17.0, 9.3 Hz, 1H), 2.28 (d, $J$ = 9.5 Hz, 1H), 2.15 – 1.99 (m, 4H), 1.73 – 1.64 (m, 3H), 1.38 – 1.18 (m, 6H), 0.90 – 0.83 (m, 3H).
**$^{13}$C NMR** (126 MHz, $CDCl_3$) δ 162.0, 156.2, 154.4, 143.9, 142.3, 137.8, 127.8, 127.7, 126.8, 115.3, 112.4, 109.2, 109.0, 101.7, 101.5, 82.0, 69.3, 55.8, 40.2, 34.0, 32.0, 31.6, 29.7, 29.4, 27.7, 22.6, 14.1.
**$^{19}$F NMR** (282 MHz, $cdcl_3$) δ -112.75.
**LRMS** (ESI, APCI) *m/z*: calc'd for $C_{28}H_{36}FO_2$ $[M+H]^+$ 421.3, found 421.9

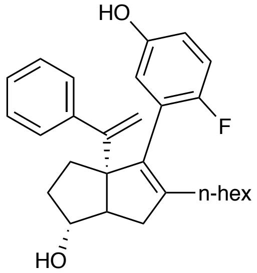


**(*exo*)-4-(2-fluoro-5-hydroxyphenyl)-5-hexyl-3a-(1-phenylvinyl)-1,2,3,3a,6,6a-hexahydropentalen-1-ol (22):** (*exo*)-4-(5-amino-2-fluorophenyl)-5-hexyl-3a-(1-phenylvinyl)-1,2,3,3a,6,6a-hexahydropentalen-1-ol (**S8e**) (29.8 mg, 0.06 mmol) was reacted and purified according to the general procedure to give the title compound (3.4 mg, 13% yield). Purity was established as the *exo* diastereomer by Method B: $t_R$ = 1.03 min, 96.9%.
**$^1$H NMR** (500 MHz, $CDCl_3$δ 7.39 – 7.35 (m, 2H), 7.31 – 7.27 (m, 3H), 6.93 (t, $J$ = 8.8 Hz, 1H), 6.70 (dt, $J$ = 8.6, 3.5 Hz, 1H), 6.65 (dd, $J$ = 5.6, 3.2 Hz, 1H), 5.10 (d, $J$ = 1.2 Hz, 1H), 4.93 (d, $J$ = 1.3 Hz, 1H), 4.61 (s, 1H), 3.97 (s, 1H), 2.52 (dd, $J$ = 17.3, 9.7 Hz, 1H), 2.30 (d, $J$ = 9.7 Hz, 1H),

2.12 – 2.03 (m, 2H), 1.95 (t, $J$ = 7.8 Hz, 2H), 1.87 – 1.78 (m, 1H), 1.73 (dd, $J$ = 12.3, 6.3 Hz, 1H), 1.68 (dd, $J$ = 13.1, 5.9 Hz, 1H), 1.45 – 1.27 (m, 2H), 1.29 – 1.16 (m, 6H), 0.86 (t, $J$ = 7.1 Hz, 3H).
**$^{13}$C NMR** (126 MHz, $CDCl_3$) δ 156.1, 154.5, 154.2, 150.6, 144.4, 143.6, 127.9, 127.4, 126.8, 117.7, 117.6, 115.9, 115.7, 115.5, 114.9, 114.9, 81.9, 69.6, 55.4, 40.6, 33.6, 33.1, 31.6, 30.0, 29.4, 27.2, 22.6, 14.1.
**$^{19}$F NMR** (282 MHz, $cdcl_3$) δ -124.47.
**LRMS** (ESI, APCI) *m/z*: calc'd for $C_{28}H_{36}FO_2$ $[M+H]^+$ 421.2, found 421.9

**General Procedure for Amination (17, 19, 21, 23)**

A solution of ᵗBuBrettPhos (0.04 equiv), sodium *tert* butoxide (3.0 equiv) in 1,4-dioxane was treated with **S5e** – **S8e** as a solution in 1,4-dioxane and ammonia (0.5M in dioxane, ca. 10 equiv) (tube A). In a separate reaction tube (B), a solution of ᵗBuBrettPhos precatalyst (0.04 equiv.) in 1,4-dioxane was prepared. The solution in tube B was transferred to tube A. The reaction mixture in tube A was heated in a closed reaction tube at 80 °C for 16 hours behind a blast shield (for larger reaction quantities, a pressure tube behind a blast shield is recommended). The mixture was cooled to room temperature, diluted with water and extracted with EtOAc three times. The combined organics were washed with water and brine, dried over $MgSO_4$, and concentrated. The crude oil was purified by silica gel chromatography in EtOAc/Hexanes eluent.

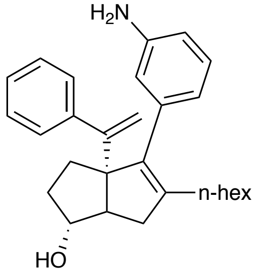


**(*exo*)-4-(3-aminophenyl)-5-hexyl-3a-(1-phenylvinyl)-1,2,3,3a,6,6a-hexahydropentalen-1-ol (17):** (*exo*)-6-(3-bromophenyl)-5-hexyl-3-(methoxymethoxy)-6a-(1-phenylvinyl)-1,2,3,3a,4,6a-hexahydropentalen-1-ol (**S5e**) (8.3 mg, 0.02 mmol) was reacted according to the general procedure. The crude oil was purified by silica gel chromatography in 20% EtOAc/Hexanes eluent (2.6 mg, 36% yield). Purity was established as the *exo* diastereomer by Method B: $t_R$ = 0.89 min, 90.1%.

**$^1$H NMR** (500 MHz, $CDCl_3$) δ 7.49 – 7.20 (m, 5H), 7.09 (dd, *J* = 9.6, 5.6 Hz, 1H), 6.61 (d, *J* = 7.4 Hz, 2H), 6.55 (s, 1H), 5.06 (d, *J* = 3.9 Hz, 1H), 5.02 (d, *J* = 4.2 Hz, 1H), 3.93 (s, 1H), 3.52 (s, 2H), 2.32 (dt, *J* = 13.6, 6.7 Hz, 1H), 2.25 (d, *J* = 8.8 Hz, 1H), 2.04 (ddd, *J* = 21.8, 11.9, 4.7 Hz, 5H), 1.77 – 1.62 (m, 3H), 1.38 – 1.06 (m, 8H), 0.86 (t, *J* = 7.1 Hz, 3H).

**$^{13}$C NMR** (126 MHz, $CDCl_3$) δ 154.7, 145.6, 144.2, 141.0, 139.0, 138.5, 128.5, 127.8, 127.7, 126.6, 120.5, 116.4, 114.9, 113.6, 110.0, 82.1, 55.9, 40.2, 34.1, 32.0, 31.7, 29.8, 29.4, 27.8, 22.6, 14.1.

**LRMS** (ESI, APCI) *m/z*: calc'd for $C_{28}H_{38}NO$ $[M+H]^+$ 402.3, found 401.9

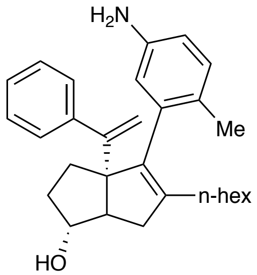


**(*exo*)-4-(5-amino-2-methylphenyl)-5-hexyl-3a-(1-phenylvinyl)-1,2,3,3a,6,6a-hexahydropentalen-1-ol (19):** (*exo*)-4-(5-bromo-2-methylphenyl)-5-hexyl-3a-(1-phenylvinyl)-

1,2,3,3a,6,6a-hexahydropentalen-1-ol (**S6e**) (10.2 mg, 0.02 mmol) was reacted according to the general procedure. The crude oil was purified by silica gel chromatography in 20-30% EtOAc/Hexanes eluent (1.2 mg, 14% yield). Purity was established as the *exo* diastereomer by Method B: $t_R$ = 0.53 min, 97.5%.

**$^1$H NMR** (500 MHz, $CDCl_3$) δ 7.44 – 7.38 (m, 2H), 7.33 – 7.27 (m, 2H), 7.02 (d, *J* = 8.0 Hz, 1H), 6.85 (dd, *J* = 8.9, 5.0 Hz, 1H), 6.54 (dd, *J* = 8.0, 2.5 Hz, 1H), 6.47 (d, *J* = 2.5 Hz, 1H), 5.18 (d, *J* = 1.1 Hz, 1H), 4.89 (d, *J* = 1.2 Hz, 1H), 3.98 (s, 1H), 3.79 (s, 2H), 3.53 (s, 3H), 2.61 (dd, *J* = 17.2, 10.0 Hz, 1H), 2.30 (d, *J* = 9.7 Hz, 1H), 2.06 – 1.99 (m, 2H), 1.97 – 1.79 (m, 2H), 1.77 – 1.68 (m, 2H), 1.38 – 1.07 (m, 11H), 0.86 (t, *J* = 7.1 Hz, 3H).

**LRMS** (ESI, APCI) *m/z*: calc'd for $C_{29}H_{38}NO$ [M+H] 416.3, found 415.9.

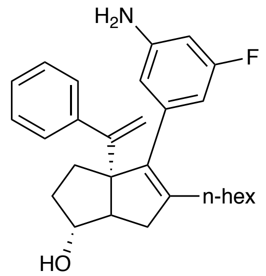


**(*exo*)-4-(3-amino-5-fluorophenyl)-5-hexyl-3a-(1-phenylvinyl)-1,2,3,3a,6,6a-hexahydropentalen-1-ol (21):** (*exo*)-4-(3-bromo-5-fluorophenyl)-5-hexyl-3a-(1-phenylvinyl)-1,2,3,3a,6,6a-hexahydropentalen-1-ol (**S7e**) (44.5 mg, 0.09 mmol) was reacted according to the general procedure. The crude oil was purified by silica gel chromatography in 20-30% EtOAc/Hexanes eluent (3.2 mg, 8% yield). Purity was established as the *exo* diastereomer by method B: $t_R$ = 1.16 min, 95.9%.

**$^1$H NMR** (500 MHz, $CDCl_3$) δ 7.36 – 7.30 (m, 2H), 7.31 – 7.20 (m, 3H), 6.41 – 6.21 (m, 3H), 5.08 (d, *J* = 1.6 Hz, 1H), 5.05 (d, *J* = 1.6 Hz, 1H), 3.92 (s, 1H), 3.79 (d, *J* = 1.7 Hz, 1H), 3.70 (s, 2H), 3.53 (d, *J* = 1.6 Hz, 1H), 2.32 (dd, *J* = 16.7, 9.3 Hz, 1H), 2.25 (d, *J* = 9.4 Hz, 1H), 2.13 – 1.97 (m, 4H), 1.79 – 1.63 (m, 3H), 1.42 – 1.12 (m, 6H), 0.87 (t, *J* = 7.1, 6.5 Hz, 3H).

**$^{13}$C NMR** (126 MHz, $CDCl_3$) δ 164.2, 162.3, 154.5, 147.2, 147.1, 144.0, 141.8, 140.2, 140.1, 138.2, 127.7, 126.7, 119.7, 115.1, 112.1, 106.9, 106.7, 100.7, 100.5, 82.0, 69.2, 55.9, 40.2, 34.1, 32.0, 31.7, 31.1, 29.7, 29.4, 28.2, 27.7, 25.4, 23.9, 23.5, 22.6, 14.1.

**$^{19}$F NMR** (282 MHz, $CDCl_3$) δ -114.23.

**LRMS** (ESI, APCI) *m/z*: calc'd for $C_{28}H_{37}FNO$ $[M+H]^+$ 420.3, found 420.8

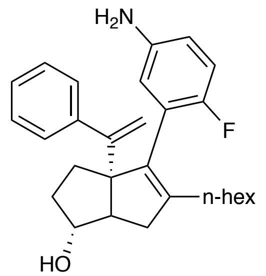


**(*exo*)-4-(5-amino-2-fluorophenyl)-5-hexyl-3a-(1-phenylvinyl)-1,2,3,3a,6,6a-hexahydropentalen-1-ol (23):** (*exo*)-4-(5-amino-2-fluorophenyl)-5-hexyl-3a-(1-phenylvinyl)-1,2,3,3a,6,6a-hexahydropentalen-1-ol (**S8e**) (33.5 mg, 0.07 mmol) was reacted according to the general procedure. The crude oil was purified by silica gel chromatography in 30-50%

EtOAc/Hexanes eluent (5.2 mg, 18 yield%). Purity was established as the *exo* diastereomer by Method B: $t_R$ = 0.81 min, 98.5%.
**$^1$H NMR** (500 MHz, $CDCl_3$) δ 7.38 (dd, *J* = 6.7, 2.9 Hz, 2H), 7.32 – 7.25 (m, 2H), 6.99 – 6.79 (m, 2H), 6.56 (dt, *J* = 8.6, 3.5 Hz, 1H), 6.50 (dd, *J* = 6.0, 2.9 Hz, 1H), 5.09 (d, *J* = 1.3 Hz, 1H), 4.93 (d, *J* = 1.3 Hz, 1H), 3.96 (d, *J* = 3.6 Hz, 1H), 3.79 (s, 1H), 3.53 (s, 1H), 3.46 (s, 2H), 2.56 – 2.44 (m, 1H), 2.28 (d, *J* = 9.7 Hz, 1H), 2.10 – 2.02 (m, 2H), 1.96 (t, *J* = 7.8 Hz, 2H), 1.88 – 1.78 (m, 1H), 1.75 (dd, *J* = 12.0, 6.4 Hz, 1H), 1.67 (dd, *J* = 12.9, 5.8 Hz, 1H), 1.35 (q, *J* = 6.9 Hz, 1H), 1.32 – 1.15 (m, 6H), 0.86 (t, *J* = 7.0 Hz, 3H).
**$^{13}$C NMR** (126 MHz, $CDCl_3$) δ 154.7, 144.6, 135.5, 129.6, 127.9, 127.6, 127.5, 126.7, 120.1, 119.7, 117.7, 115.6, 115.3, 114.9, 82.0, 69.6, 55.4, 40.5, 33.6, 33.0, 31.7, 31.1, 30.0, 29.4, 28.2, 27.3, 26.8, 25.4, 23.9, 23.5, 22.6, 14.1.
**$^{19}$F NMR** (282 MHz, $CDCl_3$) δ -126.95
**LRMS** (ESI, APCI) *m/z*: calc'd for $C_{28}H_{37}FNO$ $[M+H]^+$ 420.3, found 419.9

### d. Spectral Data for Key Compounds

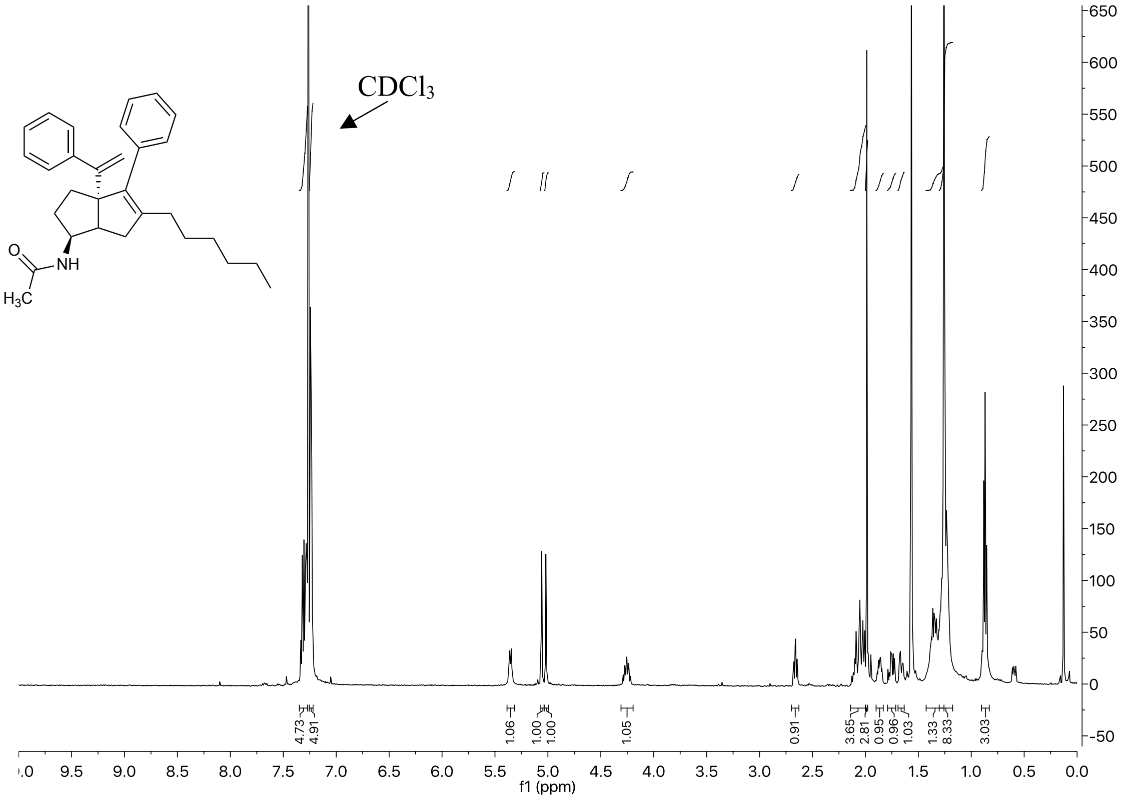


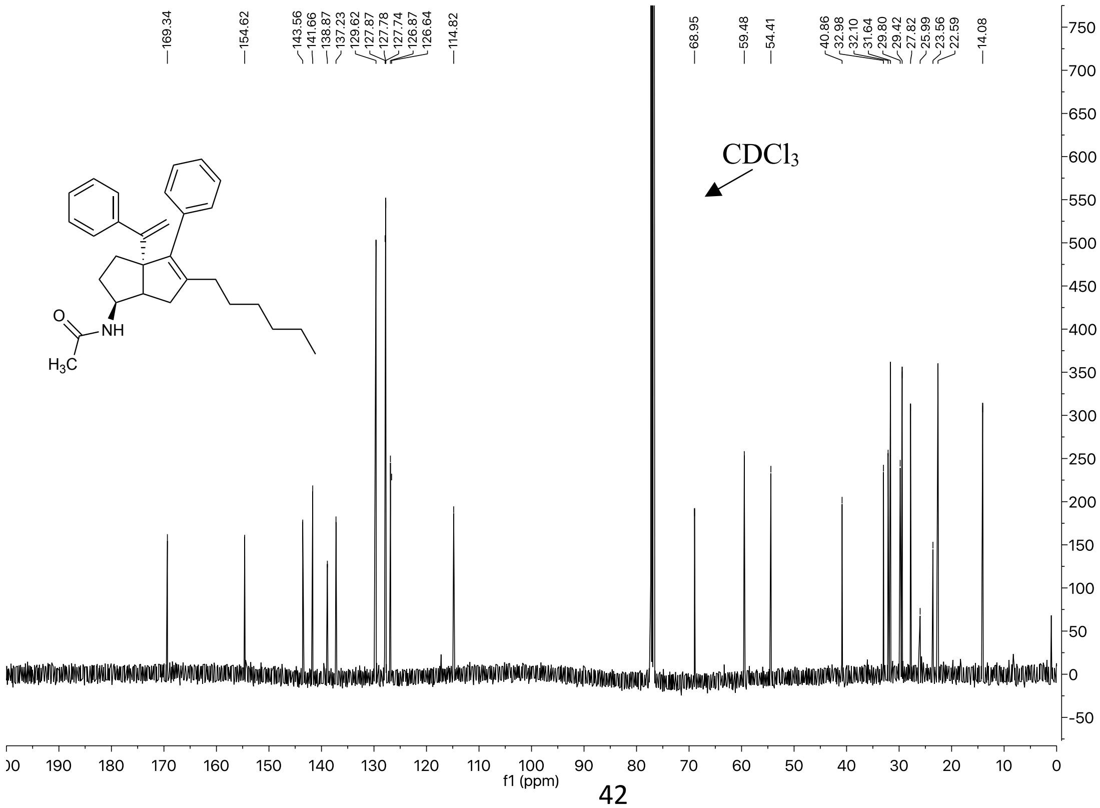

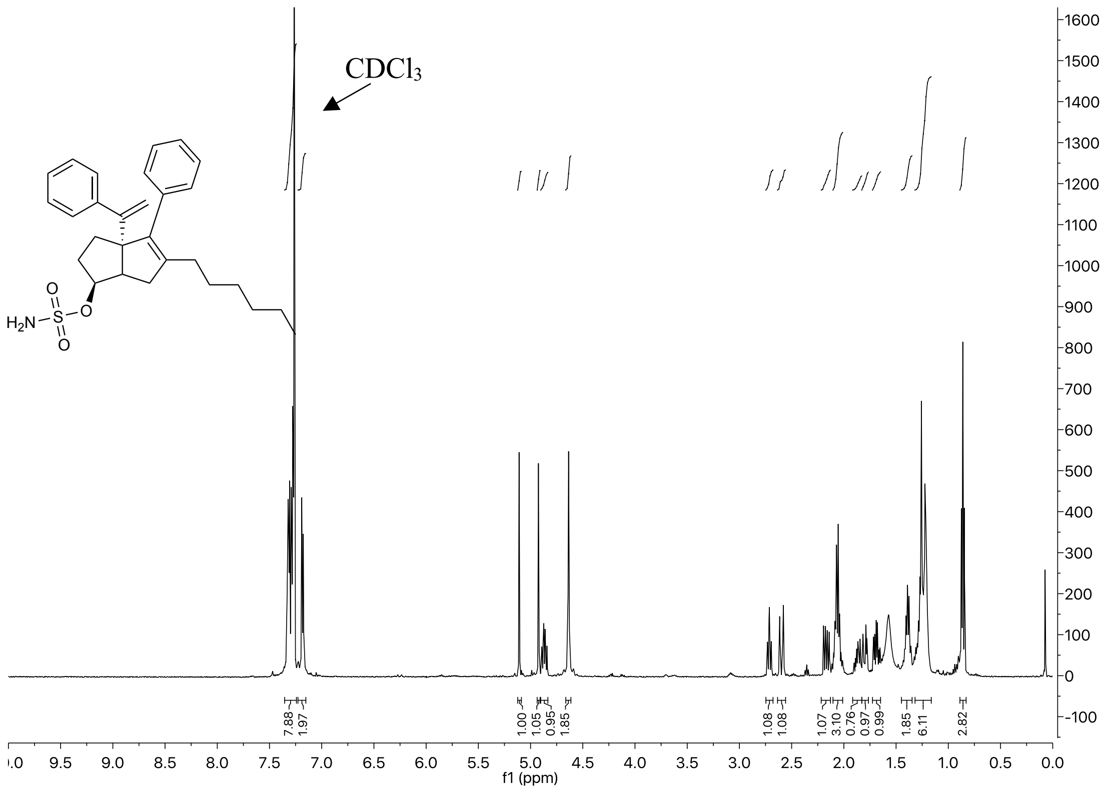
CDCl3
H2N
O
S
O
f1 (ppm)

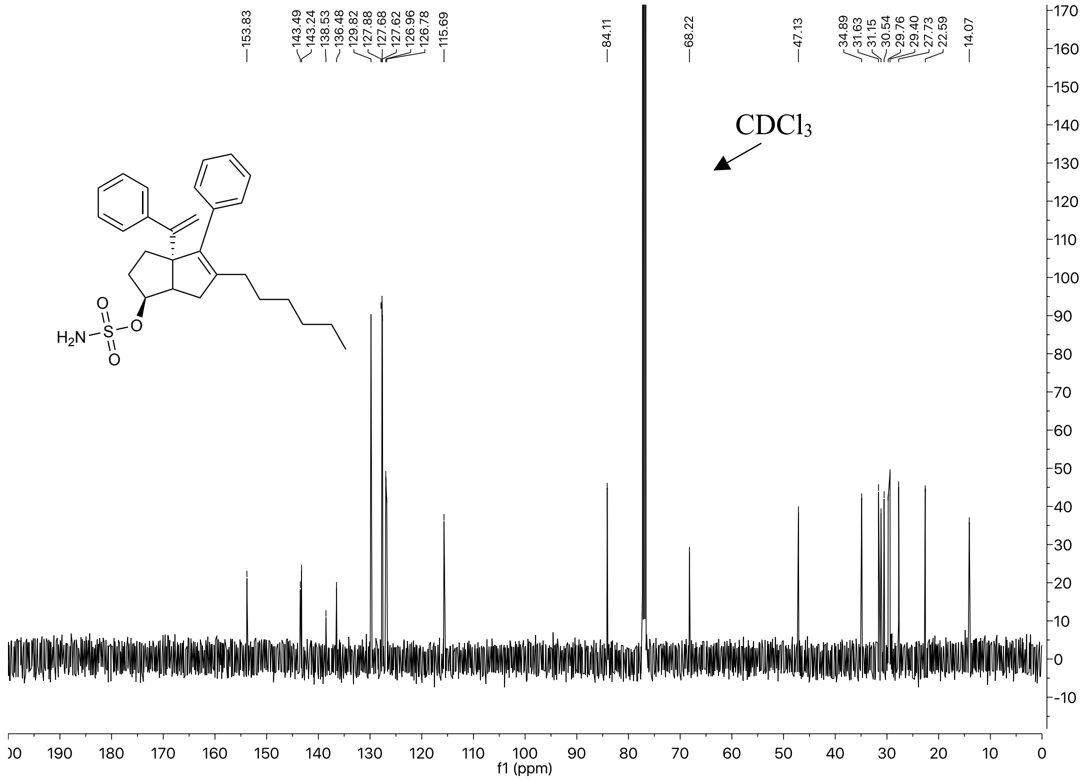
CDCl3
H2N
O
S
O
f1 (ppm)

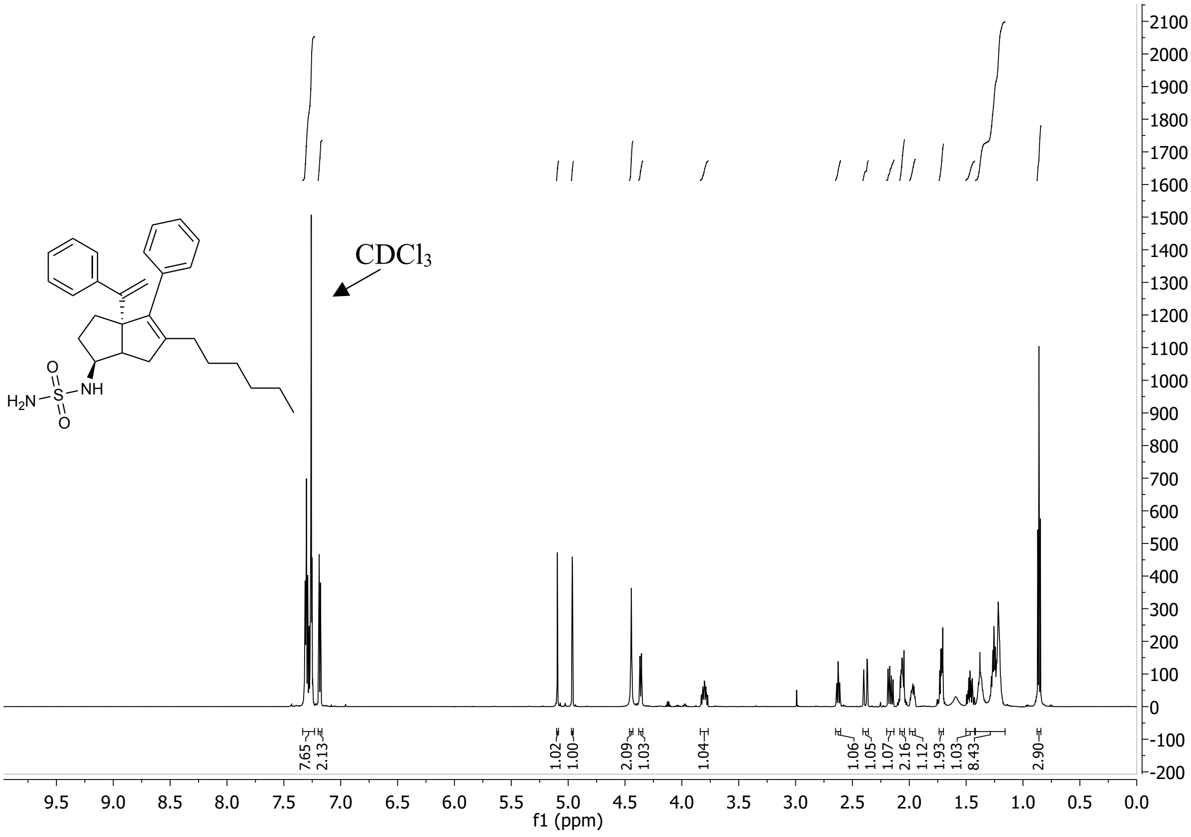
CDCl3
H2N
S
O
O
NH
7.65
2.13
1.02
1.00
2.09
1.03
1.04
1.06
1.05
1.07
2.16
1.12
1.93
1.03
8.43
2.90
f1 (ppm)

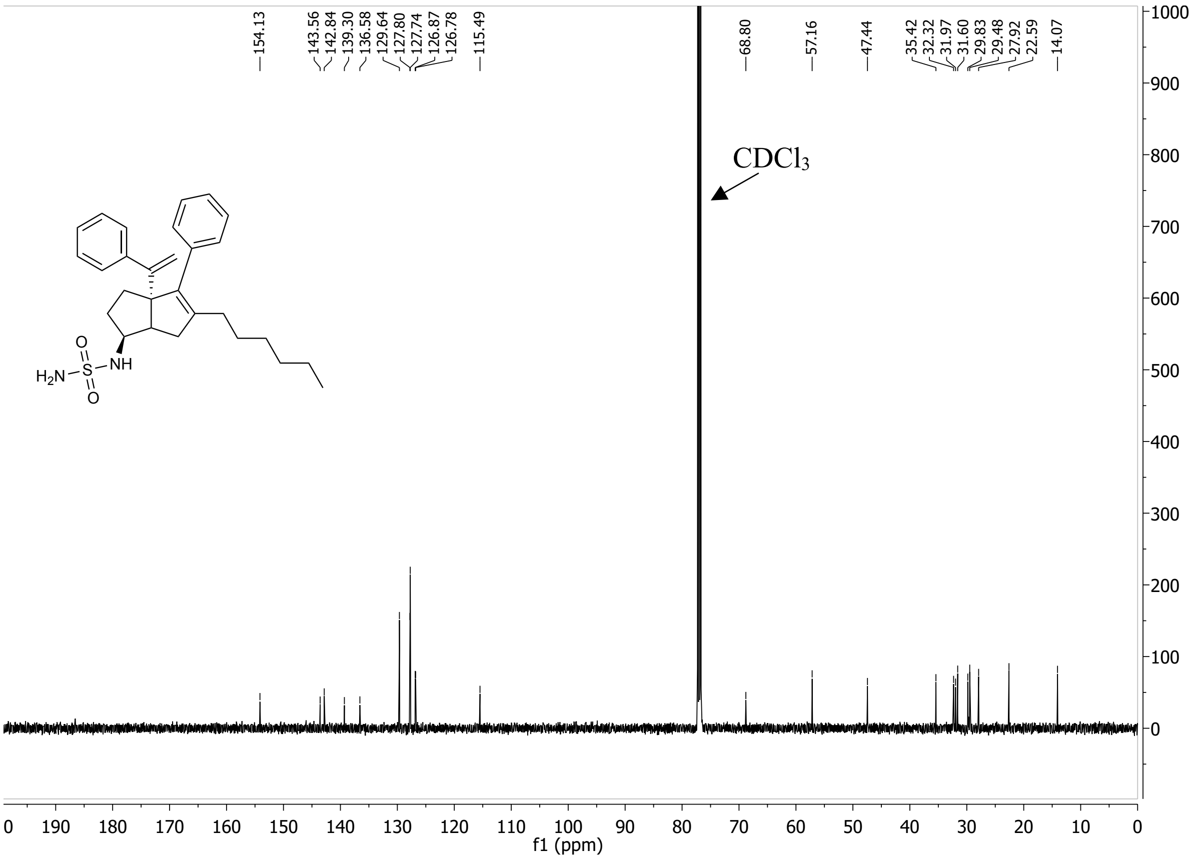
154.13
143.56
142.84
139.30
136.58
129.64
127.80
127.74
126.87
126.78
115.49
68.80
57.16
47.44
35.42
32.32
31.97
31.60
29.83
29.48
27.92
22.59
14.07
CDCl3
H2N
S
O
O
NH
f1 (ppm)
